# Golgi-associated nuclear import receptor importin-7 targets HPV from the Golgi to the nucleus to promote infection

**DOI:** 10.1101/2025.01.08.631933

**Authors:** Tai-Ting Woo, Yuka Takeo, Mara C. Harwood, Ethan T. Houck, Daniel DiMaio, Billy Tsai

## Abstract

During human papillomavirus (HPV) entry, the virus exploits COPI-dependent retrograde transport to cross the trans-Golgi network and Golgi stacks before reaching the nucleus to cause infection. How HPV enters the nucleus after exiting the Golgi is unclear, although mitotic nuclear envelop breakdown (NEB) appears important. Here we show that importin-7 (IPO7), a nuclear pore import receptor, associates with the Golgi and promotes HPV infection. Upon IPO7 knockdown, HPV infection is inhibited and the virus accumulates in the Golgi but does not enter the nucleus, demonstrating that IPO7 promotes Golgi-to-nucleus transport of HPV. We further reveal that the C-terminus of HPV capsid protein L2, which is thought to contain overlapping nuclear localization and cell-penetrating peptide sequences, binds directly to IPO7 in a step that requires COPI-dependent virus trafficking. Together these data identify a role of an importin in HPV infection, raising the possibility that the canonical nuclear pore import machinery plays an unanticipated role in NEB-dependent nuclear entry.

**Teaser:** HPV, a cancer-causing human pathogen, exploits the Golgi-associated nuclear import receptor IPO7 for Golgi-to-nucleus trafficking to promote infection.

## Introduction

The human papillomavirus (HPV) causes approximately 5% of all human cancers globally, including essentially all cervical cancer as well as a significant portion of other anogenital and non-genital cancers (*1, 2*). Although effective vaccines against HPV infection exist, vaccination uptake remains persistently low. Additionally, there are no specific anti-virals against active HPV infection. Hence, a better understanding of the mechanism of HPV infection may lead to the development of effective therapies against HPV disease.

Structurally, HPV is a small non-enveloped DNA virus composed of 72 pentamers of the major capsid protein L1 and up to 72 copies of the minor capsid protein L2. The L1 and L2 proteins encapsidate the viral DNA genome which must be delivered to the nucleus of the host cell in order to cause infection (*3*). HPV initiates entry through L1 binding to heparin sulfate proteoglycans on the host cell surface or in the extracellular matrix. This interaction imparts conformational changes to the capsid and subsequent furin-mediated cleavage of the N-terminus of L2 (*4–6*). Via an unidentified entry receptor, the virus is endocytosed and reaches the endosome (*7–9*). Here the acidic luminal environment triggers partial disassembly of the capsids, exposing the C-terminus of L2 which in turn engages the endosome-localized membrane protein γ-secretase (*10, 11*). γ-secretase deploys an unconventional chaperone activity to facilitate membrane penetration of L2 (*10*), a step that requires the cell-penetrating peptide (CPP) of L2 which is located near its C-terminus (*12, 13*). Membrane penetration effectively inserts most of L2 across the endosome membrane, exposing a large portion of the L2 capsid protein to the cytoplasm (*14, 15*).

Importantly, in this membrane-inserted topology, L2 recruits different cytosolic host factors that direct proper virus trafficking. For instance, the retromer sorting machinery, along with the dynein-BICD2 motor complex, interacts with L2 to enable trafficking of HPV from the endosome to the *trans*-Golgi network (TGN) (*16–20*). Upon arrival to the TGN, L2 further binds to the COPI sorting complex, which transports HPV across the Golgi stack membranes in a retrograde manner prior to reaching the nucleus to cause infection (*21*).

How HPV reaches the nucleus after it exits the Golgi and the role of L2 in this step are largely unknown. Nuclear envelop breakdown (NEB) during mitosis has been hypothesized as a mechanism by which HPV enters the nucleus (*22–24*). Nuclear import of many cellular proteins involves importins, which typically bind to and transport cellular cargos containing a nuclear localization signal (NLS) from the cytosol to the nucleoplasm via the nuclear pores. After nuclear entry, binding of Ran-GTP to the importin-cargo complex triggers the release of the cargo from the importins (*25*). The Ran-binding protein 10 (RanBP10) and karyopherin alpha2 (KPNA2), an importin α protein, have been implicated in nuclear entry of HPV during infection (*26*). In addition, other components of the classic nuclear import machinery including importin α proteins and nuclear import receptors are involved in nuclear import of the L2 protein expressed in the absence of L1, but these interactions may be important for nuclear import of newly synthesized L2 prior to capsid assembly during the late stages of the viral life cycle, rather than participating in nuclear targeting during virus entry (*27–30*). How (or whether) the incoming HPV exploits this nuclear import machinery to gain nuclear entry remains elusive.

In this study, by using an artificial protein that inhibits HPV infection, unbiased proteomics, pharmacological inhibition, gene-targeting knockdown (KD), as well as biochemical and microscopy approaches, we showed that the importin β family member importin-7 (IPO7) is a Golgi-associated host factor that plays a critical role in HPV infection. Specifically, we found that KD of IPO7 causes the virus to accumulate in the Golgi, thereby preventing HPV from reaching the nucleus and inhibiting infection. These results indicate that this importin family member plays a key role in Golgi-to-nuclear transport of HPV. We further show that IPO7 binds directly to a C-terminal region of L2 that harbors overlapping nuclear localization and CPP sequences. Furthermore, the IPO7-HPV L2 interaction occurs after COPI-dependent retrograde trafficking during virus entry, and COPI depletion blocks binding between IPO7 and L2. Our study suggests that HPV exploits the nuclear pore import machinery in a novel way prior to NEB-dependent nuclear entry.

## Results

### Study of a Golgi-localized traptamer reveals an important role of importin β members in HPV infection

We previously used a protein modulation screen to isolate four artificial FLAG-APEX2-tagged (FA) transmembrane proteins (called traptamers) that block different HPV entry steps (*18, 31*). These traptamers are thought to bind to and compromise the activity of host factors required for HPV infection. When the FA-JX4 traptamer was stably expressed in HeLa cells, it localizes to the TGN and Golgi apparatus (Fig. S1A) (*18*) and impairs HPV infection by trapping HPV in these compartments, thereby preventing arrival of the packaged DNA in the nucleus (*18*). We hypothesized that FA-JX4 binds to and inhibits critical Golgi-localized host factors essential for HPV trafficking. To identify host proteins that bind FA-JX4 and participate in the late stages of entry, we infected cells expressing FA-JX4 or the control FA (a non-inhibitory traptamer lacking a transmembrane domain) and performed immunoprecipitation (IP) followed by mass spectrometry (MS) analysis. For these experiments, we used HPV16 pseudovirus (PsV) that is composed of the high-risk HPV16 L1 and L2 capsid proteins (without epitope tags) encapsidating a reporter plasmid encoding the green fluorescent protein (GFP) in place of the HPV16 viral genome. In this system, expression of GFP indicates successful entry of the PsV DNA to the nucleus, thereby allowing expression of the reporter plasmid. Notably, HPV16 PsV infection recapitulates infection by authentic HPV isolated from the stratified keratinocyte raft cultures and is widely used in studies of HPV entry (*32*).

At 24 hours post infection (hpi), cell extracts were collected and subjected to IP using an anti-FLAG antibody to pull down FA or FA-JX4, and the co-precipitated material was analyzed by MS. A fraction of the precipitated material was analyzed by SDS-PAGE followed by immunoblotting to confirm that similar levels of FA and FA-JX4 proteins were IPed (Fig. 1A). The MS results showed that neither HPV16 L1 or L2 capsid protein was present in the FA or FA-JX4 IPed sample (Fig. 1B; the complete data are available in Table S1), suggesting that FA-JX4 does not directly interact with the virus to block its entry. Instead, several Golgi-associated COPI subunits were found in the FA-JX4 (but not FA) IPed sample (Fig. 1B), consistent with the observation that FA-JX4 localizes to the TGN/Golgi (Fig. S1A) (*18*). Importantly, the FA-JX4 (but not FA) IPed sample contained multiple peptides from five members of the importin β family, namely IPO1, IPO4, IPO5, IPO7, and IPO8, as well as a single peptide from IPO9 (Fig. 1B). Importin β family proteins are components of the nuclear import machinery that transports cargos from the cytosol to the nucleoplasm via nuclear pores. However, importin β family proteins are not known to be associated with the Golgi and their involvement in HPV infection has not been examined.

**Figure 1.**
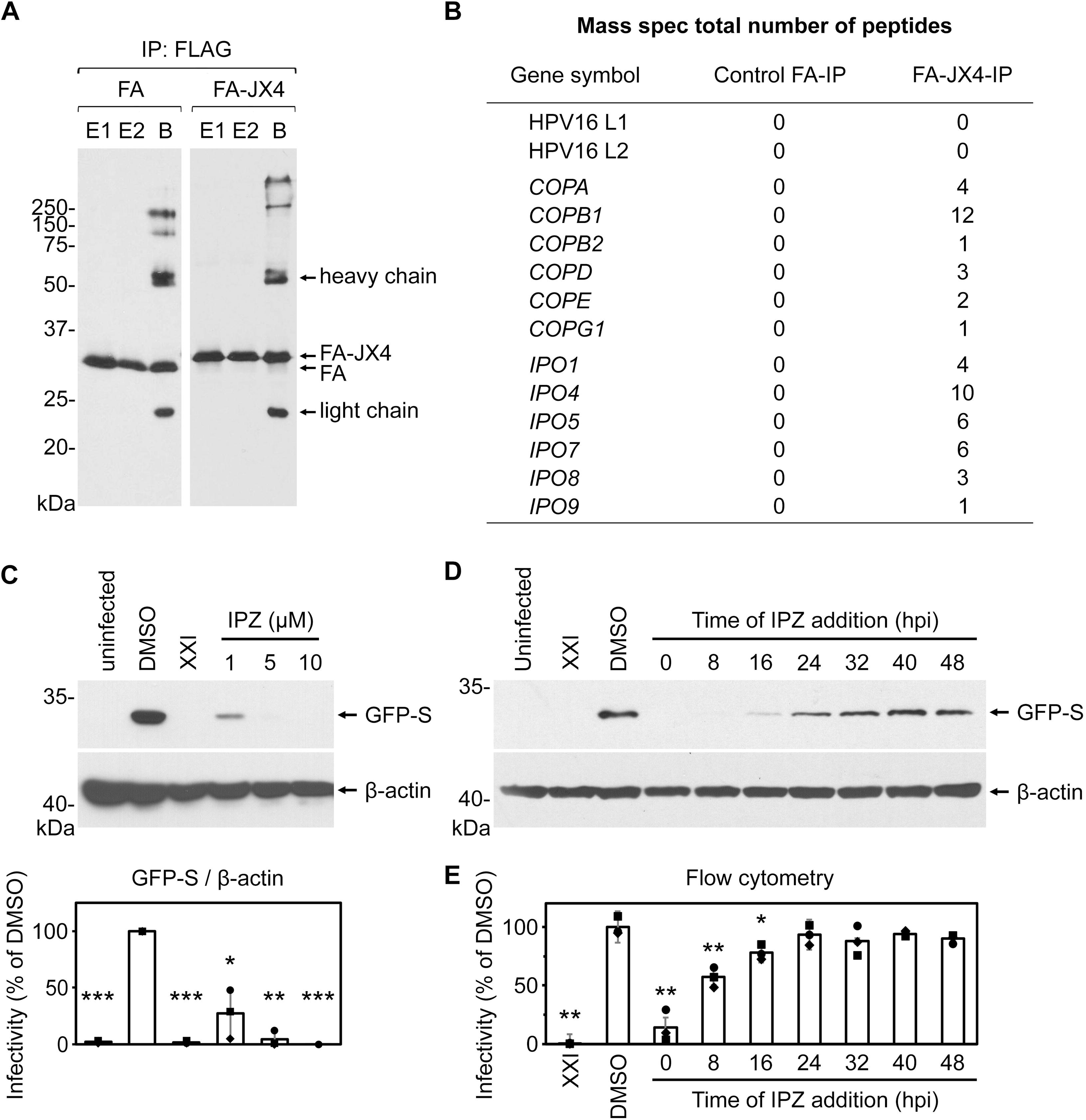
Study of a Golgi-localized traptamer reveals an important role of importin β members in HPV infection. **(A)** HeLa S3 cells stably expressing traptamer FA-JX4 or the control FA were infected with untagged HPV16 PsV (∼25 μg L1). At 24 hours post infection (hpi), cells were lysed, and the resulting extracts were subjected to immunoprecipitation using an anti-FLAG antibody followed by incubation with protein G beads. The precipitated materials were eluted using 3xFLAG peptides twice (E1, eluate 1; E2, eluate 2). The remaining materials on the beads (B, beads) were further eluted using SDS sample buffer. A portion of each eluate was analyzed by immunoblotting using an anti-FLAG antibody to evaluate the levels of FA or FA-JX4 proteins that were precipitated. E1 and E2 were combined and subjected to mass spectrometry analyses. **(B)** Total number of peptides corresponding to subunits of the COPI complex and importin β family proteins identified in the eluted samples (E1 and E2) as in panel (A) by mass spectrometry. The entire mass spectrometry data are available in Table S1. **(C)** The importin β inhibitor blocks HPV infection. HeLa cells were uninfected or infected with HPV16.L2F PsV (MOI ∼0.3) containing a GFP reporter plasmid. Importazole (IPZ) at the indicated concentration (1–10 µM), 1 µM γ-secretase inhibitor (XXI), or the solvent dimethyl sulfoxide (DMSO) was added to cells at the time of infection. At 48 hpi, cells were lysed and the resulting whole cell extract was subjected to SDS-PAGE followed by immunoblotting with antibodies recognizing GFP or β-actin as a loading control (representative immunoblots shown in upper panels). The results were quantified (lower panel), in which the intensity of the GFP-S band was normalized to that of β-actin in each sample. The infectivity in cells treated with DMSO was used as the normalization standard. The means and standard deviations of three independent experiments with individual data points are shown. A two-tailed, unequal variance *t*-test was used to determine statistical significance compared to DMSO treated cells infected with HPV16.L2F PsV. \**P* < 0.05; \*\**P* < 0.01; \*\*\**P* < 0.001. **(D)** Immunoblotting for GFP-S expression was performed as in (C), except cultures were treated with 10 µM IPZ at different time points after infection, as indicated. **(E)** As in (D), except flow cytometry was used to determine fraction of GFP-expressing cells. The results were normalized to the infected fraction of cells treated with DMSO. The individual data points, means, and standard deviations are shown (n = 3). A two-tailed, unequal variance *t*-test was used to determine statistical significance compared to DMSO-treated cells. \**P* < 0.05; \*\**P* < 0.01.

To determine if importin β proteins are important in HPV infection, we first tested whether HPV infection was perturbed by importazole (IPZ), an importin β inhibitor (*33*). In these experiments, we used HPV16 PsV containing HPV16 L1, 3xFLAG-tagged HPV16 L2 (to detect L2), and a reporter plasmid encoding S-tagged GFP (GFP-S). Following infection with this PsV, referred to as HPV16.L2F PsV, infectivity can be monitored by the expression of GFP-S. Compared to HeLa cells treated with the control dimethyl sulfoxide (DMSO), addition of IPZ at the same time as HPV16.L2F PsV reduced GFP-S expression in a concentration-dependent manner as assessed by immunoblotting (Fig. 1C; quantified in graph below). As a control, the γ-secretase inhibitor XXI blocked HPV infection, as expected (*34*). IPZ also inhibited HPV infection in the human cervical squamous carcinoma SiHa cells (Fig. S1B) and in human skin HaCaT keratinocytes (Fig. S1C). These findings suggest that importin β proteins promote HPV infection in multiple cell types.

Because IPZ treatment for as short as one hour can inhibit nuclear import events (*33*), we conducted a time-course experiment to assess when the importin β proteins act during HPV entry. In HeLa cells, IPZ was added either at the same time as HPV16.L2F PsV (0 hpi), or at eight-hour intervals after virus addition, and the expression of GFP-S was analyzed by immunoblotting. IPZ maintained its ability to reduce HPV infection even if the drug was added 16 hpi, but lost most of its inhibitory effect when added 24 hpi (Fig. 1D). Similar data were obtained when flow cytometry for GFP fluorescence was used as an alternative method to detect GFP-S expression (Fig. 1E). Because nuclear entry events associated with HPV occurs approximately 24 hpi (*23, 35*), these findings are consistent with the hypothesis that importin β family proteins play a pivotal role during relatively late stages of HPV entry.

### IPO7 promotes HPV infection

We next used a small interfering RNA (siRNA)-mediated KD approach to further support a role of the importin β family proteins in HPV infection. We tested two different siRNAs against each of the six FA-JX4-interacting importin β proteins (Fig. 1B) to deplete the intended target in HeLa cells. Compared to cells transfected with the control scrambled siRNA (Scr), all of the siRNAs except *IPO4* siRNA #1 robustly reduced the level of their intended protein (Fig. S2A). The KD cells were infected with HPV16.L2F PsV, and infectivity was assessed by flow cytometry for GFP fluorescence. Although the different siRNAs decreased infectivity by varying extents compared to control cells, KD of IPO5 or IPO7 caused the most severe inhibitory effect (Fig. 2A). *IPO7* siRNA #2 caused the strongest block of HPV infection, to an extent similar to cells depleted of the γ-secretase subunit PS1 (Fig. 2A) (*10, 34*). Immunoblotting for GFP-S after KD of IPO7 (using siRNA #2) also demonstrated a marked inhibition of infection (Fig. S2B). KD of IPO7 did not affect the distribution of cells in the cell cycle compared to control cells (Fig. S2C), suggesting that the inhibition of HPV infection by IPO7 KD is not caused by a defect in cell cycle progression. By contrast, IPO5 KD altered the cell cycle profile (Fig. S2D) and therefore may elicit unintended effects on HPV entry. Because depletion of IPO7 led to the most pronounced impairment of HPV infection without affecting cell cycle progression, for the remainder of this study we focused on elucidating the role of IPO7 in HPV infection.

**Figure 2.**
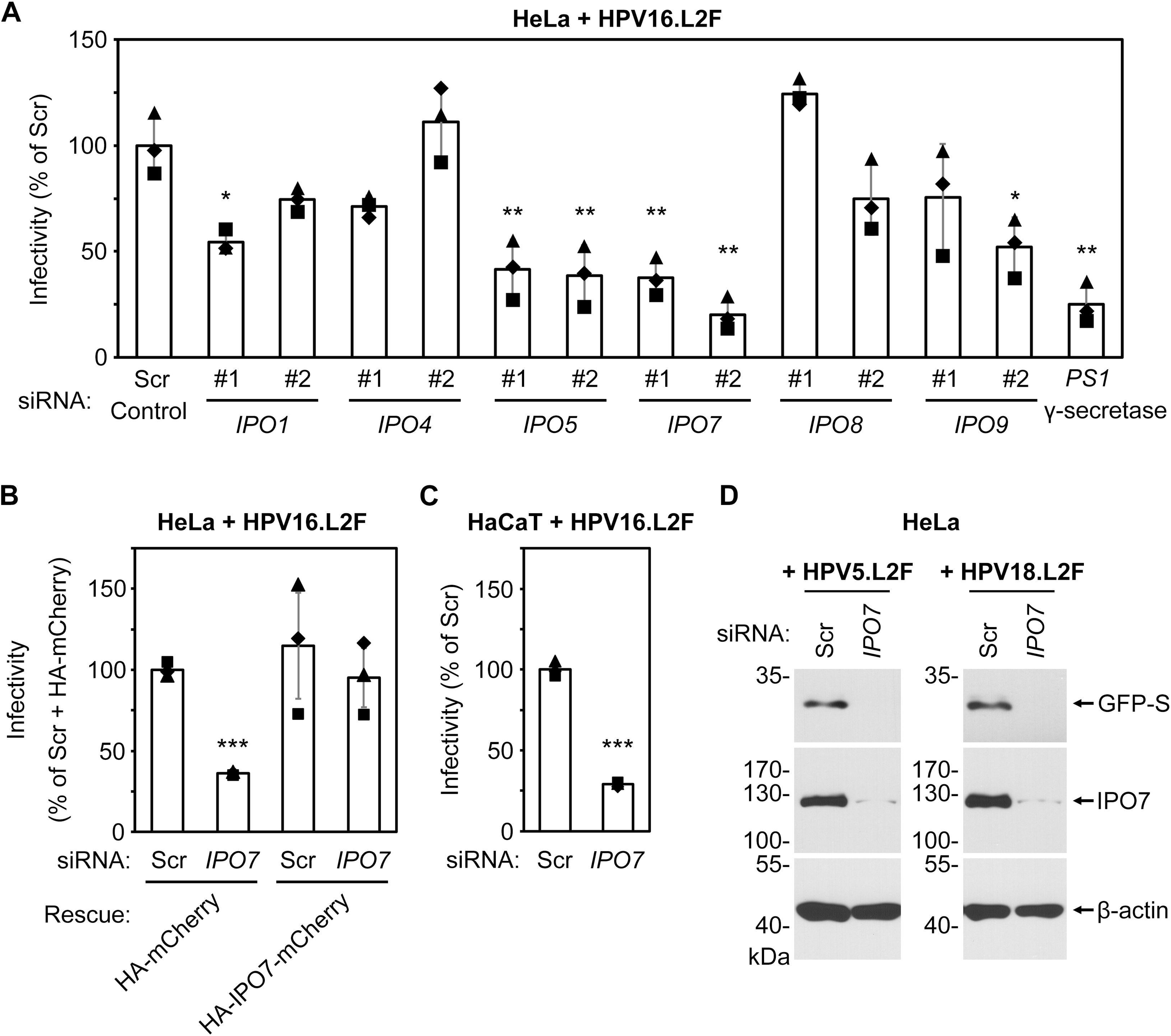
IPO7 promotes HPV infection. **(A)** HeLa cells transfected with 10 nM of the indicated siRNA for 48 h were infected at MOI ∼0.3 with HPV16.L2F PsV containing a GFP reporter plasmid. At 48 hpi, flow cytometry was used to determine the fraction of GFP-expressing cells. The results were normalized to the infected fraction of cells treated with scrambled (Scr) siRNA. The individual data points, means, and standard deviations are shown (n = 3). A two-tailed, unequal variance *t*-test was used to determine statistical significance when compared to scrambled siRNA-treated cells. \**P* < 0.05; \*\**P* < 0.01. **(B)** HeLa cells were transfected with indicated DNA constructs (rescue) for 24 h to express siRNA-resistant HA-IPO7-mCherry or the control HA-mCherry, followed by another transfection with Scr or *IPO7* siRNA #2 for 48 h. The cells were then infected at MOI ∼0.3 with HPV16.L2F PsV containing a GFP reporter plasmid. At 48 hpi, flow cytometry was used to measure GFP and mCherry fluorescence. The fraction of cells expressing GFP in the mCherry-positive population is graphed. The results were normalized to the number of infected cells in cells co-transfected with Scr siRNA and HA-mCherry-expressing plasmid. Data from three independent experiments were analyzed and presented as in (A), compared to cells transfected with Scr siRNA and control HA-mCherry plasmid. \*\*\**P* < 0.001. **(C)** As in (A), except HaCaT cells were transfected with 10 nM Scr or *IPO7* siRNA #2. \*\*\**P* < 0.001. **(D)** HeLa cells were transfected with 10 nM Scr or *IPO7* siRNA #2 for 48 h, followed by infection with HPV5.L2F or HPV18.L2F PsV containing a GFP reporter plasmid. At 48 hpi, whole cell extracts were subjected to SDS-PAGE followed by immunoblotting with antibodies recognizing GFP, IPO7, or β-actin.

To establish that the IPO7 KD phenotype in HPV infection is not due to an off-target effect, we performed a KD-rescue experiment. Control or IPO7 KD cells were transfected with a plasmid expressing hemagglutinin (HA)-tagged mCherry (HA-mCherry) or HA-mCherry fused to (siRNA-resistant) IPO7 (HA-IPO7-mCherry) (Fig. S2E). Cells were then infected with HPV16.L2F PsV, and flow cytometry was performed two days after infection. GFP fluorescence was used as a measure of infectivity and mCherry fluorescence was used as a measure of expression of the rescue construct. We found that HPV infection in cells expressing the control HA-mCherry was inhibited by IPO7 KD (Fig. 2B; compare second to first bar), as expected (Fig. 2A). In contrast, expression of HA-IPO7-mCherry almost entirely rescued the infectivity defect caused by IPO7 KD (Fig. 2B; compare fourth to second bar); expression of HA-IPO7-mCherry in control cells did not enhance infectivity (Fig. 2B; compare third to first bar). Restoration of HPV infection in IPO7 KD cells by expression of HA-IPO7-mCherry rules out off-target effects, establishing that IPO7 promotes HPV infection.

As assessed by flow cytometry (Fig. 2C) and immunoblotting (Fig. S2F), IPO7 plays an important role in HPV infection in HaCaT cells as well as in HeLa cells. Additionally, IPO7 also displays a crucial function in HPV5 and HPV18 PsV infection in HeLa cells (Fig. 2D). Together, these results reveal a requirement of IPO7 in HPV infection in several cell lines and HPV types.

### IPO7 associates with the Golgi-membrane

Because FA-JX4 localizes to the TGN/Golgi membrane (Fig. S1A) (*18*), we asked if IPO7 also localizes to the Golgi by conducting a cell fractionation experiment followed by immuno-isolation of the Golgi membrane. Extracts of uninfected HeLa cells were generated by mechanical homogenization and were layered over a discontinuous sucrose gradient. After ultracentrifugation, individual fractions were collected from the top of the gradient, and the samples were subjected to SDS-PAGE followed by immunoblotting to identify the organelle contained in each fraction (Fig. 3A). We found the *trans*-Golgi network protein TGN46 and the *cis*-Golgi protein GM130 were predominantly in fractions 2–4, consistent with our previous report (*21*). By contrast, the cytosolic protein HSP90 was predominantly found in fraction 2, the endoplasmic reticulum (ER) luminal protein PDI and the ER membrane protein BAP31 in almost every fraction, the early endosome protein EEA1 in fractions 1–2, and the nuclear protein histone H3 mostly in fractions 6– 11. Thus, fractions 2–4 contain most of the TGN/Golgi material. Strikingly, IPO7 also showed a strong enrichment in fractions 2–4 (Fig. 3A), suggesting that some IPO7 is localized to the TGN/Golgi compartment.

**Figure 3.**
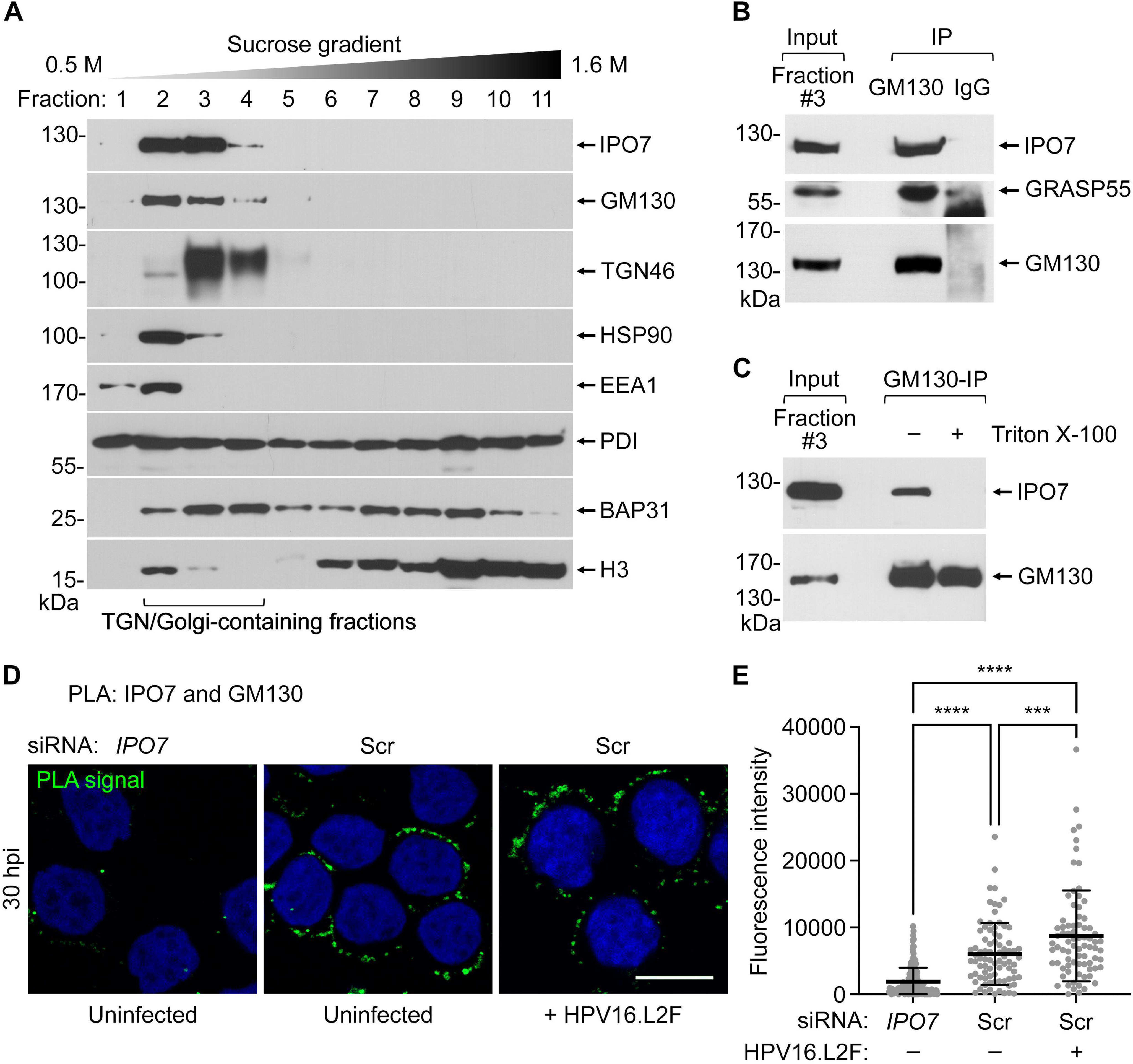
IPO7 associates with the Golgi membrane. **(A)** HeLa cells were mechanically homogenized and fractionated by centrifugation through a 0.5 M to 1.6 M sucrose gradient. To identify the TGN/Golgi-containing fractions, portions of each fraction were subjected to SDS-PAGE followed by immunoblotting with the indicated antibodies. Representative immunoblots for the fractionated cell extracts are shown. Fractions collected from top (fraction 1) to bottom (fraction 11) were displayed from left to right. Fractions 2–4 containing TGN46 and GM130 represent the TGN/Golgi-containing fractions. HSP90, a cytosolic protein; EEA1, an early endosome protein; PDI, an ER luminal protein; BAP31, an ER membrane protein; and H3, a nuclear protein. **(B)** Fraction #3 collected from (A) was subjected to immunoprecipitation with an anti-GM130 antibody or a non-specific IgG as a control. The immunoprecipitated materials were immunoblotted with the indicated antibodies. **(C)** As in (B), except the anti-GM130 immunoprecipitation was performed in the absence or presence of 1% Triton X-100. **(D)** HeLa S3 cells were transfected with 10 nM Scr or *IPO7* siRNA #2 for 48 h and then left uninfected or infected with HPV16.L2F (MOI ∼100) for 30 h. The proximity ligation assay (PLA, signals shown in green) was performed with antibodies recognizing IPO7 and GM130. Nuclei were stained with DAPI (blue). Similar results were obtained in two independent experiments. Scale bar, 10 μm. **(E)** PLA fluorescence intensity per cell in multiple images in (D) was measured, and the individual cell fluorescence intensity values, means, and standard deviations of >80 cells are shown. A two tailed, unequal variance *t*-test was used to determine statistical significance. \*\*\**P* < 0.001; \*\*\*\**P* < 0.0001.

To confirm the localization of IPO7 in the Golgi, we used an antibody against GM130 to IP the Golgi membrane from fraction 3 and found that IPO7 was pulled down (Fig. 3B), indicating that a pool of IPO7 associates with the Golgi membrane. As expected, the Golgi-associated protein GRASP55 also co-precipitated with GM130 (Fig. 3B). When the Golgi IP experiment was repeated in the presence of Triton X-100, a detergent that solubilizes membranes, IPO7 did not co-precipitate with GM130 (Fig. 3C); as a control, IP of the ER membrane protein BAP31 from fraction 3 did not pull down IPO7 regardless of the presence or absence of the detergent (Fig. S3A). Because solubilization of the Golgi membrane prevented IPO7 from co-precipitating with GM130, it is unlikely that IPO7 binds directly to GM130 but is instead pulled down by anti-GM130 because IPO7 associates with the intact Golgi membrane.

To further support the conclusion that IPO7 associates with the Golgi membrane, we performed the proximity ligation assay (PLA) to determine whether IPO7 and GM130 are in close proximity in intact cells. PLA is an immunocytochemistry-based assay that generates fluorescent signals only when two targeted proteins are within 40 nm of each other. Indeed, we found prominent PLA signals generated from antibodies against IPO7 and GM130 in cells transfected with the control Scr siRNA but not in cells transfected with the *IPO7* siRNA #2 (Fig. 3D, quantified in Fig. 3E), showing that a fraction of IPO7 and GM130 are in proximity. The IPO7/GM130 PLA signal was also present in HPV-infected cells.

As assessed by immunoblotting (Fig. S3B) and immunofluorescent staining (Fig. S3C), the GM130 level and distribution remained unchanged after IPO7 KD. Given that IPO7 is dispersed throughout the cell (Fig. S3C), only a fraction of IPO7 is likely proximal to the Golgi membrane. Together, these results suggest that a pool of IPO7 associates with the Golgi membrane. The localization of IPO7 at the Golgi in infected cells raises the possibility that this nuclear import factor might promote HPV infection by directing the virus from the Golgi to the nucleus.

### IPO7 promotes Golgi-to-nucleus transport of HPV

Our data thus far suggest that relatively late steps during HPV entry require Golgi-localized IPO7, a known nuclear entry factor. To determine if IPO7 is required for proper HPV trafficking during entry and, if so, what step of trafficking was affected by IPO7 KD, we assessed the fate of a HPV16.L2F PsV in which the pseudogenome is labeled with the nucleoside analog 5-ethynyl-2’-deoxyuridine (EdU) (*36*). In this experiment, the nuclear envelope is identified by using an antibody against Nesprin2, a nuclear membrane protein. As expected, low background EdU signal was observed in uninfected cells (Fig. 4A; the nuclear, extranuclear, and total EdU fluorescence intensity was quantified in Fig. 4B). In cells transfected with a control siRNA and infected with the EdU-labeled virus for 32 h, EdU signals appeared in the nucleus while a low level of signal was found outside of the nucleus (Fig. 4A and B). These results demonstrate successful trafficking of the incoming PsV and delivery of the EdU viral genome into the nucleus. Importantly, in IPO7 KD cells, the EdU signal in the nucleus was markedly reduced, and the signal accumulated outside of the nucleus (Fig. 4A and B). These findings demonstrate that IPO7 plays a critical role in promoting nuclear arrival of HPV.

**Figure 4.**
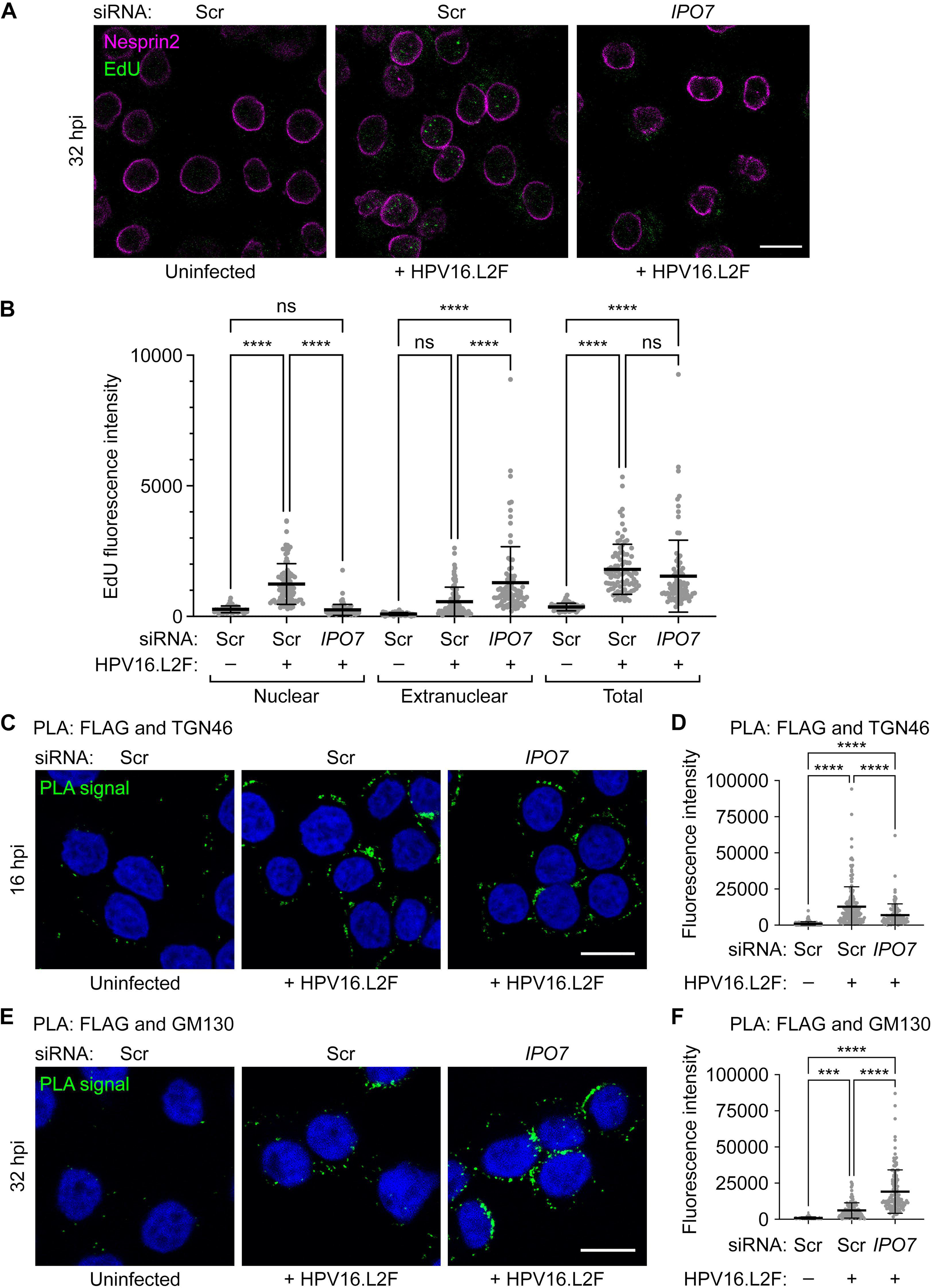
IPO7 promotes Golgi-to-nucleus transport of HPV. **(A)** HeLa S3 cells were transfected with 10 nM Scr or *IPO7* siRNA #2 for 48 h and then left uninfected or infected with HPV16.L2F (MOI ∼150) containing EdU-labeled reporter plasmid. At 32 hpi, cells were fixed and subjected to the Click-iT reaction that generated the EdU signals (green), and immunostained using an antibody recognizing Nesprin2 (magenta). Scale bar, 10 μm. **(B)** Graph shows nuclear, extranuclear, and total cell EdU fluorescence intensity per cell in multiple images as in (A). One-way ANOVA was used to determine statistical significance. ns, not significant; \*\*\*\**P* < 0.0001. **(C)** HeLa S3 cells were transfected with 10 nM Scr or *IPO7* siRNA #2 for 48 h and then left uninfected or infected with HPV16.L2F (MOI ∼100). At 16 hpi, PLA was performed with antibodies recognizing FLAG (to stain L2) and TGN46. Nuclei were stained with DAPI (blue). Similar results were obtained in two independent experiments. Scale bar, 10 μm. **(D)** PLA fluorescence intensity per cell in multiple images in (C) was measured and the individual cell fluorescence intensity values, means, and standard deviations of >100 cells are shown. One-way ANOVA was used to determine statistical significance. \*\*\*\**P* < 0.0001. **(E)** As in (C), except samples at 32 hpi were analyzed by PLA using antibodies recognizing FLAG and GM130. **(F)** PLA fluorescence intensity per cell in multiple images in (E) was analyzed as in (D). \*\*\**P* < 0.001; \*\*\*\**P* < 0.0001.

We then performed PLA to determine where the incoming HPV capsid proteins localized in cells depleted of IPO7. Upon KD of IPO7 and infection with HPV16.L2F PsV for 16 h, the PLA signal between L2-3xFLAG and the TGN protein TGN46 was lower in IPO7 KD cells than in control cells (Fig. 4C and D), suggesting that the depletion of IPO7 did not inhibit trafficking through the TGN which would have caused HPV to accumulate in the TGN. However, the PLA signal between L2-3xFLAG and the *cis*-Golgi marker GM130 at 32 hpi was markedly increased by IPO7 KD (Fig. 4E and F). Thus, in cells depleted of IPO7, HPV transits through the TGN but accumulates in the *cis-*Golgi, presumably due to a block in Golgi exit, explaining why the virus does not reach the nucleus under this compromised condition. Together, these data demonstrate that IPO7 plays a pivotal role in promoting Golgi-to-nucleus trafficking of HPV during virus entry.

### HPV binds to IPO7 during virus entry

Our data thus far identified IPO7 as a Golgi-associated protein that promotes HPV infection by enabling Golgi-to-nuclear transport. We hypothesize that the Golgi-associated IPO7 interacts directly with the cytosol-exposed segment of the HPV L2 protein to deliver HPV to the nucleus. To determine if HPV and IPO7 interact during entry, we first used PLA to assess whether HPV and IPO7 are in proximity in intact infected cells. HeLa S3 cells were uninfected or infected with HPV16.L2F PsV containing a reporter plasmid that encodes luciferase instead of GFP-S (to prevent interference with the PLA fluorescent signal), and PLA for HPV L1 or L2 and IPO7 was performed at 40 hpi. There was little PLA signal in uninfected cells, but we observed pronounced PLA signals between IPO7 and either L1 (Fig. 5A, left panels; quantified in Fig. 5B, left) or L2-3xFLAG (Fig. 5A, right panels; quantified in Fig. 5B, right) in the infected cells. These results indicate that IPO7 is proximal to HPV late during entry.

**Figure 5.**
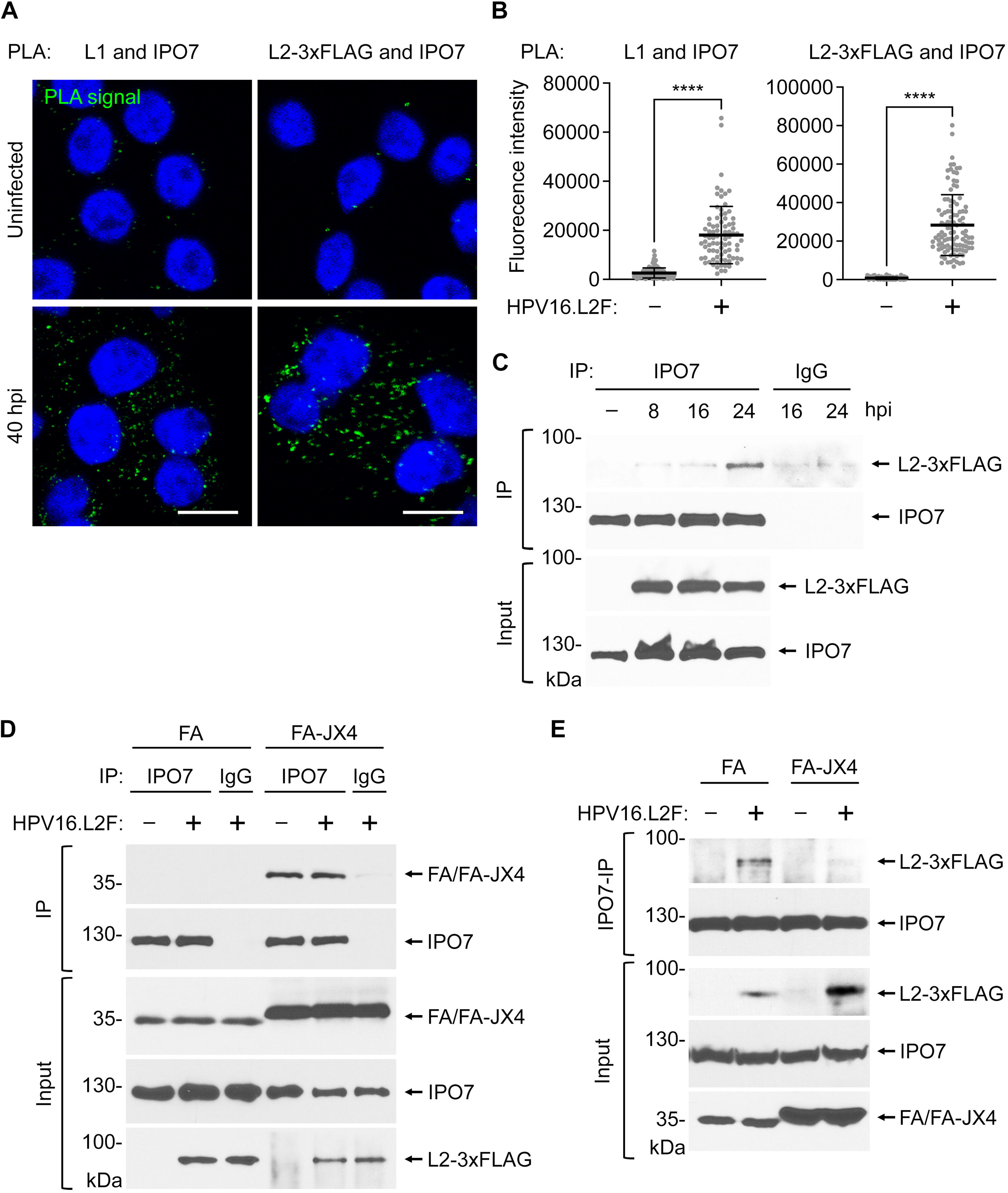
HPV binds to IPO7 during virus entry. **(A)** HeLa S3 cells were uninfected or infected with HPV16.L2F (MOI ∼100). At 40 hpi, PLA was performed with antibodies recognizing IPO7 and either HPV16 L1 (left panels) or FLAG (to detect L2) (right panels). Nuclei were stained with DAPI (blue). Scale bar, 10 μm. **(B)** PLA fluorescence intensity per cell in multiple images as in (A) was measured and the individual cell fluorescence intensity values, means, and standard deviations of >80 cells are shown. A two-tailed, unequal variance *t*-test was used to determine statistical significance. \*\*\*\**P* < 0.0001. **(C)** Whole cell extracts derived from uninfected or HPV16.L2F-infected (MOI ∼1) HeLa cells were collected and lysed at the indicated time points. A portion of the resulting extracts (input) were analyzed by immunoblotting using antibodies recognizing FLAG (to detect L2) and IPO7. The remaining extracts were subjected to immunoprecipitation with an anti-IPO7 antibody or a non-specific IgG antibody as a control. The immunoprecipitated samples were analyzed with anti-FLAG (to detect L2) and anti-IPO7 antibodies. **(D)** HeLa S3 cells expressing FA or FA-JX4 were uninfected or infected with HPV16.L2F (MOI ∼1). Cells were lysed at 24 hpi and the resulting extracts were subjected to immunoprecipitation as in (C) and immunoblotting with antibodies recognizing FLAG (to detect FA and FA-JX4) or IPO7. **(E)** As in (D) except only anti-IPO7 was used for immunoprecipitation and antibody recognizing FLAG was used to detect L2 in the immunoprecipitated samples.

To test whether HPV L2 is in a physical complex with IPO7 during entry, HeLa cells were uninfected or infected with HPV16.L2F for 8, 16, and 24 h. Cells were then harvested, and the resulting cell extract was subjected to IP using an anti-IPO7 antibody. The precipitated material was subjected to SDS-PAGE followed by immunoblotting (Fig. 5C). Anti-IPO7 pulled down L2-3xFLAG predominantly at 24 hpi, whereas low level background signals were detected in samples from earlier time points (or in the negative IgG control). These findings suggest that IPO7 binds to HPV L2 at 24 hpi, a time consistent with nuclear entry events of HPV.

IPO7 was identified by MS as a host protein that specifically interacts with the FA-JX4 traptamer (Fig. 1B). We found that precipitation of IPO7 selectively pulled down FA-JX4 but not FA (Fig. 5D, top panel), in agreement with our IP-MS results (Fig. 1B). The IPO7/FA-JX4 interaction was also observed in uninfected cells. Because FA-JX4 localizes to the TGN/Golgi and traps the virus in this compartment (Fig. S1A) (*18*), we hypothesized that binding of FA-JX4 to IPO7 disrupts the ability of IPO7 to bind HPV L2. To test this idea, control FA-and FA-JX4-expressing cells were infected with HPV16.L2F PsV. Extracts prepared from cells 24 hpi were subjected to IP using an anti-IPO7 (or control IgG) antibody, and the precipitated material was evaluated by SDS-PAGE followed by immunoblotting for FLAG-tagged L2. The IPO7-HPV L2 interaction was readily detected in cells expressing the control protein FA but was markedly diminished in cells expressing FA-JX4 (Fig. 5E, top panel). These results suggest that FA-JX4 inhibits the ability of HPV to engage IPO7, which is required to transport the virus to the nucleus, thus explaining, at least in part, how FA-JX4 impairs HPV entry.

### The HPV L2-IPO7 interaction requires COPI

We previously reported that HPV recruits the COPI complex upon arrival to the TGN to enable transit across the Golgi stacks (*21*). The interaction between HPV L2 and COPI could be detected beginning at 16 hpi. HPV L2 binds to IPO7 at 24 hpi, presumably after the virus has transported across the TGN/Golgi stacks, a step that requires COPI. Furthermore, in COPI knockdown cells the virus accumulates in both the TGN and the Golgi, whereas in the IPO7 knockdown cells the virus exits the TGN and accumulates in the Golgi. Therefore, we hypothesize that COPI is required for the HPV L2-IPO7 interaction. To test this, HeLa cells were transfected with Scr siRNA or siRNAs against *COPA* and *COPG1* (to co-deplete the α-and γ1-COP subunits, respectively), followed by infection with HPV16.L2F. The cells were lysed at 24 hpi, and the resulting extracts were IPed with antibody recognizing IPO7. We found that anti-IPO7 pulled down L2-3xFLAG from extracts of control but not COPI KD cells (Fig. 6A, top panel), indicating that the IPO7-HPV L2 interaction requires COPI. In contrast, IPO7 depletion did not inhibit the HPV L2-COPI interaction (Fig. 6B, top panel): anti-γ-COP immunoprecipitated HPV16 L2-3xFLAG at 24 hpi even in the absence of IPO7; in fact, the HPV L2-COPI interaction was modestly enhanced in the absence of IPO7. These findings strongly suggest that the HPV L2-IPO7 interaction occurs downstream of COPI action after the virus has transited across the Golgi stacks.

**Figure 6.**
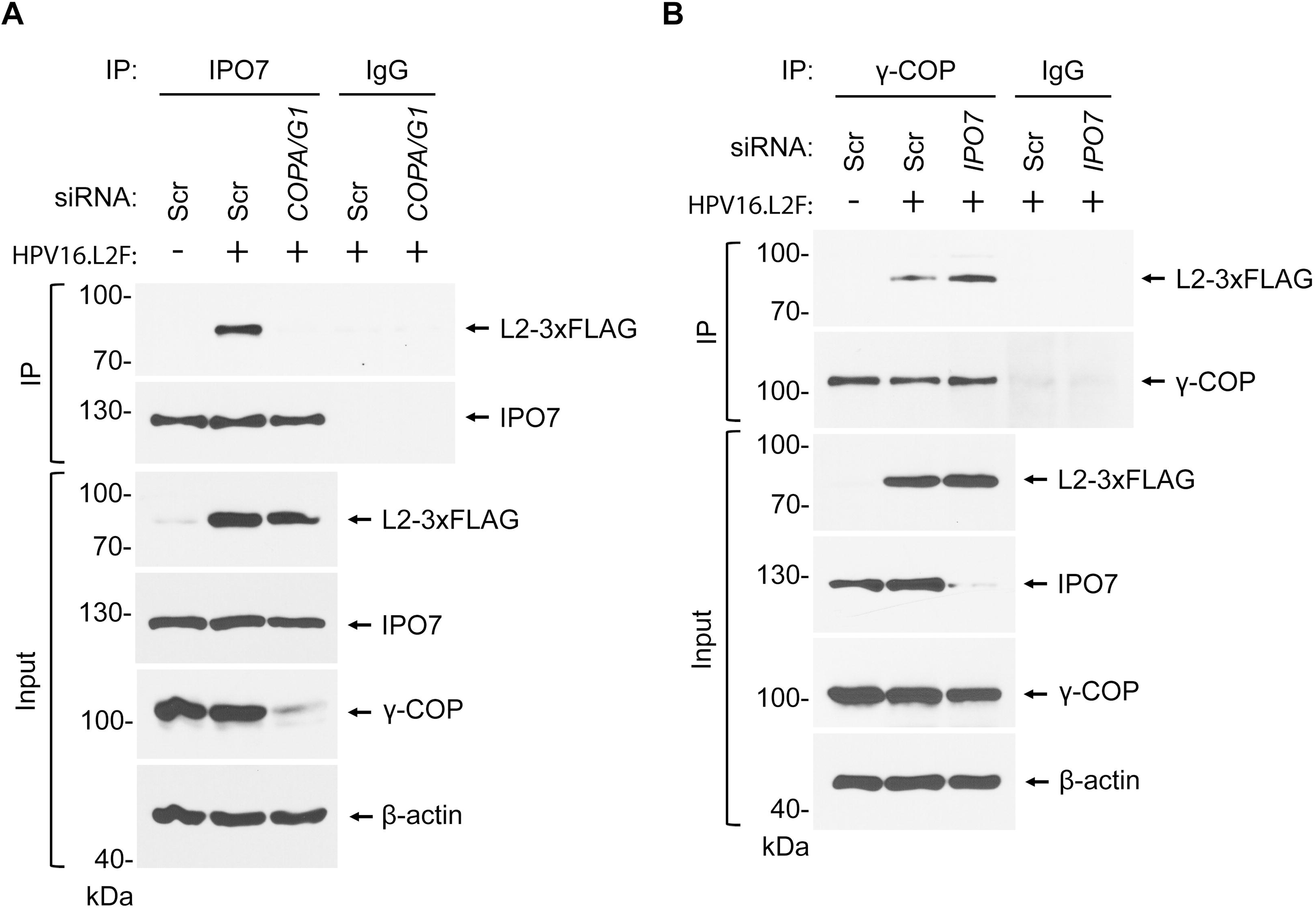
The HPV L2-IPO7 interaction requires COPI. **(A)** HeLa cells treated with 2 nM Scr or a mixture of 1 nM *COPA* and 1 nM *COPG1* (*COPA/G1*) siRNA were uninfected or infected with HPV16.L2F PsV (MOI ∼1). At 24 hpi, cells were lysed and a portion of the resulting extracts (input) were analyzed by immunoblotting using antibodies recognizing FLAG, IPO7, γ-COP, or β-actin (as a loading control). The remaining extracts were subjected to immunoprecipitation with an anti-IPO7 antibody or a non-specific IgG as a negative control. Immunoprecipitated samples were analyzed with anti-FLAG and anti-IPO7 antibodies. **(B)** As in (A), except HeLa cells were transfected with 10 nM Scr or *IPO7* siRNA #2 rather than *COPA/G1* siRNA before infection, and an anti-γ-COP antibody or a non-specific IgG was used for immunoprecipitation. The immunoprecipitated samples were analyzed with anti-FLAG and anti-γ-COP antibodies.

### IPO7 binds directly to the C-terminal NLS motif of HPV L2

We next determined the sequence motifs in HPV L2 that interact with IPO7. IPO7 typically recognizes the NLS of cellular cargos to deliver them to the nucleus through the nuclear pore (*37*). Therefore, we hypothesize that HPV L2 engages IPO7 via an NLS in L2 to promote nuclear targeting of HPV. There are three predicted NLS motifs in HPV16 L2, each enriched with arginine (R) and lysine (K) residues that typically define an NLS motif (*27, 30, 38*). These putative NLS motifs are located at the N-terminus (nNLS), central region (mNLS), and C-terminus (cNLS) of L2 (Fig. 7A); the highly conserved cNLS, composed of six R/K amino acids in HPV16 (RKRRKR), is also termed the CPP because it exerts an intrinsic membrane penetration activity that mediates penetration of L2 across the endosome membrane during the early steps of HPV entry (*12*). Because the nNLS is removed after cleavage by furin during early entry steps (*39*), this NLS is not likely to participate in nuclear transport events of the incoming HPV. We therefore tested whether the mNLS or the cNLS of L2, which are expected to be exposed to the cytosol when L2 is inserted into the membrane (Fig. 7A), interact with IPO7.

**Figure 7.**
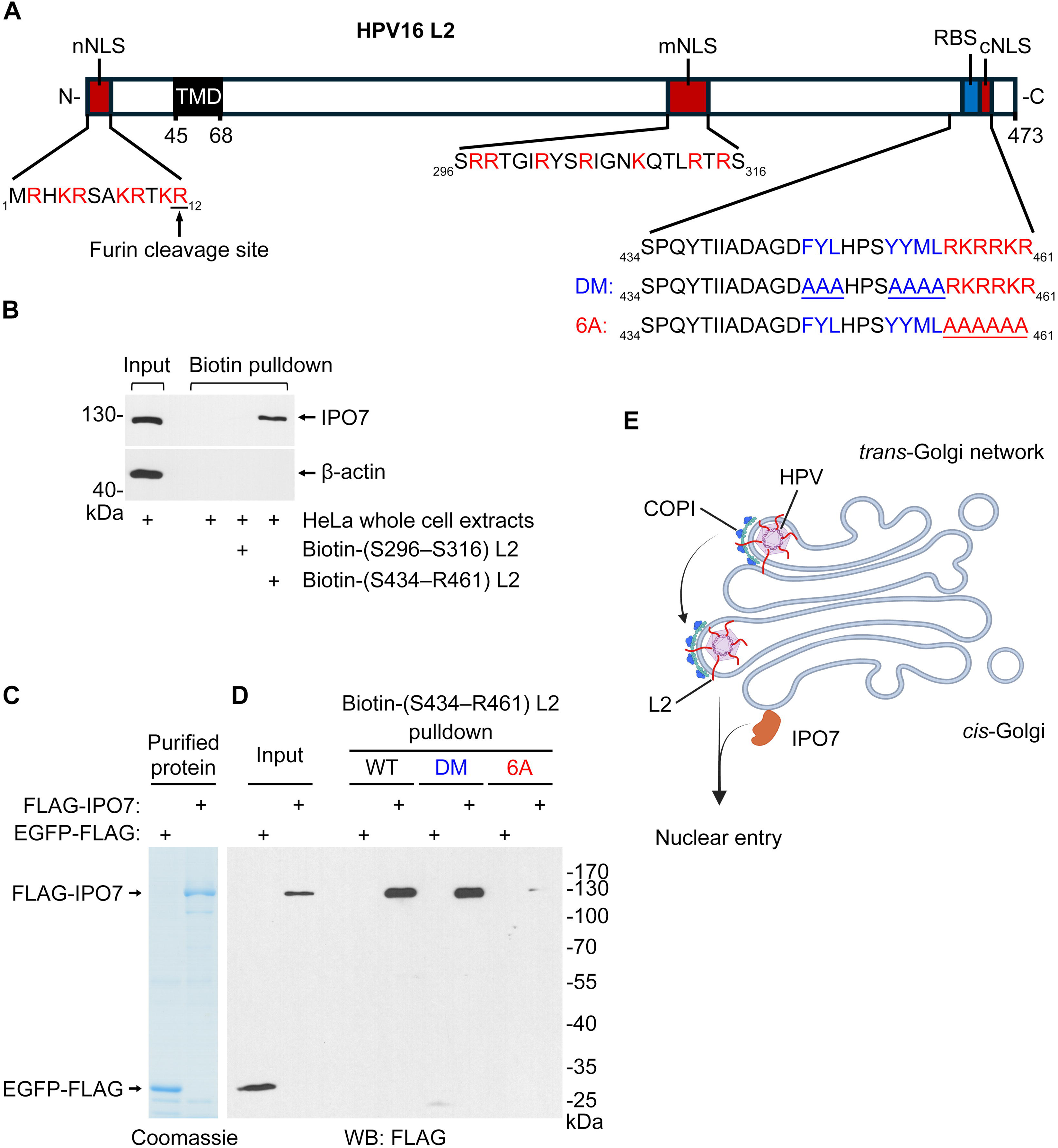
IPO7 binds directly to the C-terminal NLS motif of HPV L2. **(A)** Diagram of the nuclear localization sequences (NLSs) in HPV16 L2. Three postulated NLSs are indicated as red boxes, and the basic amino acids in the NLSs are highlighted in red. The predicted transmembrane domain (TMD) spanning amino acid 45–67 is denoted as a black box. The retromer binding site (RBS) is indicated as a blue box, the amino acids required for retromer-binding are highlighted in blue, and the corresponding RBS mutation to alanines are underlined in the DM peptide. The cNLS mutation to alanines are underlined in the 6A peptide. **(B)** Whole cell extracts of uninfected HeLa cells were incubated with N-terminal biotin-tagged peptides containing residues S296–S316 or S434–R461 of HPV16 L2. The pulldown experiment without peptide was used as a negative control. Samples captured with Streptavidin-beads were analyzed by immunoblotting using antibodies recognizing IPO7 or β-actin. **(C)** Coomassie stain of the purified proteins. EGFP-FLAG, C-terminal FLAG-tagged EGFP. FLAG-IPO7, N-terminal FLAG-tagged IPO7. **(D)** N-terminal biotin-tagged peptides containing residues S434–R461 of HPV16 L2 (WT) and the corresponding DM and 6A mutant peptides as indicated in (A) were incubated with purified EGFP-FLAG or FLAG-IPO7. Samples captured with Streptavidin-beads were analyzed by immunoblotting using an antibody recognizing FLAG. **(E)** During entry, HPV is transported from the TGN and across the Golgi stacks to the *cis*-Golgi in a COPI-dependent mechanism. After arrival in the Golgi, HPV is targeted to the nucleus via the activity of the Golgi-associated IPO7 nuclear import receptor. Figure created with BioRender.com.

To test this possibility, we synthesized N-terminal biotinylated peptides composed of amino acid 296–316 of L2 [Biotin-(S296–S316) L2] that harbors the mNLS sequence or of amino acid 434–461 of L2 [Biotin-(S434–R461) L2] that contains the cNLS sequence (Fig. 7A). Extracts from uninfected HeLa cells were incubated with either of these peptides (or no peptide addition as the negative control), the peptides and any associated proteins were pulled down using Streptavidin beads, and the precipitated material was subjected to SDS-PAGE and immunoblotting for IPO7. The peptide harboring cNLS [Biotin-(S434–R461) L2] but not the mNLS peptide pulled down IPO7 (Fig. 7B, top panel), whereas neither of the peptides pulled down the control protein β-actin (Fig. 7B, bottom panel). These results indicate that a short segment of L2 containing the cNLS is sufficient to pull down IPO7 from cell extracts.

To evaluate if the IPO7-peptide interaction is direct, we performed a similar biotin-pulldown assay, with purified FLAG-tagged IPO7 (FLAG-IPO7) and the control EGFP-FLAG proteins instead of the cell extract (Fig. 7C). We found that the Biotin-(S434– R461) L2 peptide (WT) efficiently pulled down FLAG-IPO7 but not EGFP-FLAG (Fig. 7D), demonstrating that IPO7 binds to this peptide directly. However, if the basic RKRRKR cNLS sequence in the Biotin-(S434–R461) L2 peptide was mutated to AAAAAA generating the 6A mutant (Fig. 7A), this peptide no longer pulled down FLAG-IPO7 (Fig. 7D). By contrast, if the retromer binding site (RBS) sequences (FYL/YYML) that lie upstream of the RKRRKR sequence (*40*) were mutated to A (generating the DM peptide unable to bind retromer, Fig. 7A), this peptide pulled down FLAG-IPO7 with a similar efficiency as the WT peptide (Fig. 7D). We conclude that the cNLS/CPP at the C-terminus of L2 is critical for engaging IPO7, consistent with a previous report showing that IPO7 recognizes sequences enriched in basic amino acids (*37*). In contrast, the retromer binding site in this peptide is not required for IPO7 binding.

## Discussion

Although HPV is responsible for approximately 5% of all human cancers, effective anti-viral therapies remain elusive due in part to an incomplete understanding of the virus entry mechanism. Early during entry, HPV undergoes endocytosis and reaches the endosome. In this compartment, the virus capsid protein L2 is inserted across the endosome membrane so that most of L2 is exposed to the cytosol (*12, 14*). In this membrane-inserted topology, the cytosol-exposed region of L2 recruits critical cytosolic sorting factors that mediate proper trafficking to the nucleus for infection. For example, HPV exploits COPI, BicD2/dynein, and Rab6a to traffic from the TGN in a retrograde direction to cross the Golgi stacks (*19–21*). However, after traversing the Golgi, it is not known how HPV is targeted to the nucleus. In this study, we show that HPV relies on the Golgi-associated importin β member IPO7, a component of the nuclear pore import machinery, to target to the nucleus and cause infection (Fig. 7E).

We previously reported that expression of an artificial transmembrane protein (named FA-JX4) traps HPV in the TGN/Golgi and blocks virus infection (*18*). Based on this observation, we hypothesized that FA-JX4 binds to a putative host factor that is required for infection, preventing the host factor from engaging HPV. Using an unbiased proteomics strategy, we identified IPO7 as an FA-JX4 binding-partner that supports infection. IPO7 was not identified in previous biochemical and genetic screens for HPV entry factors. Similarly, the Rab7 GTPase activating protein, TBC1D5, was identified as an HPV entry factor by studying a different inhibitory protein, FA-JX2, which traps incoming HPV in the endosome (*31*). These results emphasize the value of using orthogonal approaches to dissect HPV entry (*41*).

By using a cell fractionation approach and a Golgi-IP method, we found that a pool of IPO7 is associated with the Golgi membrane in infected and uninfected cells. This finding is further supported by PLA analysis, which demonstrated that IPO7 is proximal to the Golgi in intact cells. The strategic positioning of IPO7 at the Golgi is suited for HPV entry because incoming HPV traffics through the Golgi *en route* to the nucleus. IPO7 lacks a transmembrane domain, so the mechanism by which IPO7 engages the Golgi membrane is unclear. It is possible that a Golgi-resident transmembrane/peripheral protein serves as an IPO7 binding-partner, anchoring importin β to the Golgi membrane. Alternatively, IPO7 may simply be localized to the Golgi because it associates with various Golgi-associated factors as cargos, such as the SREBP-2 transcription factor, whose transport from the Golgi to the nucleus was reported to depend on importin β family proteins (*42, 43*).

Functionally, we demonstrated that chemical inactivation of the importin β family proteins with importazole robustly blocked HPV infection if added prior to 24 hpi, a time point when a majority of HPV has already reached the nucleus (*23, 35*). We then used RNA interference against importin β members that interacted with FA-JX4 to show that depletion of IPO7 (and other importin β family members) led to a substantial inhibition of HPV infection by different HPV types and in different cell lines, indicating that IPO7 plays a key role in infection. IPO5 but not IPO7 was identified in a large-scale siRNA screen for host factors involved in HPV infection (*23*). However, IPO5 KD disrupts the cell cycle (Fig. S2D), consistent with a previous report (*44*), which raises the question of whether loss of IPO5 causes secondary effects on HPV entry due to the impact on cell cycle progression.

We used labeled encapsidated DNA and PLA for HPV capsid proteins to examine the fate of HPV in cells depleted of IPO7. These studies showed that in these cells the virus is trapped in the Golgi and that the infectious virus cannot reach the nucleus. These observations provide a simple explanation for the inhibition of HPV infection by IPO7 KD and indicate that IPO7 is responsible for Golgi-to-nucleus transport of HPV. Published work from other laboratories showed that IPO7 also plays an important role in infection by other viruses. IPO7 directly translocates the flavivirus core protein into the nucleus to promote infectious virus production, even though viral genome replication occurs in the cytoplasm (*45*). Similarly, the retrovirus HIV-1 exploits IPO7 to maximize nuclear import of its genome (*46*), as well as to facilitate replication via interaction between the viral integrase and IPO7 (*47*).

Our time-course co-IP experiments demonstrated that HPV L2 is in a complex with IPO7 by 24 hpi, after the virus enters the TGN/Golgi, consistent with the idea that IPO7 acts after the virus has transited across the Golgi stacks (Fig. 7E). In contrast, HPV and retromer are in a complex by 8 hpi (*16*), and HPV and COPI are in a complex by 16 hpi (*21*). In addition, COPI depletion caused HPV PsV to accumulate in both the TGN and the Golgi (*21*), whereas IPO7 depletion caused HPV accumulation in the Golgi (i.e., GM130 compartment) but not the TGN (i.e., TGN46 compartment). These results suggest that COPI acts prior to IPO7 during retrograde trafficking. Indeed, depletion of COPI interfered with the ability of HPV L2 to interact with IPO7 in infected cells, suggesting that COPI-dependent transport of HPV from the TGN to the Golgi stacks and across the Golgi stack is upstream of IPO7 action during HPV entry.

Importin β typically binds to highly basic NLSs. Our *in vitro* binding studies revealed that IPO7 interacts directly with a postulated NLS motif located at the C-terminus of HPV L2. IPO7 does not bind to mNLS *in vitro*, but recent work indicated that this element has only weak nuclear localization activity, and it has been reclassified as a nuclear retention or chromatin tethering signal (*30, 38, 48*). The inability of this element to bind IPO7 suggests that in our assay IPO7 is not binding promiscuously to all basic segments. The cNLS motif overlaps with a CPP sequence that mediates penetration of L2 across the endosome membrane (*12*), indicating that the C-terminus of L2 is responsible for at least two independent steps during HPV entry. Because all known mutations in this basic segment inhibit L2 membrane protrusion activity and prevent entry of HPV into the retrograde pathway, we cannot assess the effects of these mutations on IPO7 binding or late entry events in infected cells. Our finding that FA-JX4 interferes with IPO7-HPV binding provides a simple explanation for the inhibitory effect of the traptamer on HPV trafficking. We note that FA-JX4 caused HPV to accumulate in the TGN and the Golgi (*18*), whereas the block caused by IPO7 depletion was confined to the Golgi. It is possible that FA-JX4 has activities other than inhibition of L2-IPO7 binding, or that FA-JX4 exerts a more complete block to IPO7 activity than does siRNA-mediated KD.

Although classic nuclear entry of small cellular cytosolic cargoes typically relies on an importin α and β heterodimer for delivery to the nuclear pore (*25*), our MS analysis did not identify any importin α family members as FA-JX4-binding partners, suggesting that IPO7 targets HPV to the nucleus without assistance from importin α proteins. Indeed, importin β family proteins have been shown to be sufficient to drive nuclear entry of some cellular proteins ((*42, 49, 50*). However, because a previous study suggested that the importin α member KPNA2 supports HPV infection (*26*), it is possible that IPO7 cooperates with KPNA2 to target HPV to the nucleus. Clearly this question deserves future investigation.

NEB during mitosis is thought to enable entry of HPV into the nucleoplasm (*22–24*). However, it is largely unknown whether NEB-dependent nuclear entry of HPV involves the classic nuclear import machinery, including importin α/β, the Ran GTPase, or nucleoporins (NUPs) of the nuclear pore complex (NPC). NUPs appear to be dispensable for HPV infection because depletion of NUP153, the essential component of the NPC basket (*51*), does not affect HPV16 infection (*23*). Our study suggests that at least one component of the nuclear import machinery, IPO7, plays a pivotal role during or prior to NEB-mediated nuclear entry of viral components. We postulate that during HPV entry, the importin β member IPO7 targets the virus to the nuclear envelope without using the channel function of the NPC to gain nuclear entry. Instead, we propose that the IPO7-dependent targeting step positions the virus next to the nucleus so that upon NEB during mitosis, the virus can efficiently enter the nucleus. The association of the L2 protein with chromatin then traps the L2 protein and associated viral DNA in the nucleus when the nuclear envelope reforms (*30, 48*). Our results raise the possibility that some cellular cargos also use a combination of the canonical nuclear import machinery and NEB to gain nuclear entry.

## Materials and Methods

### Antibodies and inhibitors

Antibodies and inhibitors used in this study are listed in Table 1.

**Table 1.** Antibodies and inhibitors.

| **Antibodies** |  |  |  |
| --- | --- | --- | --- |
| **Antigen** | **Species** | **Cat # / Source** | **Application** |
| IPO1 | rabbit | 10077-1-AP; Proteintech | WB |
| IPO4 | rabbit | 11679-1-AP; Proteintech | WB |
| IPO5 | mouse | sc-55527; Santa Cruz Biotechnology | WB |
| IPO7 | mouse | sc-365231; Santa Cruz Biotechnology | WB, IF, PLA, IP |
| IPO7 | rabbit | 28289-1-AP; Proteintech | IP, PLA |
| IPO8 | mouse | sc-398854; Santa Cruz Biotechnology | WB |
| IPO9 | rabbit | SAB4200155; Millipore Sigma | WB |
| GRASP55 | mouse | sc-271840; Santa Cruz Biotechnology | WB |
| FLAG | mouse | F3165; Millipore Sigma | WB, IF, IP, PLA |
| FLAG | rabbit | 14793S; Cell Signaling | PLA |
| γ-COP | rabbit | 12393-1-AP; Proteintech | WB, IP |
| PS1 | rabbit | 5643S; Cell Signaling | WB |
| TGN46 | rabbit | 13573-1-AP; Proteintech | WB, PLA |
| GM130 | rabbit | ab52649; abcam | WB, IF, PLA, IP |
| EEA1 | rabbit | C45B10; Cell Signaling | WB |
| BAP31 | rat | MA3-002; Invitrogen | WB, IP |
| PDI | mouse | ab2792; abcam | WB |
| Histone H3 | rabbit | 9715S; Cell Signaling | WB |
| GFP | mouse | 632380; Takara Bio Clontech | WB |
| β-actin | rabbit | 4967S; Cell Signaling | WB |
| HSP90 | mouse | sc13119; Santa Cruz Biotechnology | WB |
| HA | rat | ROAHAHA; Roche | WB, IP |
| HPV16 L1 | mouse | 554171; BD | PLA |
| Nesprin2 | rabbit | ab314872; abcam | IF |

| Other antibodies | Species | Cat # / Source | Application |
| --- | --- | --- | --- |
| anti-mouse IgG peroxidase | goat | A4416; Millipore Sigma | WB |
| anti-mouse IgG peroxidase | goat | W4021; Promega | WB |
| anti-rabbit IgG peroxidase | goat | A4914; Millipore Sigma | WB |
| anti-rabbit IgG peroxidase | goat | W4011; Promega | WB |
| anti-rat IgG peroxidase | rabbit | A5795; Millipore Sigma | WB |
| anti-mouse Alexa Fluor 488 | donkey | A21202; Thermo Fisher | IF |
| anti-mouse Alexa Fluor 488 | goat | A11029; Thermo Fisher | IF |
| anti-rabbit Alexa Fluor 594 | goat | A11037; Thermo Fisher | IF |
| anti-rabbit Alexa Fluor 647 | goat | A21245; Thermo Fisher | IF |
| Normal rabbit IgG | rabbit | 30000-0-AP; Proteintech | IP |
| Inhibitors |  |  |  |
| Compound | Solvent | Cat # / Source | Concentration |
| XXI<br>(inhibiting γ-secretase) | DMSO | 565790; Millipore Sigma | 1 μM |
| Importazole<br>(inhibiting importin $\beta$) | DMSO | 401105; Millipore Sigma | 1–10 μM |

### DNA constructs

For PsV production, the p5sheLL.L2F, p16sheLL.L2F, and p18sheLL.L2F constructs were modified from p5sheLL, p16sheLL, and p18sheLL plasmids (gifts from Dr. John Schiller, National Cancer Institute, Rockville, MD; Addgene plasmids #46953, #37320, and #37321), respectively, as previously described (*21*). The pcDNA3.1 plasmid expressing GFP with a C-terminal S-tag was used as the pseudoviral genome (reporter plasmid) as previously described (*21, 52*). The pCINeo-Gluc construct generated in the previous study (*21*) was used as the reporter of HPV16 PsV for the PLA assay and EdU staining. For KD-rescue experiments, pCMVTNT-HA-mCherry generated in the previous study (*21*) was used as the control vector. The *IPO7* coding sequence was amplified from pUC19-mIPO7 (MG5A1798-U, Sino Biological) and cloned into the pCMVTNT-HA-COPG1-mCherry plasmid (*21*) to replace *COPG1*. Site-directed mutagenesis was used to introduce siRNA-resistant mutations in the *IPO7* siRNA #2 target site. For recombinant protein production, the siRNA-resistant *IPO7* coding sequence was amplified from pCMVTNT-HA-IPO7-mCherry and cloned into pFLAG-CMV2 vector in-frame with the 5’ FLAG to encode FLAG-IPO7. pcDNA3.1(-)-EGFP-FLAG was described previously (*53*). All plasmids constructed in this study were verified by sequencing.

### Cell culture

HeLa (ATCC, Cat# CCL-2), HeLa-S3 (ATCC, Cat# CCL-2.2), and SiHa (ATCC, Cat# HTB-35) cells were obtained from American Type Culture Collection (ATCC). HaCaT cells were purchased from AddexBio Technologies. HEK 293TT cells were obtained from Dr. Christopher Buck (National Cancer Institute, Rockville, MD). HeLa cells (ATCC, Cat# CCL-2) were used throughout this study unless otherwise specified. HeLa-tTA clonal cells derived from HeLa S3 stably expressing FA or FA-JX4 were described previously (*18*). All cell lines were cultured in Dulbecco’s modified Eagle’s medium (DMEM) supplemented with 10% fetal bovine serum (FBS), penicillin and streptomycin, and incubated at 37°C and 5% CO_2_. Cell lines were authenticated by using the ATCC cell line authentication service.

### HPV PsV production

HPV PsV were produced by co-transfecting HEK 293TT cells with a reporter plasmid and p16sheLL, p16sheLL.L2F, p5sheLL.L2F, or p18sheLL.L2F using polyethyleneimine (PEI, Polysciences). Packaged PsVs were purified by density gradient centrifugation in OptiPrep (Millipore Sigma) as described (*54, 55*). Purified PsVs were subjected to SDS-PAGE followed by staining using SimplyBlue SafeStain (Invitrogen) to assess the quality and quantity of L1 and L2.

### Immunoprecipitation and Mass-Spectrometry

Cells stably expressing FA or FA-JX4 were seeded in 15-cm plates and grown to ∼80% confluency and infected with HPV16 PsV (∼25 μg L1). At 24 hpi, cells from three 15-cm plates were pooled and lysed with 1% Triton X-100 in HN buffer [50 mM HEPES, 150 mM NaCl, 1 mM phenylmethylsulfonyl fluoride (PMSF)] on ice for 15 min. The resulting extracts were centrifuged at 16,100 *g* at 4°C for 10 min, and the supernatants were then incubated with anti-FLAG M2 antibody (∼8 μg antibody per 1 mL of lysate) (F3165; Millipore Sigma) at 4°C overnight. The immune complex was captured with protein G-coated magnetic beads (Invitrogen, 10003D) at 4°C for 2 h. Beads were washed on ice four times with 0.1% Triton X-100 in HN buffer. Bound proteins were eluted twice with 250 μg/mL 3xFLAG peptide (Millipore Sigma) in HN buffer containing 0.1% Triton X-100 at room temperature for 30 min. After the 3xFLAG peptide elution, the beads were incubated with 1X SDS sample buffer at 95°C for 10 min to elute the material still bound to the beads. A portion of the 3xFLAG peptide eluate was analyzed by SDS-PAGE followed by immunoblotting to confirm that equivalent amounts of traptamers (FA or FA-JX4) were precipitated. The remaining eluate was treated with 10% trichloroacetic acid and incubated on ice for 10 min. The sample was subjected to centrifugation, and the precipitated material washed twice with ice-cold acetone. The precipitate was subject to mass-spectrometry analysis at the Taplin Mass-Spectrometry Core Facility (Harvard Medical School). LC/MS-MS was performed using an Orbitrap mass spectrometer (Thermo Fisher Scientific).

### HPV infectivity

HeLa, SiHa, or HaCaT cells were treated as indicated and infected with HPV16.L2F, HPV5.L2F, or HPV18.L2F PsV [multiplicity of infection (MOI) ∼0.3] containing a GFP reporter construct. Where indicated, inhibitors (1 µM XXI or 1–10 µM IPZ) were added at the time of infection or after infection at 8-h time intervals. An equivalent volume of the carrier solvent DMSO was added to the control cultures. At 48 hpi, cells were washed with PBS and lysed in HN buffer containing 1% Triton X-100 on ice for 10 min. Cells were centrifuged at 16,100 *g* at 4°C for 10 min. The resulting supernatant was incubated with SDS sample buffer containing 2-mercaptoethanol, denatured by incubating at 95 °C for 10 min, and analyzed by SDS-PAGE followed by immunoblotting for GFP-S expression using an antibody recognizing GFP. Alternatively, infected cells were harvested by trypsinization, resuspended in ice-cold PBS containing 2% FBS and 0.1 μg/mL DAPI, followed by the flow cytometry analysis performed with a Bio-Rad ZE5 cell analyzer (University of Michigan Flow Cytometry Core Facility). After gating for size and singlets, the population of DAPI-negative cells (∼1 × 10^4^ live cells) was analyzed for GFP fluorescence. For knockdown-rescue experiments, trypsin without phenol red was used to prevent interference of mCherry signals and the infectivity was determined as GFP-positive population among mCherry-positive cells.

### Immunoblotting

Protein samples were separated via SDS-PAGE and transferred to nitrocellulose membranes (Amersham, Millipore Sigma). The membranes were then subjected to incubation with primary antibodies in Tris-buffered saline (TBS) supplemented with 3% skim milk and 0.2% Tween-20 at 4°C overnight and then to incubation with horseradish peroxidase (HRP)-conjugated secondary antibodies in TBS/3% skim milk/0.2% Tween-20 at room temperature for 1 h. HRP-substrates (Immobilon, Millipore Sigma) were used to generate the chemiluminescence signals that were detected by exposing to X-ray films (Ece Scientific, Thermo Fisher Scientific).

### siRNA transfection

siRNA oligos used in this study are listed in the Table 2. 12-well plates were seeded with 7 × 10^4^ HeLa, 4 × 10^5^ HeLa S3, or 5 × 10^4^ HaCaT cells per well and simultaneously reverse-transfected with 10 nM of indicated siRNA by using Lipofectamine RNAiMAX (Thermo Fisher Scientific) according to the manufacturer’s instructions.

**Table 2.**
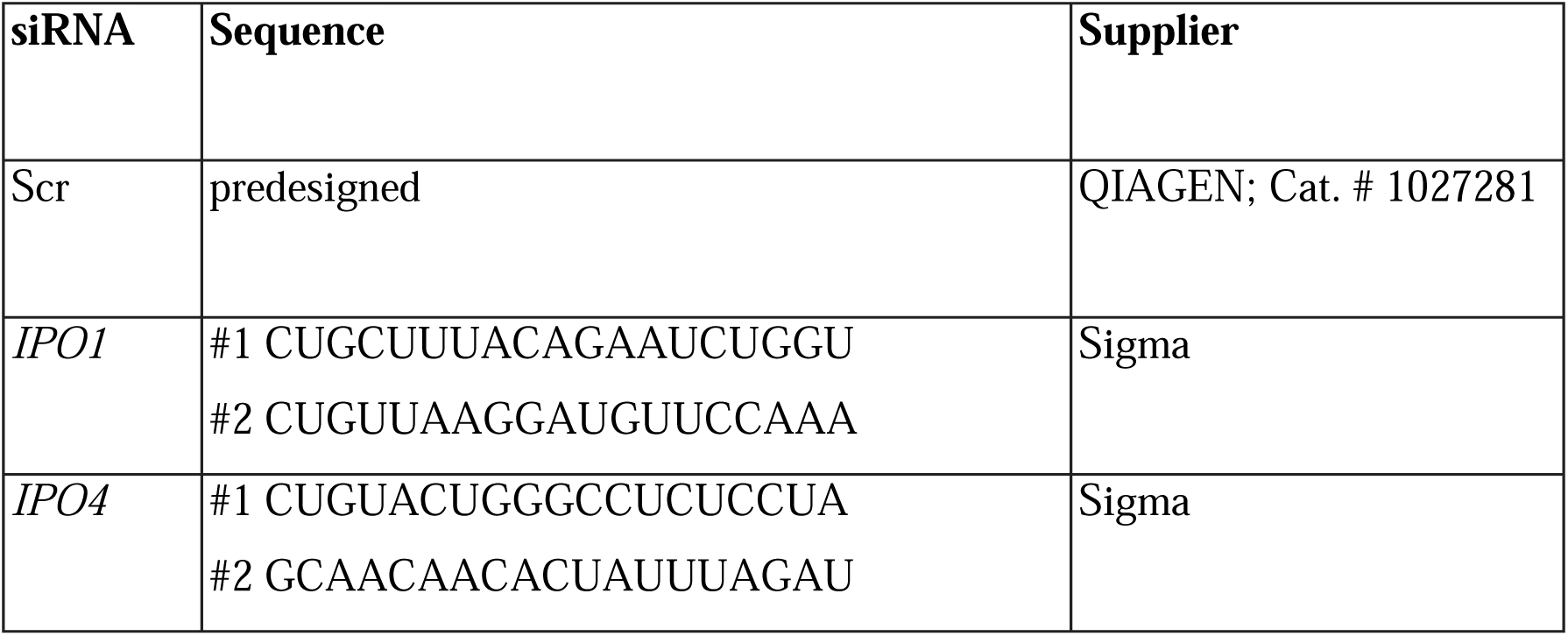
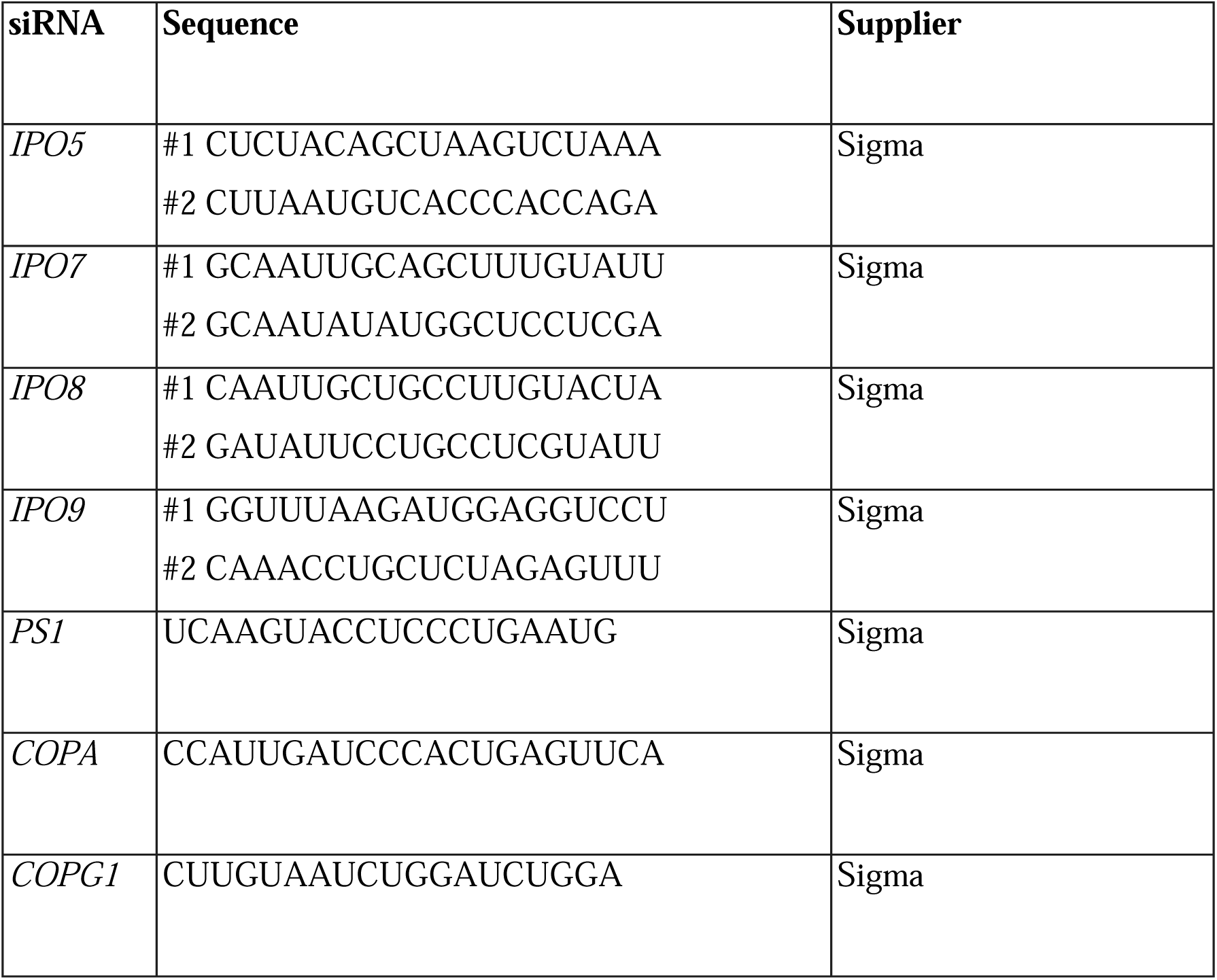
siRNA oligos for gene-knockdowns.

### DNA transfection

For knockdown-rescue experiments, HeLa cells were grown to ∼80% confluency in 10-cm plates and then transfected with 3 μg of pCMVTNT-HA-mCherry or 8 μg of pCMVTNT-HA-IPO7-mCherry plasmids for 24 h using the transfection reagent FuGENE HD (Promega). The cells were then trypsinized and 1.5 × 10^5^ cells per well were seeded in 6-well plates and reverse-transfected with 10 nM of Scr siRNA or *IPO7* siRNA #2 using the transfection reagent Lipofectamine RNAiMAX (Thermo Fisher Scientific) for 48 h. Attached cells were washed with PBS twice and infected with HPV16.L2F (MOI ∼0.3) for another 48 h. Infectivity was analyzed by flow cytometry to determine the fraction of GFP-expressing cells among mCherry-expressing cells. A portion of cells were collected for immunoblot analyses to verify the expression of rescue constructs and IPO7 KD. For recombinant protein production, HEK 293TT cells were seeded in 10-cm plates for 24 h to reach ∼80% confluency and then transfected with 5 μg of indicated plasmids for 48 h using PEI.

### Cell cycle analysis

DNA content was measured using Hoechst 33342 staining and flow cytometry analysis. Non-permeabilized cells were incubated with 10 μg/mL Hoechst 33342 (Biotechne) at 37°C for 30 min. Cells were then washed with PBS, harvested by trypsinization, and resuspended in ice-cold PBS containing 2% FBS. Flow cytometry was performed with a Bio-Rad ZE5 cell analyzer (University of Michigan Flow Cytometry Core Facility). The percentage of cells in G1, S, or G2/M phases was determined from control cell cultures using FlowJo software (BD). Data from three independent experiments are presented as the percentage of cells in each phase. A two-tailed, unequal variance *t*-test was used to determine statistical significance between the knockdown and control cells for each cell cycle phase.

### Sucrose gradient fractionation

Sucrose gradient fractionation was carried out to isolate TGN/Golgi-containing cellular fractions as described (*21*). In brief, HeLa cells were plated in 15-cm plates and grown to ∼80% confluency. Two 15-cm plates per preparation were processed as follows. Cells were trypsinized and resuspended in HBS buffer [10 mM HEPES pH 7.2, 1 mM Mg(OAc)_2_, 0.25 M sucrose, 1 mM EDTA, 1 mM dithiothreitol (DTT), 1 mM PMSF], and homogenized by passing 31 times through an 8-µm clearance ball bearing homogenizer on ice (Isobiotech). The homogenate was centrifuged at 2,000 *g* for 30 min at 4°C, and the resulting supernatant (∼1.2 mL) was layered over a sucrose gradient comprised of 1 mL 1.6 M, 1 mL 1.4 M, 2 mL 1.2 M, 3 mL 1.0 M, 2 mL 0.8 M, and 1 mL 0.5 M sucrose in MEB buffer (50 mM Tris-HCl pH 7.4, 50 mM KCl, 20 mM β-glycerol phosphate, 15 mM EGTA, 10 mM MgCl_2_, 1 mM DTT) supplemented with 250 mM KCl. The sucrose gradients were ultracentrifuged at 30,000 rpm in a SW41Ti swinging-bucket rotor (Beckman) at 4°C for 20 h. After centrifugation, 11 fractions were collected from top to bottom. A portion of each fraction was analyzed by SDS-PAGE followed by immunoblotting to identify the TGN/Golgi-containing fractions.

### Proximity ligation assay (PLA)

2 × 10^5^ HeLa S3 cells were seeded onto glass coverslips in a 24-well plate. For IPO7 KD experiments, cells were transfected with 10 nM siRNA for 48 h. The cells were infected with HPV16.L2F PsV (MOI ∼100) or left uninfected. At 16, 32, or 40 hpi, cells were fixed with 4% paraformaldehyde (PFA), permeabilized with 0.1% saponin, and incubated at 4°C overnight with the combination of rabbit and mouse antibodies as listed in Table 3.

**Table 3.** Combination of antibodies used in PLA.

| PLA targets | Rabbit antibody | Mouse antibody |
| --- | --- | --- |
| IPO7 and HPV16 L1 | anti-IPO7<br>(1:100 dilution) | anti-L1<br>(1:500 dilution) |
| IPO7 and HPV16 L2-3xFLAG | anti-FLAG<br>(1:200 dilution) | anti-IPO7<br>(1:100 dilution) |
| GM130 and IPO7 | anti-GM130<br>(1:200 dilution) | anti-IPO7<br>(1:100 dilution) |
| GM130 and HPV16 L2-3xFLAG | anti-GM130<br>(1:200 dilution) | anti-FLAG<br>(1:500 dilution) |
| TGN46 and HPV16 L2-3xFLAG | anti-TGN46<br>(1:200 dilution) | anti-FLAG<br>(1:500 dilution) |

PLA was performed with Duolink reagents (Millipore Sigma) as described previously (*56*). In brief, cells were incubated with PLA probes in a humidified chamber at 37°C for 1 h. Ligation was performed at 37°C for 45 min, and amplification was performed for 3 h at 37°C. Coverslips were mounted with mounting medium containing DAPI (Abcam ab104139) and visualized by confocal fluorescence microscopy (Zeiss LSM 980 or LSM 800). Images were processed by ZEN software (Zeiss) and quantitatively analyzed by Fiji software (*57*) to measure the total fluorescence intensity per cell in each sample. All PLA experiments were performed independently at least two (in most cases three) times with similar results. Data points are shown with the means and standard deviations from one representative independent experiment. A two-tailed, unequal variance *t*-test or one-way ANOVA were used to determine statistical significance.

### EdU labeling and immunofluorescence

To produce Edu-labeled HPV16 PsV, 100 μM of EdU (Thermo Fisher Scientific, C10337) was added to the HEK 293TT cell growth medium at 6 hours after transfection of p16SheLL.L2F and the Gluc reporter plasmid. PsV was purified from transfected cells as described above. HeLa cells were uninfected or infected (MOI ∼150) with EdU-labeled HPV16.L2F PsV. At 32 hpi, cells were fixed in 4% PFA and permeabilized with 1% saponin. The cells were incubated with the Click-iT reaction mixture using the Alexa Fluor 488 Imaging Kit (Thermo Fisher Scientific, C10425) according to the manufacturer’s instruction. The cells were then incubated with antibodies recognizing anti-Nesprin2 antibody and the appropriate secondary antibodies.

For immunofluorescence of GM130 and the traptamer FA or FA-JX4, 1× 10^5^ HeLa S3 or cells stably expressing FA or FA-JX4 were seeded onto glass coverslips in a 12-well for 48 h. Cells were fixed with 4% PFA for 10 min, permeabilized and blocked in DMEM supplemented with 0.1% Triton X-100 and 10% FBS for 30 min at room temperature. Samples were incubated overnight at 4°C with 1:400 mouse anti-FLAG M2 antibody and 1:200 rabbit anti-GM130 antibody in DMEM/0.1% Triton X-100/10% FBS. Alexa Fluor– conjugated secondary antibodies (Life Technologies) were 1:1000 diluted in DMEM/0.1% Triton X-100/10% FBS and incubated at room temperature for 1 h.

For immunofluorescence of IPO7 and GM130, 7× 10^4^ HeLa cells were reverse-transfected with 10 nM of Scr or *IPO7* siRNA #2 and seeded onto glass coverslips in a 12-well for 48 h. Cells were fixed with 4% PFA for 15 min, permeabilized with 0.2% Triton X-100/ TBS/3% BSA for 20 min, and blocked with 0.2% Tween-20/TBS/3% BSA for 1 hour at room temperature. Samples were incubated overnight at 4°C with 1:250 mouse anti-IPO7 antibody and 1:1000 rabbit anti-GM130 antibody in 0.2% Tween-20/TBS/3% BSA. Alexa Fluor–conjugated secondary antibodies (Life Technologies) were 1:2000 diluted in 0.2% Tween-20/TBS/3% BSA and incubated at room temperature for 1 h.

Coverslips were mounted with mounting medium containing DAPI (Abcam, ab104139) and visualized by confocal fluorescence microscopy (Zeiss LSM 800 confocal laser scanning microscope with a Plan-Apochromat 40×/1.4 oil differential interference contrast M27 objective).

### Immunoprecipitation

For Golgi membrane IP, cell fraction #3 collected from the sucrose gradient was mixed with an anti-GM130 antibody (∼1 μg/μL) or a non-specific rabbit normal IgG antibody (∼1 μg/μL) and incubated at 4°C overnight. Where indicated, the overnight incubation was supplemented with 1% Triton X-100. Protein G-coated magnetic beads (Thermo Fisher Scientific, 10003D) were pre-equilibrated with low-DTT HBS buffer [10 mM HEPES pH 7.2, 1 mM Mg(OAc)_2_, 0.25 M sucrose, 1 mM EDTA, 0.1 mM DTT, 1 mM PMSF] and used to capture the immune complex at 4°C for 30 min. After incubation, the beads were washed on ice once with low-DTT HBS buffer, three times with low-DTT HBS buffer supplemented with 300 mM NaCl, and once with low-DTT HBS buffer. The samples were eluted with 1X SDS sample buffer containing 2-mercaptoethanol at 95°C for 10 min. As a control experiment, a rat anti-BAP31 antibody (∼1 μg/μL) was used to isolate the ER membrane, with a rat anti-HA antibody used for the control IP.

For IP of IPO7 from the whole cell extract, cells were grown to ∼80% confluency in 10-cm plates and infected with HPV16.L2F PsV (MOI ∼1) or left uninfected. At the indicated time after infection, cells from one 10-cm plate were washed with PBS three times and lysed in 400 μL of 1% Triton X-100 in HN buffer and then incubated on ice for 10 min. After centrifugation at 16,100 *g* at 4°C for 10 min, the resulting supernatant was incubated with anti-IPO7 antibodies (∼25 μg/mL) at 4°C overnight. The immune complex was captured with protein G-coated magnetic beads (Thermo Fisher Scientific, 10003D) at 4°C for 30 min. Beads were washed on ice once with 1% Triton X-100 in HN buffer, followed by one wash with 1% Triton X-100 in high-salt HN buffer (50 mM HEPES, 500 mM NaCl, 1 mM PMSF), two washes with 1% Triton X-100 in HN buffer, and elution in 1X SDS sample buffer containing 2-mercaptoethanol at 95°C for 10 min. For IPO7 IP experiments following *COPA/G1* knockdown, cells were treated with 2 nM of Scr siRNA or a mixture of 1 nM *COPA* and 1 nM *COPG1* siRNAs for 24 h before infection. γ-COP IP experiments were performed as previously described (*21*), except cells were transfected with 10 nM of Scr or *IPO7* siRNA #2 for 48 h before infection.

### Biotin-peptide pull-down assay

Peptides consisting of the HPV16 L2 sequence were synthesized with an N-terminal biotin tag (GenScript). Peptides were dissolved in DMSO or water according to manufacturer’s instruction, and peptide stocks (5 mg/mL) were aliquoted and stored at −80°C. 1 × 10^6^ HeLa cells were plated in 6-cm dishes. 24 h later, cells were collected by trypsinization, washed with PBS, and lysed with 165 μL HEPES buffer (1% Triton X-100, 20 mM HEPES pH 7.4, 50 mM NaCl, 5 mM MgCl_2_, 1 mM DTT, 1 mM PMSF). The lysate was incubated on ice for 45 min followed by centrifugation at 14,000 *g* at 4°C for 15 min. The resulting supernatant was incubated with 10 μg of indicated biotinylated peptide or an equivalent volume (2 μL) of DMSO as a negative control at 4°C for 2 h. Streptavidin magnetic beads (Pierce, 88817) pre-equilibrated with HEPES buffer were added to the lysates and incubated at 4°C for 1 h. The beads were captured and washed on ice three times with HEPES buffer and incubated at 95°C for 10 min in 1X SDS sample buffer containing 2-mercaptoethanol. For the biotin-peptide pull-down assay using purified FLAG-IPO7, the same conditions were used except ∼100 ng FLAG-IPO7 or ∼80 ng EGFP-FLAG instead of cellular extracts was incubated with 10 μg of indicated biotinylated peptide and captured with streptavidin magnetic beads.

### Preparation of recombinant proteins

Two 10-cm plates of HEK 293TT cells transfected with an expression plasmid for 48 h were harvested and lysed in 1.2 mL of HN buffer containing 1% Triton X-100 on ice for 20 min. Following centrifugation at 16,100 *g* at 4°C for 10 min, the resulting supernatant was incubated with anti-FLAG M2 antibody-conjugated agarose beads (Millipore Sigma, A2220) at 4°C for 2 h. The beads were recovered by centrifugation and washed on ice twice with HN buffer containing 1% Triton X-100 supplemented with 300 mM NaCl and 1 mM ATP, once with HN buffer containing 1% Triton X-100, and then once with HN buffer containing 0.1% Triton X-100. The recombinant proteins were eluted twice with 150 μg/mL 3xFLAG peptide (Millipore Sigma) in HN buffer containing 0.1% Triton X-100 at room temperature for 30 min. A portion of the sample was analyzed with BSA as the standard by SDS-PAGE followed by SimplyBlue SafeStain (Invitrogen) to assess the quality and quantity of the purified recombinant proteins.

### Statistical Analysis

#### Quantitation of HPV infectivity by flow cytometry

Flow cytometry data was processed by Everest software (Bio-Rad). After gating for size and singlets, the population of DAPI-negative cells (∼1 × 10^4^ cells per condition) was analyzed for GFP or mCherry fluorescence. Data are presented as the means and standard deviations of three independent experiments. A two-tailed, unequal variance *t*-test was used to determine statistical significance.

#### Quantitation of cell cycle by flow cytometry

DNA content was measured by Hoechst 33342 staining. The percentage of cells in G1, S, or G2/M phases was determined from control cell cultures using FlowJo software (BD). Data from three independent experiments are presented as the percentage of cells in each phase. A two-tailed, unequal variance *t*-test was used to determine statistical significance between the knockdown and control cells for each cell cycle phase.

#### Quantitation of PLA and Edu images

Images were analyzed by Fiji software (*57*) to measure the total fluorescence intensity per cell in each sample. Approximately 80 or more cells from 10 representative images were quantified for each condition. All PLA and EdU-staining experiments were performed independently at least two times with similar results. Because it is difficult to directly compare the quantitative results between experiments, data points are shown with the means and standard deviations from one representative independent experiment. A two-tailed, unequal variance *t*-test or one-way ANOVA were used to determine statistical significance. Fluorescence intensity of nuclear, extranuclear, and total were analyzed per cell in each sample.

## Acknowledgments

We would like to thank Pierre Coulombe (University of Michigan) for sharing the Zeiss LSM 800 confocal microscope. We also thank members of the Tsai lab, the DiMaio lab, and the Chelsey Spriggs lab (University of Michigan) for helpful discussion.

## Author contributions

Conceptualization: TTW, BT

Methodology: TTW, YT, MCH

Investigation: TTW, YT, MCH, ETH

Supervision: BT, DD

Writing—original draft: TTW, BT

Writing—review & editing: TTW, YT, MCH, ETH, DD, BT

## Data and materials availability

All data are available in the main text or the supplementary materials.

---

**Figure S1. (related to Fig. 1).**
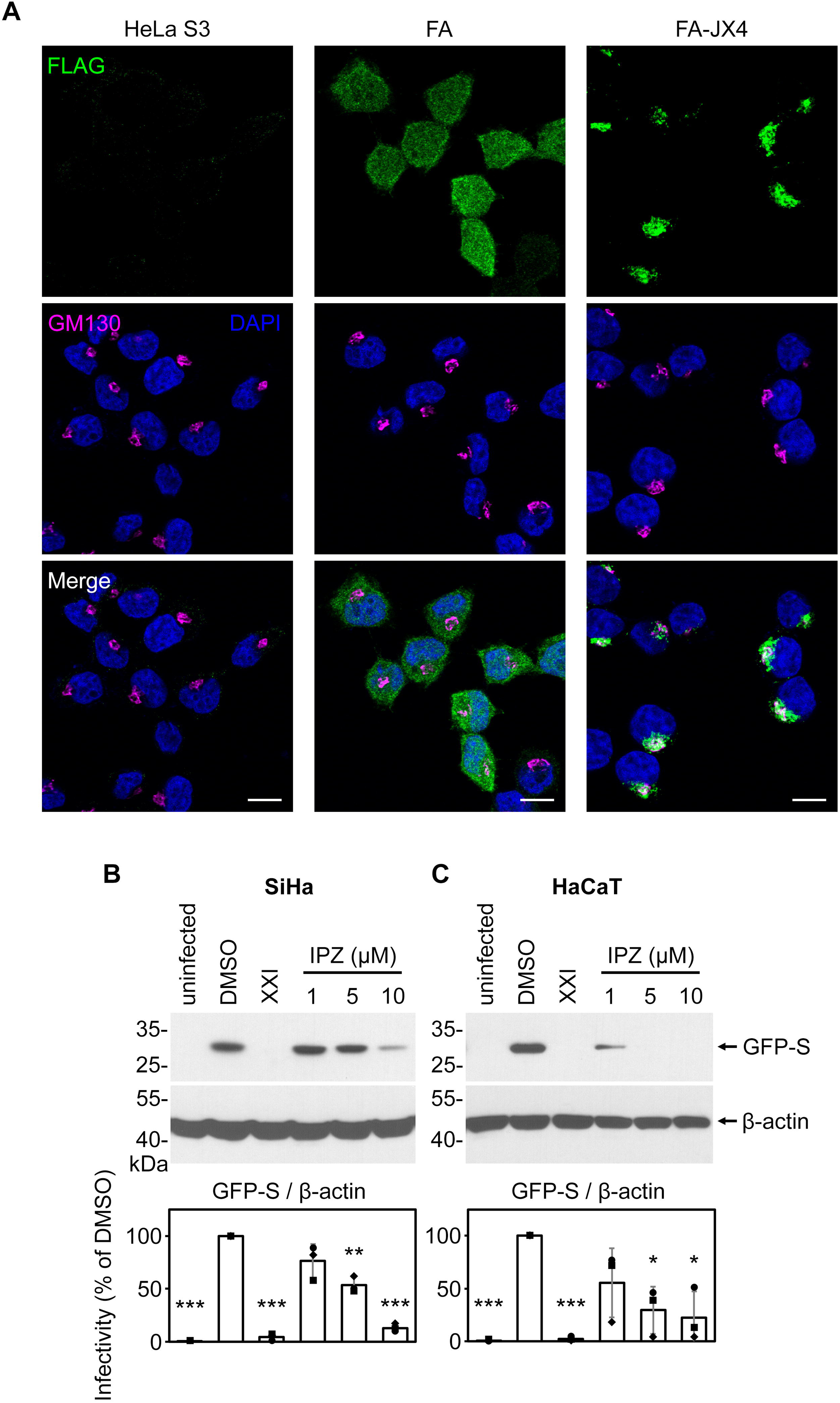
Study of a Golgi-localized traptamer reveals an important role of importin β members in HPV infection. **(A)** HeLa S3 or cells stably expressing traptamer FA-JX4 or the control FA were fixed, permeabilized, and subjected to immunofluorescent staining using antibodies recognizing FLAG (green) and the *cis*-Golgi protein GM130 (magenta), where colocalization is pseudo-colored in white in the merged image. Nuclei were stained with DAPI (blue). Scale bar, 10 µm. **(B)** The importin β inhibitor blocks HPV infection in SiHa cells. SiHa cells were uninfected or infected with HPV16.L2F PsV (MOI ∼0.3) containing a GFP reporter plasmid. Importazole (IPZ) at the indicated concentration (1–10 µM), 1 µM of the γ-secretase inhibitor (XXI), or the solvent dimethyl sulfoxide (DMSO) was added to cells at the time of infection. At 48 hours post-infection (hpi), cells were lysed and the resulting whole cell extract was subjected to SDS-PAGE followed by immunoblotting with antibodies recognizing GFP or β-actin as a loading control; representative immunoblots are shown in the upper panels. The results were quantified (lower panel), in which the intensity of GFP-S was normalized to that of β-actin in each sample. The infectivity in cells treated with DMSO was used for normalization. The means and standard deviations of three independent experiments with individual data points are shown. A two-tailed, unequal variance *t*-test was used to determine statistical significance when compared to DMSO-treated cells infected with HPV16.L2F PsV. \**P* < 0.05; \*\**P* < 0.01; \*\*\**P* < 0.001. **(C)** As in (B), except HaCaT cells were analyzed.

**Figure S2. (related to Fig. 2).**
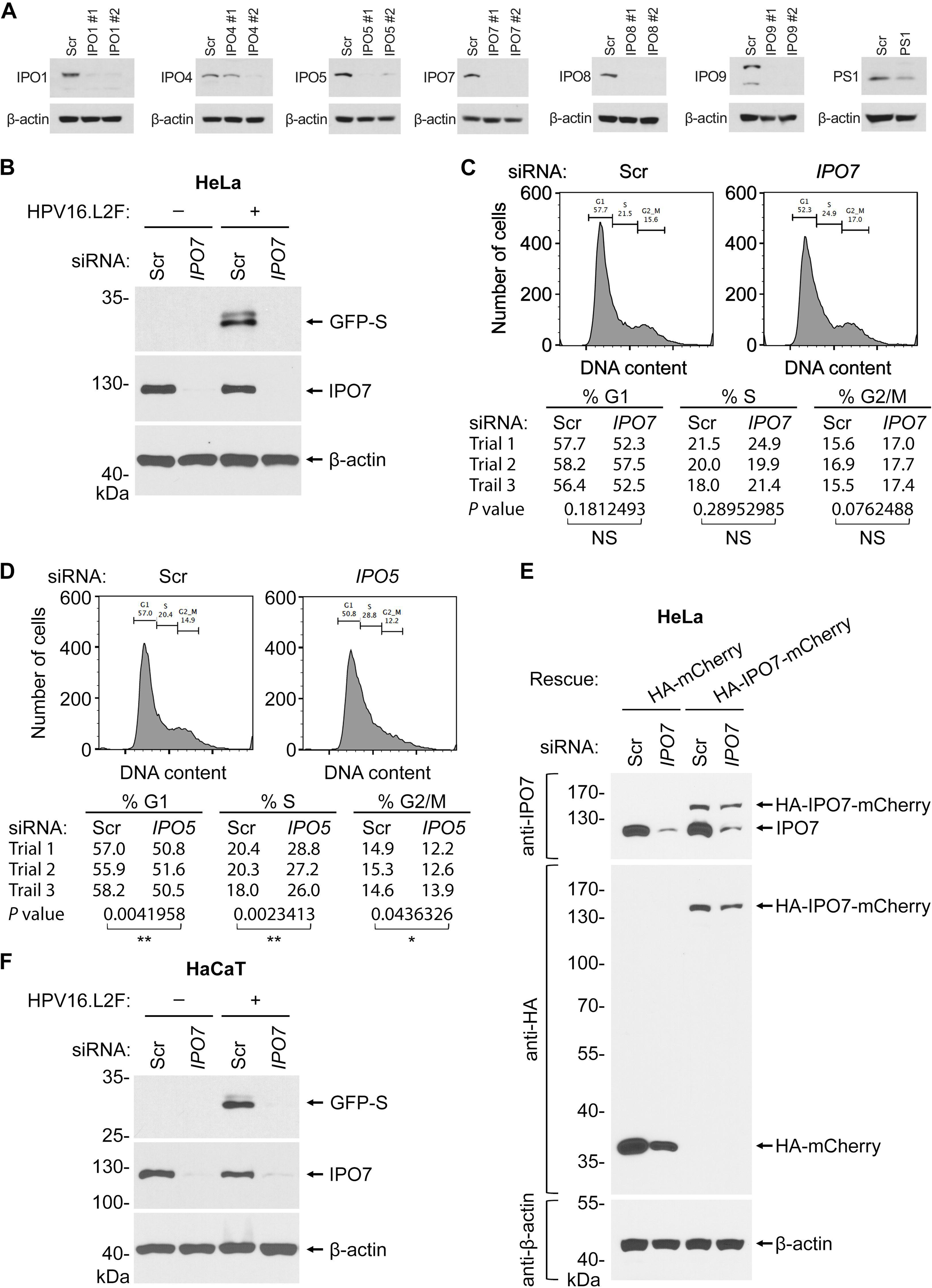
IPO7 promotes HPV infection. **(A)** HeLa cells transfected with 10 nM of the control scramble (Scr) or the indicated siRNAs for 48 h were harvested to verify the protein levels of the intended targets. The resulting cell extracts were subjected to SDS-PAGE analyses followed by immunoblotting using the indicated antibodies. **(B)** HeLa cells transfected with 10 nM Scr or *IPO7* siRNA #2 were uninfected or infected with HPV16.L2F PsV (MOI ∼0.3) containing a GFP reporter plasmid. At 48 hpi, cells were harvested, and the resulting cell extracts were subjected to SDS-PAGE analyses followed by immunoblotting using antibodies recognizing GFP, IPO7, or β-actin. **(C)** HeLa cells were transfected with 10 nM Scr or *IPO7* siRNA #2 for 48 h and incubated with the cell membrane-permeable DNA dye Hoechst 33342 for 30 min and analyzed by flow cytometry to determine the relative Hoechst 33342 fluorescence as a measure of DNA content. The percentage of cells in each phase of the cell cycle is indicated. The representative histograms of three independent experiments are shown, with cell cycle phase distributions summarized below. Statistical significance for each phase was determined by a two-tailed, unequal variance *t*-test. NS, not significant (*P* > 0.05). **(D)** As in (C), except cells transfected with 10 nM Scr or *IPO5* siRNA #2 were analyzed. *, *P* < 0.05; **, *P* < 0.01. **(E)** HeLa cells were transfected with indicated DNA constructs (rescue) for 24 h to express HA-IPO7-mCherry or the control HA-mCherry, followed by another transfection with 10 nM Scr or *IPO7* siRNA #2 for 48 h. Cells were lysed, and the resulting whole cell extract was subjected to SDS-PAGE followed by immunoblotting with the indicated antibodies. **(F)** As in (B), except HaCaT cells were analyzed.

**Figure S3. (related to Fig. 3).**
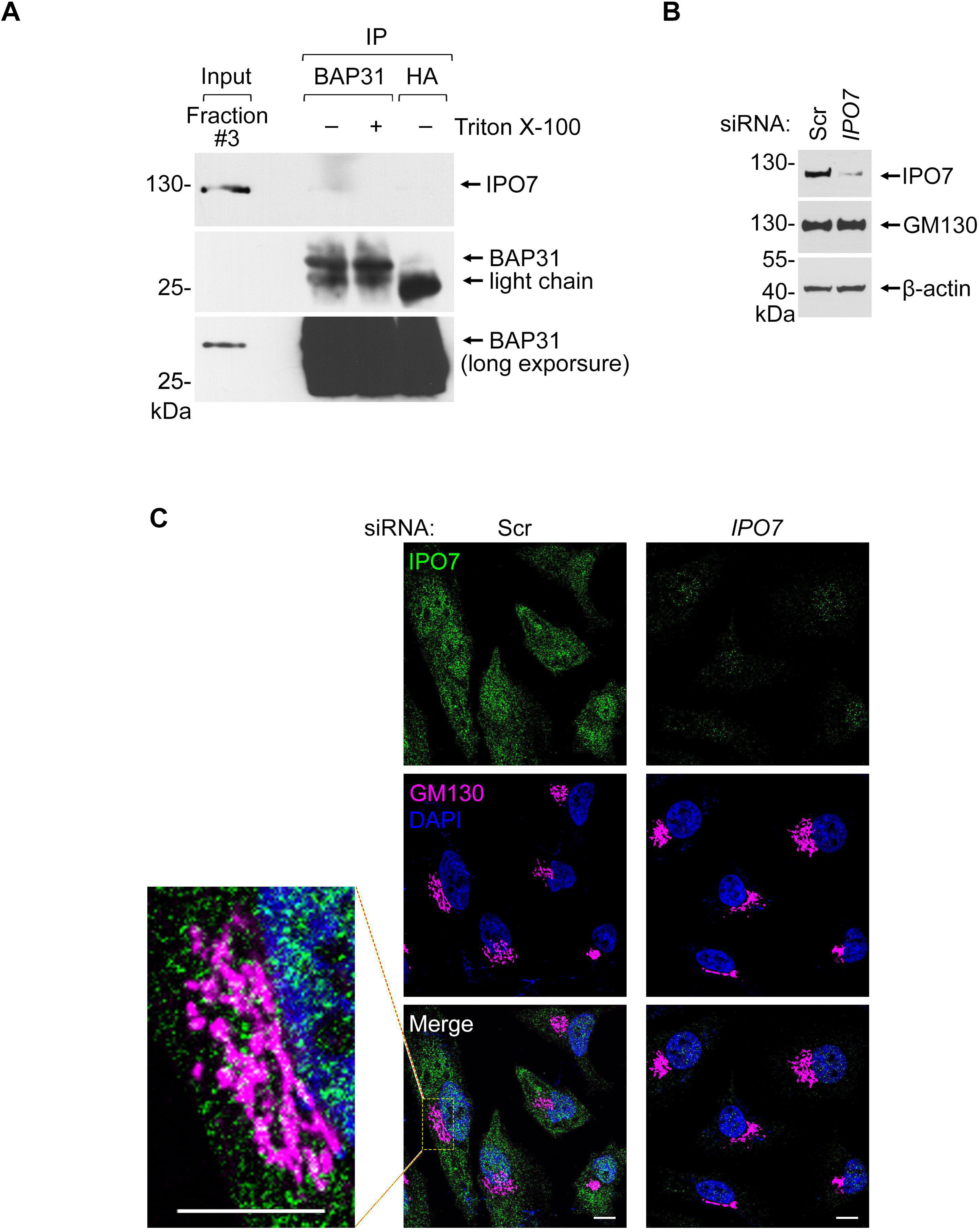
IPO7 associates with the Golgi membrane. **(A)** Fraction #3 collected from the cell fractionation via sucrose gradient (Fig. 3A) was subjected to immunoprecipitation with an anti-BAP31 antibody or an anti-HA antibody as a control, in the absence or presence of 1% Triton X-100. The immunoprecipitated materials were immunoblotted with the indicated antibodies. **(B)** HeLa S3 cells were transfected with 10 nM Scr or *IPO7* siRNA #2 for two days and harvested. The resulting cell extracts were subjected to SDS-PAGE analyses followed by immunoblotting using antibodies recognizing IPO7, GM130, or β-actin (as a loading control). **(C)** HeLa cells were transfected with 10 nM Scr or *IPO7* siRNA #2 for two days, fixed and permeabilized, and subjected to immunofluorescent staining using antibodies recognizing IPO7 (green) and GM130 (magenta), where colocalization is pseudo-colored in white in the merged image. Nuclei were stained with DAPI (blue). Scale bar, 10 μm.

