## Supplemental Figures S1 to S3 and Table S1 for "Golgi-associated nuclear import receptor importin-7 targets HPV from the Golgi to the nucleus to promote infection"


**This PDF file includes:**

**Figs. S1 to S3**
**Table S1**

**Fig. S1 (related to Fig. 1) Study of a Golgi-localized traptamer reveals an important role of importin β members in HPV infection**

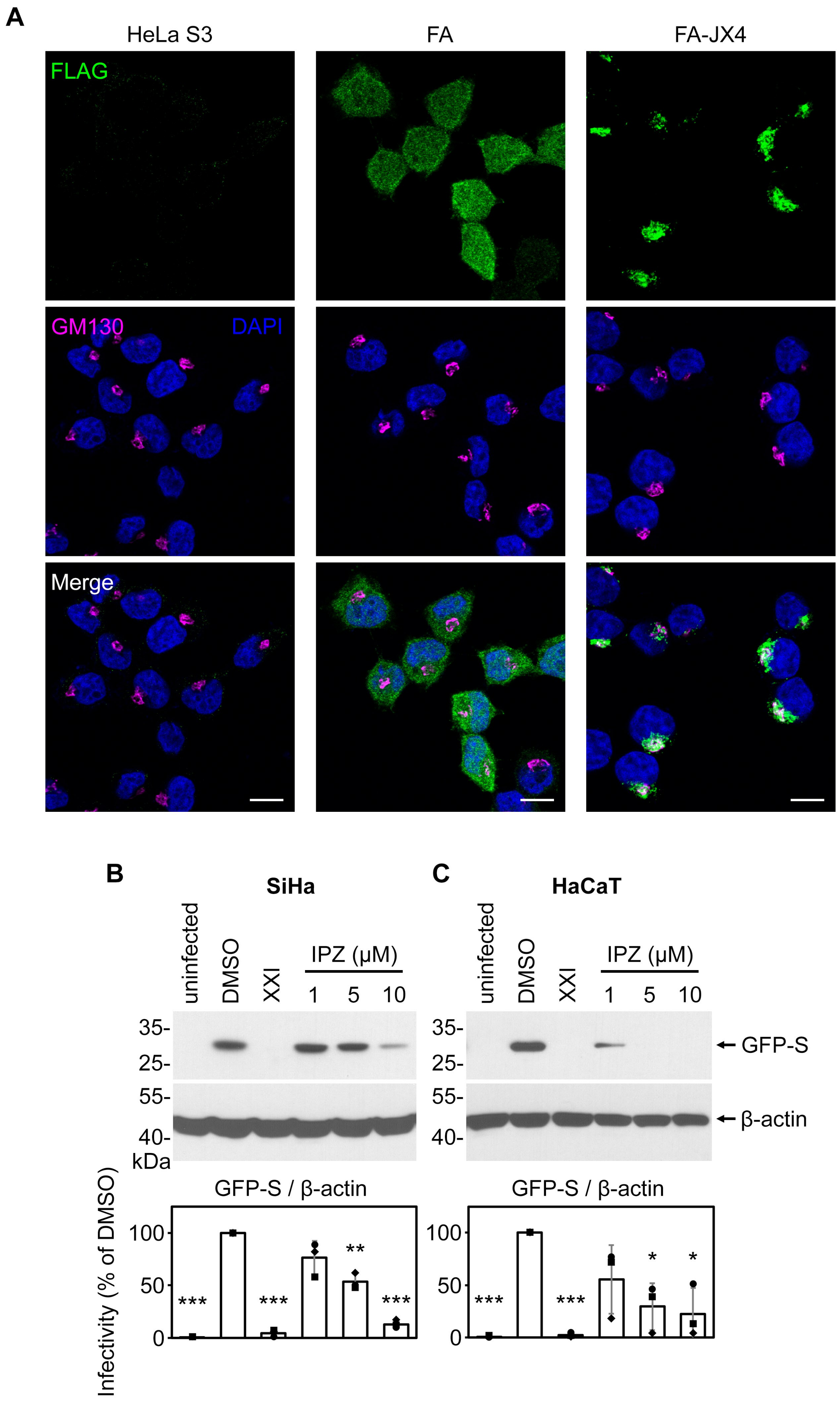

**Figure S1. (related to Fig. 1) Study of a Golgi-localized traptamer reveals an important role of importin β members in HPV infection**

**(A)** HeLa S3 or cells stably expressing traptamer FA-JX4 or the control FA were fixed, permeabilized, and subjected to immunofluorescent staining using antibodies recognizing FLAG (green) and the *cis*-Golgi protein GM130 (magenta), where colocalization is pseudo-colored in white in the merged image. Nuclei were stained with DAPI (blue). Scale bar, 10 μm.

**(B)** The importin β inhibitor blocks HPV infection in SiHa cells. SiHa cells were uninfected or infected with HPV16.L2F PsV (MOI ~0.3) containing a GFP reporter plasmid. Importazole (IPZ) at the indicated concentration (1–10 μM), 1 μM of the γ-secretase inhibitor (XXI), or the solvent dimethyl sulfoxide (DMSO) was added to cells at the time of infection. At 48 hours post-infection (hpi), cells were lysed and the resulting whole cell extract was subjected to SDS-PAGE followed by immunoblotting with antibodies recognizing GFP or β-actin as a loading control; representative immunoblots are shown in the upper panels. The results were quantified (lower panel), in which the intensity of GFP-S was normalized to that of β-actin in each sample. The infectivity in cells treated with DMSO was used for normalization. The means and standard deviations of three independent experiments with individual data points are shown. A two-tailed, unequal variance *t*-test was used to determine statistical significance when compared to DMSO-treated cells infected with HPV16.L2F PsV. *$P < 0.05$; **$P < 0.01$; ***$P < 0.001$.

**(C)** As in (B), except HaCaT cells were analyzed.

**Fig. S2 (related to Fig. 2) IPO7 promotes HPV infection**

**A**

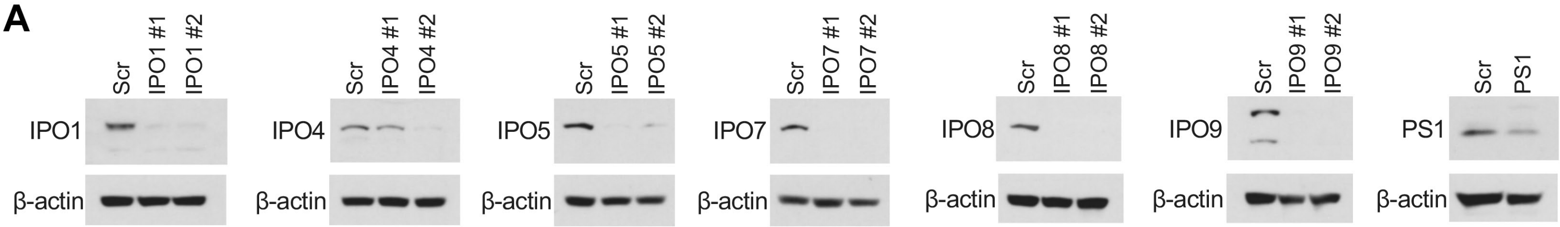


**B**

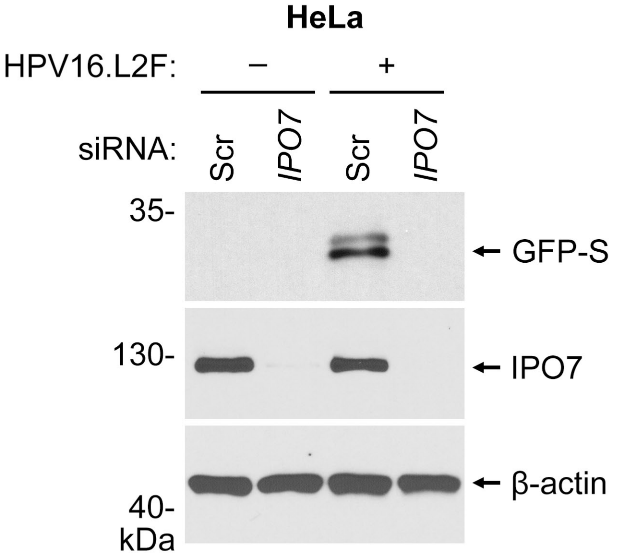


**C**

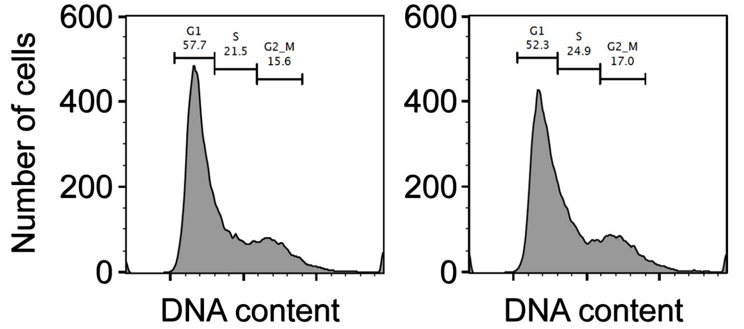


| | % G1 | | % S | | % G2/M | |
|---|---|---|---|---|---|---|
| siRNA: | Scr | *IPO7* | Scr | *IPO7* | Scr | *IPO7* |
| Trial 1 | 57.7 | 52.3 | 21.5 | 24.9 | 15.6 | 17.0 |
| Trial 2 | 58.2 | 57.5 | 20.0 | 19.9 | 16.9 | 17.7 |
| Trail 3 | 56.4 | 52.5 | 18.0 | 21.4 | 15.5 | 17.4 |
| *P* value | 0.1812493 | | 0.28952985 | | 0.0762488 | |
| | NS | | NS | | NS | |

**D**

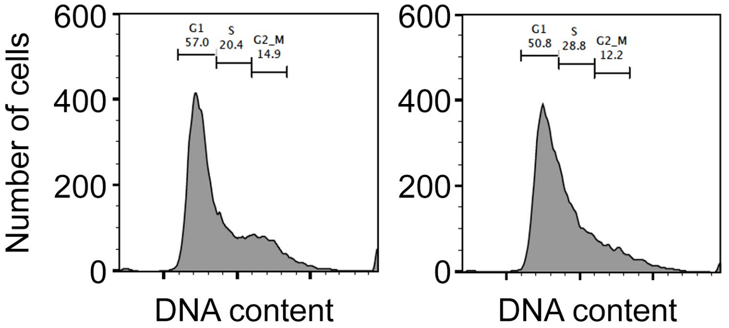


| | % G1 | | % S | | % G2/M | |
|---|---|---|---|---|---|---|
| siRNA: | Scr | *IPO5* | Scr | *IPO5* | Scr | *IPO5* |
| Trial 1 | 57.0 | 50.8 | 20.4 | 28.8 | 14.9 | 12.2 |
| Trial 2 | 55.9 | 51.6 | 20.3 | 27.2 | 15.3 | 12.6 |
| Trail 3 | 58.2 | 50.5 | 18.0 | 26.0 | 14.6 | 13.9 |
| *P* value | 0.0041958 | | 0.0023413 | | 0.0436326 | |
| | ** | | ** | | * | |

**E**

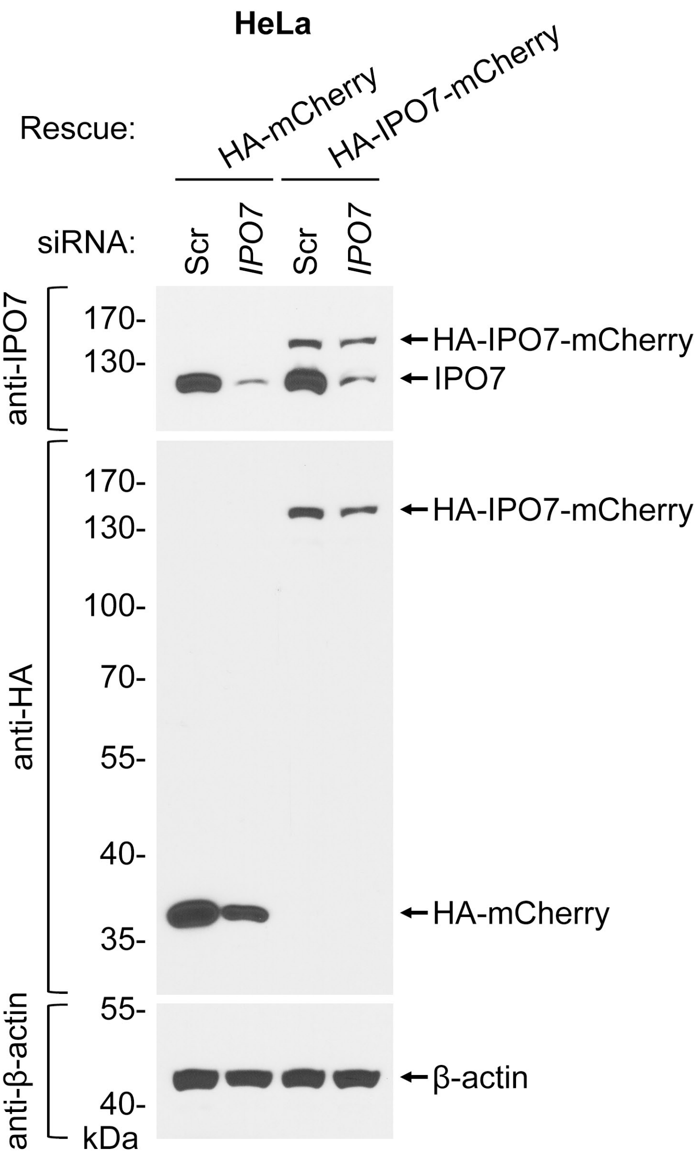


**F**

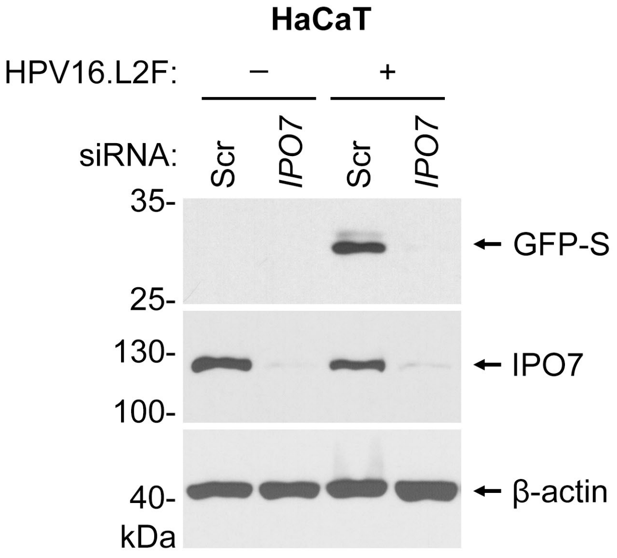

**Figure S2. (related to Fig. 2) IPO7 promotes HPV infection**

- **(A)** HeLa cells transfected with 10 nM of the control scramble (Scr) or the indicated siRNAs for 48 h were harvested to verify the protein levels of the intended targets. The resulting cell extracts were subjected to SDS-PAGE analyses followed by immunoblotting using the indicated antibodies.
- **(B)** HeLa cells transfected with 10 nM Scr or *IPO7* siRNA #2 were uninfected or infected with HPV16.L2F PsV (MOI ~0.3) containing a GFP reporter plasmid. At 48 hpi, cells were harvested, and the resulting cell extracts were subjected to SDS-PAGE analyses followed by immunoblotting using antibodies recognizing GFP, IPO7, or β-actin.
- **(C)** HeLa cells were transfected with 10 nM Scr or *IPO7* siRNA #2 for 48 h and incubated with the cell membrane-permeable DNA dye Hoechst 33342 for 30 min and analyzed by flow cytometry to determine the relative Hoechst 33342 fluorescence as a measure of DNA content. The percentage of cells in each phase of the cell cycle is indicated. The representative histograms of three independent experiments are shown, with cell cycle phase distributions summarized below. Statistical significance for each phase was determined by a two-tailed, unequal variance *t*-test. NS, not significant ($P > 0.05$).
- **(D)** As in (C), except cells transfected with 10 nM Scr or *IPO5* siRNA #2 were analyzed. *, $P < 0.05$; **, $P < 0.01$.
- **(A)** HeLa cells were transfected with indicated DNA constructs (rescue) for 24 h to express HA-IPO7-mCherry or the control HA-mCherry, followed by another transfection with 10 nM Scr or *IPO7* siRNA #2 for 48 h. Cells were lysed, and the resulting whole cell extract was subjected to SDS-PAGE followed by immunoblotting with the indicated antibodies.
- **(B)** As in (B), except HaCaT cells were analyzed.

**Fig. S3 (related to Fig. 3) IPO7 associates with the Golgi membrane**

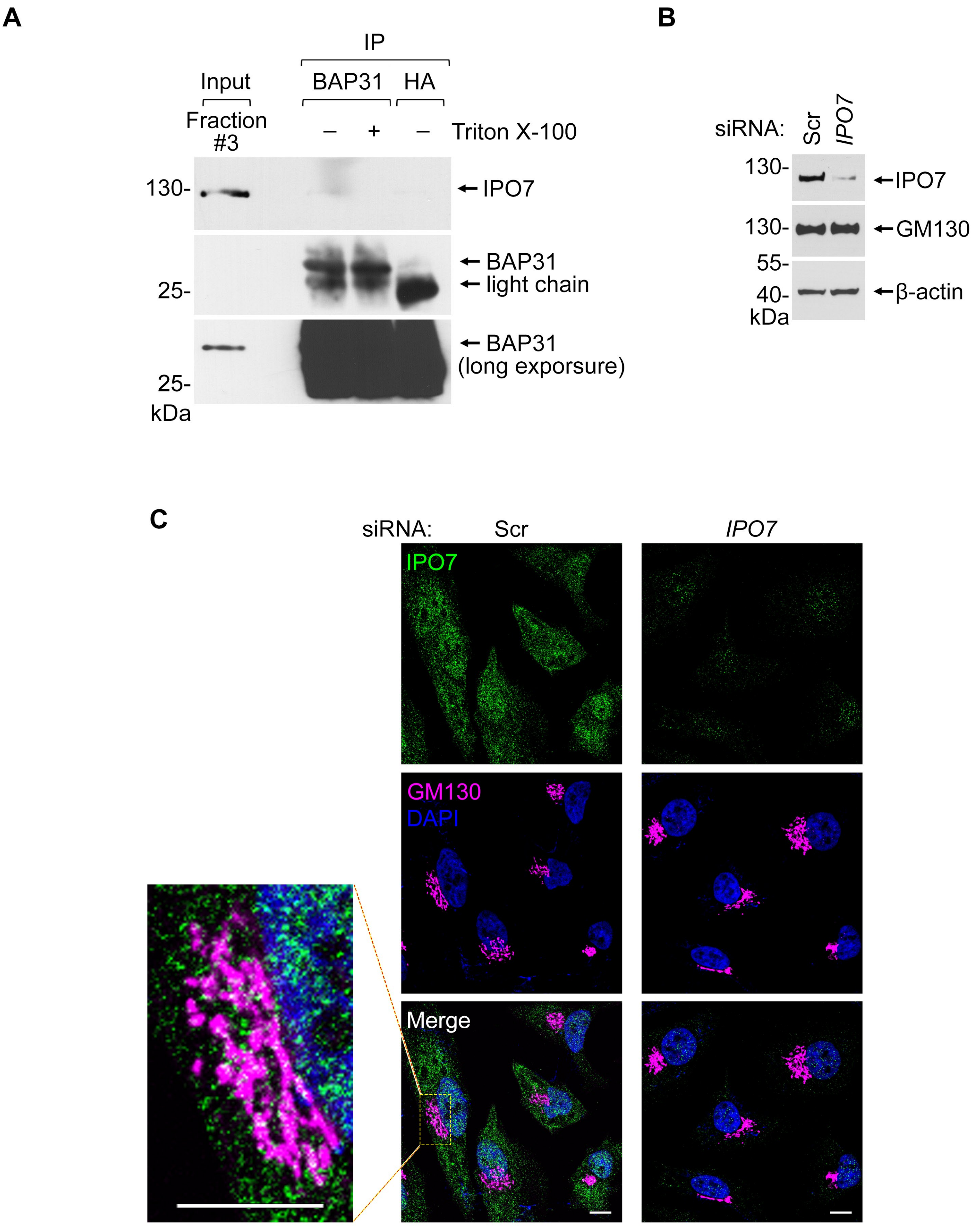

**Figure S3. (related to Fig. 3) IPO7 associates with the Golgi membrane**

- **(A)** Fraction #3 collected from the cell fractionation via sucrose gradient (Fig. 3A) was subjected to immunoprecipitation with an anti-BAP31 antibody or an anti-HA antibody as a control, in the absence or presence of 1% Triton X-100. The immunoprecipitated materials were immunoblotted with the indicated antibodies.
- **(B)** HeLa S3 cells were transfected with 10 nM Scr or *IPO7* siRNA #2 for two days and harvested. The resulting cell extracts were subjected to SDS-PAGE analyses followed by immunoblotting using antibodies recognizing IPO7, GM130, or β-actin (as a loading control).
- **(C)** HeLa cells were transfected with 10 nM Scr or *IPO7* siRNA #2 for two days, fixed and permeabilized, and subjected to immunofluorescent staining using antibodies recognizing IPO7 (green) and GM130 (magenta), where colocalization is pseudo-colored in white in the merged image. Nuclei were stained with DAPI (blue). Scale bar, 10 μm.

### Table S1. Mass-spectrometry results

| Reference | Gene Symbol | Annotation | MWT(kDa) | Unique FA | Total FA | Unique JX4-FA | Total JX4-FA | Sum Intensity FA | Sum Intensity JX4-FA |
|---|---|---|---|---|---|---|---|---|---|
| VL1_HPV16 | HPV16 L1 | additional search for L1 | 56.24 | 0 | 0 | 0 | 0 | | |
| VL2_HPV16 | HPV16 L2 | additional search for L2 | 50.66 | 0 | 0 | 0 | 0 | | |
| sp \| P78527 \| PRKDC_HUMAN | PRKDC | DNA-dependent protein kinase catalytic subunit OS=Homo sapiens GN=PRKDC PE=1 SV=3 | 468.79 | 0 | 0 | 75 | 127 | NF | 1.10E+07 |
| sp \| Q92616 \| GCN1_HUMAN | GCN1 | eIF-2-alpha kinase activator GCN1 OS=Homo sapiens GN=GCN1 PE=1 SV=6 | 292.57 | 0 | 0 | 52 | 106 | NF | 1.20E+07 |
| sp \| Q969Z0 \| TBRG4_HUMAN | TBRG4 | Protein TBRG4 OS=Homo sapiens GN=TBRG4 PE=1 SV=1 | 70.69 | 0 | 0 | 18 | 56 | NF | 1.50E+07 |
| sp \| P42345 \| MTOR_HUMAN | MTOR | Serine/threonine-protein kinase mTOR OS=Homo sapiens GN=MTOR PE=1 SV=1 | 288.71 | 0 | 0 | 25 | 32 | NF | 1.30E+06 |
| sp \| P06576 \| ATPB_HUMAN | ATP5B | ATP synthase subunit beta, mitochondrial OS=Homo sapiens GN=ATP5B PE=1 SV=3 | 56.52 | 0 | 0 | 17 | 32 | NF | 3.20E+06 |
| sp \| O14975 \| S27A2_HUMAN | SLC27A2 | Very long-chain acyl-CoA synthetase OS=Homo sapiens GN=SLC27A2 PE=1 SV=2 | 70.27 | 0 | 0 | 16 | 32 | NF | 2.90E+06 |
| sp \| P25705 \| ATPA_HUMAN | ATP5A1 | ATP synthase subunit alpha, mitochondrial OS=Homo sapiens GN=ATP5A1 PE=1 SV=1 | 59.71 | 0 | 0 | 15 | 26 | NF | 2.70E+06 |
| sp \| P08195 \| 4F2_HUMAN | SLC3A2 | 4F2 cell-surface antigen heavy chain OS=Homo sapiens GN=SLC3A2 PE=1 SV=3 | 67.95 | 0 | 0 | 14 | 24 | NF | 2.70E+06 |
| sp \| Q12905 \| ILF2_HUMAN | ILF2 | Interleukin enhancer-binding factor 2 OS=Homo sapiens GN=ILF2 PE=1 SV=2 | 43.04 | 0 | 0 | 13 | 24 | NF | 2.00E+06 |
| sp \| Q9BVA1 \| TBB2B_HUMAN | TUBB2B | Tubulin beta-2B chain OS=Homo sapiens GN=TUBB2B PE=1 SV=1 | 49.92 | 0 | 0 | 2 | 22 | NF | 6.80E+06 |
| sp \| O14980 \| XPO1_HUMAN | XPO1 | Exportin-1 OS=Homo sapiens GN=XPO1 PE=1 SV=1 | 123.31 | 0 | 0 | 12 | 20 | NF | 7.40E+05 |
| sp \| Q8WUX1 \| S38A5_HUMAN | SLC38A5 | Sodium-coupled neutral amino acid transporter 5 OS=Homo sapiens GN=SLC38A5 PE=1 SV=1 | 51.42 | 0 | 0 | 5 | 20 | NF | 1.00E+07 |
| sp \| Q3MII6 \| TBC25_HUMAN | TBC1D25 | TBC1 domain family member 25 OS=Homo sapiens GN=TBC1D25 PE=1 SV=2 | 76.28 | 0 | 0 | 11 | 19 | NF | 1.40E+06 |
| sp \| Q13535 \| ATR_HUMAN | ATR | Serine/threonine-protein kinase ATR OS=Homo sapiens GN=ATR PE=1 SV=3 | 301.17 | 0 | 0 | 14 | 16 | NF | 4.50E+05 |
| sp \| Q2VPB7 \| AP5B1_HUMAN | AP5B1 | AP-5 complex subunit beta-1 OS=Homo sapiens GN=AP5B1 PE=1 SV=4 | 93.89 | 0 | 0 | 10 | 16 | NF | 4.30E+06 |
| sp \| Q86XA9 \| HTR5A_HUMAN | HEATR5A | HEAT repeat-containing protein 5A OS=Homo sapiens GN=HEATR5A PE=1 SV=2 | 221.86 | 0 | 0 | 12 | 15 | NF | 4.10E+05 |
| sp \| Q99720 \| SGMR1_HUMAN | SIGMAR1 | Sigma non-opioid intracellular receptor 1 OS=Homo sapiens GN=SIGMAR1 PE=1 SV=1 | 25.11 | 0 | 0 | 5 | 15 | NF | 1.80E+06 |
| sp \| Q13395 \| TARB1_HUMAN | TARBP1 | Probable methyltransferase TARBP1 OS=Homo sapiens GN=TARBP1 PE=1 SV=1 | 181.56 | 0 | 0 | 12 | 14 | NF | 2.80E+05 |
| sp \| Q01813 \| PFKAP_HUMAN | PFKP | ATP-dependent 6-phosphofructokinase, platelet type OS=Homo sapiens GN=PFKP PE=1 SV=2 | 85.54 | 0 | 0 | 8 | 13 | NF | 4.00E+05 |
| sp \| Q9BUF5 \| TBB6_HUMAN | TUBB6 | Tubulin beta-6 chain OS=Homo sapiens GN=TUBB6 PE=1 SV=1 | 49.82 | 0 | 0 | 7 | 13 | NF | 2.30E+06 |
| sp \| P55060 \| XPO2_HUMAN | CSE1L | Exportin-2 OS=Homo sapiens GN=CSE1L PE=1 SV=3 | 110.35 | 0 | 0 | 9 | 12 | NF | 3.30E+05 |
| sp \| Q9Y6D5 \| BIG2_HUMAN | ARFGEF2 | Brefeldin A-inhibited guanine nucleotide-exchange protein 2 OS=Homo sapiens GN=ARFGEF2 PE=1 SV=3 | 201.91 | 0 | 0 | 11 | 12 | NF | 1.50E+05 |
| sp \| P27708 \| PYR1_HUMAN | CAD | CAD protein OS=Homo sapiens GN=CAD PE=1 SV=3 | 242.83 | 0 | 0 | 10 | 12 | NF | 1.20E+06 |
| sp \| O43592 \| XPOT_HUMAN | XPOT | Exportin-T OS=Homo sapiens GN=XPOT PE=1 SV=2 | 109.89 | 0 | 0 | 9 | 12 | NF | 4.70E+05 |
| sp \| P53618 \| COPB_HUMAN | COPB1 | Coatomer subunit beta OS=Homo sapiens GN=COPB1 PE=1 SV=3 | 107.07 | 0 | 0 | 7 | 12 | NF | 3.00E+05 |
| sp \| Q9H5Q4 \| TFB2M_HUMAN | TFB2M | Dimethyladenosine transferase 2, mitochondrial OS=Homo sapiens GN=TFB2M PE=1 SV=1 | 45.32 | 0 | 0 | 9 | 11 | NF | 6.60E+05 |
| sp \| P04843 \| RPN1_HUMAN | RPN1 | Dolichyl-diphosphooligosaccharide--protein glycosyltransferase subunit 1 OS=Homo sapiens GN=RPN1 PE=1 SV=1 | 68.53 | 0 | 0 | 8 | 11 | NF | 4.50E+05 |
| sp \| Q9C0E2 \| XPO4_HUMAN | XPO4 | Exportin-4 OS=Homo sapiens GN=XPO4 PE=1 SV=2 | 130.06 | 0 | 0 | 7 | 11 | NF | 4.00E+05 |
| sp \| P48960 \| CD97_HUMAN | CD97 | CD97 antigen OS=Homo sapiens GN=CD97 PE=1 SV=4 | 91.81 | 0 | 0 | 8 | 10 | NF | 2.70E+05 |
| sp \| Q9HAV4 \| XPO5_HUMAN | XPO5 | Exportin-5 OS=Homo sapiens GN=XPO5 PE=1 SV=1 | 136.22 | 0 | 0 | 8 | 10 | NF | 1.70E+05 |
| sp \| Q8TEX9 \| IPO4_HUMAN | IPO4 | Importin-4 OS=Homo sapiens GN=IPO4 PE=1 SV=2 | 118.64 | 0 | 0 | 8 | 10 | NF | 3.80E+05 |
| sp \| Q9BQG0 \| MBB1A_HUMAN | MYBBP1A | Myb-binding protein 1A OS=Homo sapiens GN=MYBBP1A PE=1 SV=2 | 148.76 | 0 | 0 | 7 | 10 | NF | 1.50E+05 |
| sp \| Q9UPY5 \| XCT_HUMAN | SLC7A11 | Cystine/glutamate transporter OS=Homo sapiens GN=SLC7A11 PE=1 SV=1 | 55.39 | 0 | 0 | 3 | 10 | NF | 6.20E+05 |
| sp \| Q96JI7 \| SPTCS_HUMAN | SPG11 | Spatacsin OS=Homo sapiens GN=SPG11 PE=1 SV=3 | 278.69 | 0 | 0 | 8 | 9 | NF | 1.60E+05 |
| sp \| Q9P035 \| HACD3_HUMAN | HACD3 | Very-long-chain (3R)-3-hydroxyacyl-CoA dehydratase 3 OS=Homo sapiens GN=HACD3 PE=1 SV=2 | 43.13 | 0 | 0 | 4 | 9 | NF | 1.30E+06 |
| sp \| P04844 \| RPN2_HUMAN | RPN2 | Dolichyl-diphosphooligosaccharide--protein glycosyltransferase subunit 2 OS=Homo sapiens GN=RPN2 PE=1 SV=3 | 69.24 | 0 | 0 | 6 | 8 | NF | 2.40E+05 |
| sp \| Q14204 \| DYHC1_HUMAN | DYNC1H1 | Cytoplasmic dynein 1 heavy chain 1 OS=Homo sapiens GN=DYNC1H1 PE=1 SV=5 | 532.07 | 0 | 0 | 7 | 8 | NF | 7.50E+04 |
| sp \| Q86VP6 \| CAND1_HUMAN | CAND1 | Cullin-associated NEDD8-dissociated protein 1 OS=Homo sapiens GN=CAND1 PE=1 SV=2 | 136.29 | 0 | 0 | 7 | 8 | NF | 2.20E+05 |
| sp \| Q9BTW9 \| TBCD_HUMAN | TBCD | Tubulin-specific chaperone D OS=Homo sapiens GN=TBCD PE=1 SV=2 | 132.52 | 0 | 0 | 7 | 8 | NF | 2.60E+06 |
| sp \| O43175 \| SERA_HUMAN | PHGDH | D-3-phosphoglycerate dehydrogenase OS=Homo sapiens GN=PHGDH PE=1 SV=4 | 56.61 | 0 | 0 | 5 | 8 | NF | 4.00E+05 |
| sp \| P43490 \| NAMPT_HUMAN | NAMPT | Nicotinamide phosphoribosyltransferase OS=Homo sapiens GN=NAMPT PE=1 SV=1 | 55.49 | 0 | 0 | 5 | 8 | NF | 1.10E+05 |
| sp \| P39656 \| OST48_HUMAN | DDOST | Dolichyl-diphosphooligosaccharide--protein glycosyltransferase 48 kDa subunit OS=Homo sapiens GN=DDOST PE=1 SV=4 | 50.77 | 0 | 0 | 5 | 8 | NF | 1.90E+05 |
| sp \| Q15758 \| AAAT_HUMAN | SLC1A5 | Neutral amino acid transporter B(0) OS=Homo sapiens GN=SLC1A5 PE=1 SV=2 | 56.56 | 0 | 0 | 4 | 8 | NF | 1.20E+06 |
| sp \| Q92542 \| NICA_HUMAN | NCSTN | Nicastrin OS=Homo sapiens GN=NCSTN PE=1 SV=2 | 78.36 | 0 | 0 | 4 | 8 | NF | 7.00E+05 |
| tr \| A0A024RBS1 \| A0A024RBS1_HUMAN | GCN1L1 | GCN1 general control of amino-acid synthesis 1-like 1 (Yeast), isoform CRA_b OS=Homo sapiens GN=GCN1L1 PE=4 SV=1 | 266.71 | 0 | 0 | 1 | 8 | NF | 3.70E+06 |
| sp \| P02786 \| TFR1_HUMAN | TFRC | Transferrin receptor protein 1 OS=Homo sapiens GN=TFRC PE=1 SV=2 | 84.82 | 0 | 0 | 6 | 7 | NF | 2.60E+05 |
| sp \| P31327 \| CPSM_HUMAN | CPS1 | Carbamoyl-phosphate synthase [ammonia], mitochondrial OS=Homo sapiens GN=CPS1 PE=1 SV=2 | 164.83 | 0 | 0 | 7 | 7 | NF | 1.30E+05 |
| sp \| Q68DK2 \| ZFY26_HUMAN | ZFYVE26 | Zinc finger FYVE domain-containing protein 26 OS=Homo sapiens GN=ZFYVE26 PE=1 SV=3 | 284.4 | 0 | 0 | 7 | 7 | NF | 1.30E+05 |
| sp \| Q86Y56 \| DAAF5_HUMAN | DNAAF5 | Dynein assembly factor 5, axonemal OS=Homo sapiens GN=DNAAF5 PE=1 SV=4 | 93.46 | 0 | 0 | 5 | 7 | NF | 2.10E+05 |
| sp \| Q92562 \| FIG4_HUMAN | FIG4 | Polyphosphoinositide phosphatase OS=Homo sapiens GN=FIG4 PE=1 SV=1 | 103.57 | 0 | 0 | 5 | 7 | NF | 2.30E+05 |
| sp \| Q08AM6 \| VAC14_HUMAN | VAC14 | Protein VAC14 homolog OS=Homo sapiens GN=VAC14 PE=1 SV=1 | 87.92 | 0 | 0 | 5 | 7 | NF | 5.80E+05 |
| sp \| Q86U38 \| NOP9_HUMAN | NOP9 | Nucleolar protein 9 OS=Homo sapiens GN=NOP9 PE=1 SV=1 | 69.39 | 0 | 0 | 4 | 7 | NF | 2.10E+05 |
| sp \| P52701 \| MSH6_HUMAN | MSH6 | DNA mismatch repair protein Msh6 OS=Homo sapiens GN=MSH6 PE=1 SV=2 | 152.69 | 0 | 0 | 5 | 6 | NF | 1.30E+05 |
| sp \| Q96IU4 \| ABHEB_HUMAN | ABHD14B | Protein ABHD14B OS=Homo sapiens GN=ABHD14B PE=1 SV=1 | 22.33 | 0 | 0 | 5 | 6 | NF | 3.40E+05 |
| sp \| Q15645 \| PCH2_HUMAN | TRIP13 | Pachytene checkpoint protein 2 homolog OS=Homo sapiens GN=TRIP13 PE=1 SV=2 | 48.52 | 0 | 0 | 5 | 6 | NF | 2.10E+05 |
| sp \| Q9NVI1 \| FANCI_HUMAN | FANCI | Fanconi anemia group I protein OS=Homo sapiens GN=FANCI PE=1 SV=4 | 149.23 | 0 | 0 | 6 | 6 | NF | 8.90E+04 |
| sp \| Q63HN8 \| RN213_HUMAN | RNF213 | E3 ubiquitin-protein ligase RNF213 OS=Homo sapiens GN=RNF213 PE=1 SV=3 | 591.03 | 0 | 0 | 6 | 6 | NF | 6.80E+04 |
| sp \| P05556 \| ITB1_HUMAN | ITGB1 | Integrin beta-1 OS=Homo sapiens GN=ITGB1 PE=1 SV=2 | 88.36 | 0 | 0 | 5 | 6 | NF | 2.70E+05 |
| sp \| Q14746 \| COG2_HUMAN | COG2 | Conserved oligomeric Golgi complex subunit 2 OS=Homo sapiens GN=COG2 PE=1 SV=1 | 83.16 | 0 | 0 | 5 | 6 | NF | 8.60E+04 |
| sp \| O95373 \| IPO7_HUMAN | IPO7 | Importin-7 OS=Homo sapiens GN=IPO7 PE=1 SV=1 | 119.44 | 0 | 0 | 5 | 6 | NF | 1.00E+05 |
| sp \| O60762 \| DPM1_HUMAN | DPM1 | Dolichol-phosphate mannosyltransferase subunit 1 OS=Homo sapiens GN=DPM1 PE=1 SV=1 | 29.62 | 0 | 0 | 4 | 6 | NF | 2.90E+05 |
| sp \| P12236 \| ADT3_HUMAN | SLC25A6 | ADP/ATP translocase 3 OS=Homo sapiens GN=SLC25A6 PE=1 SV=4 | 32.85 | 0 | 0 | 4 | 6 | NF | 1.30E+06 |
| sp \| Q96G23 \| CERS2_HUMAN | CERS2 | Ceramide synthase 2 OS=Homo sapiens GN=CERS2 PE=1 SV=1 | 44.85 | 0 | 0 | 3 | 6 | NF | 9.50E+04 |
| sp \| O00410 \| IPO5_HUMAN | IPO5 | Importin-5 OS=Homo sapiens GN=IPO5 PE=1 SV=4 | 123.55 | 0 | 0 | 3 | 6 | NF | 1.80E+06 |
| sp \| P55072 \| TERA_HUMAN | VCP | Transitional endoplasmic reticulum ATPase OS=Homo sapiens GN=VCP PE=1 SV=4 | 89.27 | 0 | 0 | 4 | 5 | NF | 1.40E+05 |
| sp \| P50454 \| SERPH_HUMAN | SERPINH1 | Serpin H1 OS=Homo sapiens GN=SERPINH1 PE=1 SV=2 | 46.41 | 0 | 0 | 5 | 5 | NF | 1.20E+05 |
| sp \| Q5T160 \| SYRM_HUMAN | RARS2 | Probable arginine--tRNA ligase, mitochondrial OS=Homo sapiens GN=RARS2 PE=1 SV=1 | 65.46 | 0 | 0 | 4 | 5 | NF | 1.20E+05 |
| sp \| P17858 \| PFKAL_HUMAN | PFKL | ATP-dependent 6-phosphofructokinase, liver type OS=Homo sapiens GN=PFKL PE=1 SV=6 | 84.96 | 0 | 0 | 4 | 5 | NF | 1.20E+05 |
| sp \| O60313 \| OPA1_HUMAN | OPA1 | Dynamin-like 120 kDa protein, mitochondrial OS=Homo sapiens GN=OPA1 PE=1 SV=3 | 111.56 | 0 | 0 | 4 | 5 | NF | 2.80E+06 |
| sp \| O75531 \| BAF_HUMAN | BANF1 | Barrier-to-autointegration factor OS=Homo sapiens GN=BANF1 PE=1 SV=1 | 10.05 | 0 | 0 | 4 | 5 | NF | 2.10E+05 |
| sp \| Q6NXR4 \| TTI2_HUMAN | TTI2 | TELO2-interacting protein 2 OS=Homo sapiens GN=TTI2 PE=1 SV=1 | 56.88 | 0 | 0 | 4 | 5 | NF | 2.10E+05 |
| sp \| Q9NYY8 \| FAKD2_HUMAN | FASTKD2 | FAST kinase domain-containing protein 2, mitochondrial OS=Homo sapiens GN=FASTKD2 PE=1 SV=1 | 81.41 | 0 | 0 | 4 | 5 | NF | 1.00E+05 |
| sp \| Q9H583 \| HEAT1_HUMAN | HEATR1 | HEAT repeat-containing protein 1 OS=Homo sapiens GN=HEATR1 PE=1 SV=3 | 242.22 | 0 | 0 | 4 | 5 | NF | 6.60E+04 |
| sp \| Q9BTV4 \| TMM43_HUMAN | TMEM43 | Transmembrane protein 43 OS=Homo sapiens GN=TMEM43 PE=1 SV=1 | 44.85 | 0 | 0 | 3 | 5 | NF | 9.50E+04 |
| sp \| Q9Y679 \| AUP1_HUMAN | AUP1 | Ancient ubiquitous protein 1 OS=Homo sapiens GN=AUP1 PE=1 SV=1 | 52.99 | 0 | 0 | 3 | 5 | NF | 5.60E+04 |
| sp \| O00161 \| SNP23_HUMAN | SNAP23 | Synaptosomal-associated protein 23 OS=Homo sapiens GN=SNAP23 PE=1 SV=1 | 23.34 | 0 | 0 | 3 | 5 | NF | 8.80E+04 |
| sp \| P55209 \| NP1L1_HUMAN | NAP1L1 | Nucleosome assembly protein 1-like 1 OS=Homo sapiens GN=NAP1L1 PE=1 SV=1 | 45.35 | 0 | 0 | 3 | 5 | NF | 1.30E+05 |
| sp \| Q8IWA5 \| CTL2_HUMAN | SLC44A2 | Choline transporter-like protein 2 OS=Homo sapiens GN=SLC44A2 PE=1 SV=3 | 80.07 | 0 | 0 | 3 | 5 | NF | 1.40E+05 |
| sp \| P04350 \| TBB4A_HUMAN | TUBB4A | Tubulin beta-4A chain OS=Homo sapiens GN=TUBB4A PE=1 SV=2 | 49.55 | 0 | 0 | 2 | 5 | NF | 1.80E+06 |
| sp \| P51571 \| SSRD_HUMAN | SSR4 | Translocon-associated protein subunit delta OS=Homo sapiens GN=SSR4 PE=1 SV=1 | 18.99 | 0 | 0 | 2 | 5 | NF | 3.60E+05 |
| sp \| P30508 \| 1C12_HUMAN | HLA-C | HLA class I histocompatibility antigen, Cw-12 alpha chain OS=Homo sapiens GN=HLA-C PE=1 SV=2 | 40.86 | 0 | 0 | 1 | 5 | NF | 9.70E+05 |
| sp \| P53621 \| COPA_HUMAN | COPA | Coatomer subunit alpha OS=Homo sapiens GN=COPA PE=1 SV=2 | 138.26 | 0 | 0 | 4 | 4 | NF | 4.80E+05 |
| sp \| Q92973 \| TNPO1_HUMAN | TNPO1 | Transportin-1 OS=Homo sapiens GN=TNPO1 PE=1 SV=2 | 102.29 | 0 | 0 | 3 | 4 | NF | 1.20E+05 |
| sp \| P62081 \| RS7_HUMAN | RPS7 | 40S ribosomal protein S7 OS=Homo sapiens GN=RPS7 PE=1 SV=1 | 22.11 | 0 | 0 | 3 | 4 | NF | 2.10E+05 |
| sp \| Q14974 \| IMB1_HUMAN | KPNB1 | Importin subunit beta-1 OS=Homo sapiens GN=KPNB1 PE=1 SV=2 | 97.11 | 0 | 0 | 3 | 4 | NF | 1.30E+05 |
| sp \| Q9UID3 \| VPS51_HUMAN | VPS51 | Vacuolar protein sorting-associated protein 51 homolog OS=Homo sapiens GN=VPS51 PE=1 SV=2 | 85.99 | 0 | 0 | 4 | 4 | NF | 4.00E+04 |
| sp \| A0FGR8 \| ESYT2_HUMAN | ESYT2 | Extended synaptotagmin-2 OS=Homo sapiens GN=ESYT2 PE=1 SV=1 | 102.29 | 0 | 0 | 4 | 4 | NF | 5.50E+04 |
| sp \| Q6DD88 \| ATLA3_HUMAN | ATL3 | Atlastin-3 OS=Homo sapiens GN=ATL3 PE=1 SV=1 | 60.5 | 0 | 0 | 4 | 4 | NF | 2.00E+05 |
| sp \| P26599 \| PTBP1_HUMAN | PTBP1 | Polypyrimidine tract-binding protein 1 OS=Homo sapiens GN=PTBP1 PE=1 SV=1 | 57.19 | 0 | 0 | 4 | 4 | NF | 7.90E+04 |
| sp \| P06493 \| CDK1_HUMAN | CDK1 | Cyclin-dependent kinase 1 OS=Homo sapiens GN=CDK1 PE=1 SV=3 | 34.07 | 0 | 0 | 3 | 4 | NF | 9.50E+04 |
| sp \| Q03518 \| TAP1_HUMAN | TAP1 | Antigen peptide transporter 1 OS=Homo sapiens GN=TAP1 PE=1 SV=2 | 87.16 | 0 | 0 | 2 | 4 | NF | 4.10E+04 |

(NF: not found)

| Reference | Gene Symbol | Annotation | MWT(kDa) | Unique FA | Total FA | Unique JX4-FA | Total JX4-FA | Sum Intensity FA | Sum Intensity JX4-FA |
|---|---|---|---|---|---|---|---|---|---|
| sp\|Q12797\|ASPH_HUMAN | ASPH | Aspartyl/asparaginyl beta-hydroxylase OS=Homo sapiens GN=ASPH PE=1 SV=3 | 85.81 | 0 | 0 | 3 | 4 | NF | 1.60E+05 |
| sp\|O75947\|ATP5H_HUMAN | ATP5H | ATP synthase subunit d, mitochondrial OS=Homo sapiens GN=ATP5H PE=1 SV=3 | 18.48 | 0 | 0 | 3 | 4 | NF | 1.30E+05 |
| sp\|Q9H3U1\|UN45A_HUMAN | UNC45A | Protein unc-45 homolog A OS=Homo sapiens GN=UNC45A PE=1 SV=1 | 103.01 | 0 | 0 | 3 | 4 | NF | 1.10E+05 |
| sp\|P62847\|RS24_HUMAN | RPS24 | 40S ribosomal protein S24 OS=Homo sapiens GN=RPS24 PE=1 SV=1 | 15.41 | 0 | 0 | 3 | 4 | NF | 9.60E+04 |
| sp\|P57678\|GEMI4_HUMAN | GEMIN4 | Gem-associated protein 4 OS=Homo sapiens GN=GEMIN4 PE=1 SV=2 | 119.96 | 0 | 0 | 3 | 4 | NF | 2.70E+04 |
| sp\|Q9NZ08\|ERAP1_HUMAN | ERAP1 | Endoplasmic reticulum aminopeptidase 1 OS=Homo sapiens GN=ERAP1 PE=1 SV=3 | 107.17 | 0 | 0 | 3 | 4 | NF | 8.70E+04 |
| sp\|Q99714\|HCD2_HUMAN | HSD17B10 | 3-hydroxyacyl-CoA dehydrogenase type-2 OS=Homo sapiens GN=HSD17B10 PE=1 SV=3 | 26.91 | 0 | 0 | 3 | 4 | NF | 5.90E+04 |
| sp\|Q9BRX8\|F213A_HUMAN | FAM213A | Redox-regulatory protein FAM213A OS=Homo sapiens GN=FAM213A PE=1 SV=3 | 25.75 | 0 | 0 | 3 | 4 | NF | 2.10E+05 |
| sp\|O60427\|FADS1_HUMAN | FADS1 | Fatty acid desaturase 1 OS=Homo sapiens GN=FADS1 PE=1 SV=3 | 51.93 | 0 | 0 | 3 | 4 | NF | 5.00E+04 |
| sp\|Q9Y6N5\|SQOR_HUMAN | SQOR | Sulfide:quinone oxidoreductase, mitochondrial OS=Homo sapiens GN=SQOR PE=1 SV=1 | 49.93 | 0 | 0 | 3 | 4 | NF | 4.70E+04 |
| sp\|P16615\|AT2A2_HUMAN | ATP2A2 | Sarcoplasmic/endoplasmic reticulum calcium ATPase 2 OS=Homo sapiens GN=ATP2A2 PE=1 SV=1 | 114.68 | 0 | 0 | 3 | 4 | NF | 7.70E+04 |
| sp\|P68366\|TBA4A_HUMAN | TUBA4A | Tubulin alpha-4A chain OS=Homo sapiens GN=TUBA4A PE=1 SV=1 | 49.89 | 0 | 0 | 2 | 4 | NF | 7.10E+05 |
| sp\|A6NHL2\|TBAL3_HUMAN | TUBAL3 | Tubulin alpha chain-like 3 OS=Homo sapiens GN=TUBAL3 PE=1 SV=2 | 49.88 | 0 | 0 | 2 | 4 | NF | 1.10E+05 |
| sp\|Q8NCG7\|DGLB_HUMAN | DAGLB | Sn1-specific diacylglycerol lipase beta OS=Homo sapiens GN=DAGLB PE=1 SV=2 | 73.68 | 0 | 0 | 2 | 4 | NF | 7.90E+04 |
| sp\|Q9Y230\|RUVB2_HUMAN | RUVBL2 | RuvB-like 2 OS=Homo sapiens GN=RUVBL2 PE=1 SV=3 | 51.12 | 0 | 0 | 2 | 4 | NF | 1.40E+05 |
| sp\|P43003\|EAA1_HUMAN | SLC1A3 | Excitatory amino acid transporter 1 OS=Homo sapiens GN=SLC1A3 PE=1 SV=1 | 59.53 | 0 | 0 | 2 | 4 | NF | 3.30E+05 |
| sp\|Q9Y6M0\|TEST_HUMAN | PRSS21 | Testisin OS=Homo sapiens GN=PRSS21 PE=1 SV=1 | 34.86 | 0 | 0 | 2 | 4 | NF | 9.40E+04 |
| sp\|Q8N6M3\|FITM2_HUMAN | FITM2 | Fat storage-inducing transmembrane protein 2 OS=Homo sapiens GN=FITM2 PE=2 SV=1 | 29.84 | 0 | 0 | 2 | 4 | NF | 4.20E+05 |
| sp\|P39023\|RL3_HUMAN | RPL3 | 60S ribosomal protein L3 OS=Homo sapiens GN=RPL3 PE=1 SV=2 | 46.08 | 0 | 0 | 2 | 4 | NF | 1.40E+05 |
| sp\|Q9Y2S7\|PDIP2_HUMAN | POLDIP2 | Polymerase delta-interacting protein 2 OS=Homo sapiens GN=POLDIP2 PE=1 SV=1 | 42.01 | 0 | 0 | 3 | 3 | NF | 5.10E+04 |
| sp\|P40939\|ECHA_HUMAN | HADHA | Trifunctional enzyme subunit alpha, mitochondrial OS=Homo sapiens GN=HADHA PE=1 SV=2 | 82.95 | 0 | 0 | 3 | 3 | NF | 4.00E+04 |
| sp\|Q9Y265\|RUVB1_HUMAN | RUVBL1 | RuvB-like 1 OS=Homo sapiens GN=RUVBL1 PE=1 SV=1 | 50.2 | 0 | 0 | 2 | 3 | NF | 8.60E+04 |
| sp\|O43570\|CAH12_HUMAN | CA12 | Carbonic anhydrase 12 OS=Homo sapiens GN=CA12 PE=1 SV=1 | 39.43 | 0 | 0 | 2 | 3 | NF | 1.40E+05 |
| sp\|Q6IAA8\|LTOR1_HUMAN | LAMTOR1 | Ragulator complex protein LAMTOR1 OS=Homo sapiens GN=LAMTOR1 PE=1 SV=2 | 17.73 | 0 | 0 | 2 | 3 | NF | 2.00E+04 |
| sp\|O00560\|SDCB1_HUMAN | SDCBP | Syntenin-1 OS=Homo sapiens GN=SDCBP PE=1 SV=1 | 32.42 | 0 | 0 | 1 | 3 | NF | 2.30E+04 |
| sp\|P27105\|STOM_HUMAN | STOM | Erythrocyte band 7 integral membrane protein OS=Homo sapiens GN=STOM PE=1 SV=3 | 31.71 | 0 | 0 | 3 | 3 | NF | 4.00E+04 |
| sp\|Q14697\|GANAB_HUMAN | GANAB | Neutral alpha-glucosidase AB OS=Homo sapiens GN=GANAB PE=1 SV=3 | 106.81 | 0 | 0 | 3 | 3 | NF | 3.50E+04 |
| sp\|P54652\|HSP72_HUMAN | HSPA2 | Heat shock-related 70 kDa protein 2 OS=Homo sapiens GN=HSPA2 PE=1 SV=1 | 69.98 | 0 | 0 | 2 | 3 | NF | 3.40E+05 |
| sp\|Q14257\|RCN2_HUMAN | RCN2 | Reticulocalbin-2 OS=Homo sapiens GN=RCN2 PE=1 SV=1 | 36.85 | 0 | 0 | 2 | 3 | NF | 8.10E+04 |
| sp\|Q9Y6E2\|BZW2_HUMAN | BZW2 | Basic leucine zipper and W2 domain-containing protein 2 OS=Homo sapiens GN=BZW2 PE=1 SV=1 | 48.13 | 0 | 0 | 3 | 3 | NF | 6.90E+04 |
| sp\|O15397\|IPO8_HUMAN | IPO8 | Importin-8 OS=Homo sapiens GN=IPO8 PE=1 SV=2 | 119.86 | 0 | 0 | 2 | 3 | NF | 6.20E+04 |
| sp\|P62805\|H4_HUMAN | HIST1H4A | Histone H4 OS=Homo sapiens GN=HIST1H4A PE=1 SV=2 | 11.36 | 0 | 0 | 2 | 3 | NF | 6.30E+04 |
| sp\|Q99733\|NP1L4_HUMAN | NAP1L4 | Nucleosome assembly protein 1-like 4 OS=Homo sapiens GN=NAP1L4 PE=1 SV=1 | 42.8 | 0 | 0 | 3 | 3 | NF | 4.80E+04 |
| sp\|P51648\|AL3A2_HUMAN | ALDH3A2 | Fatty aldehyde dehydrogenase OS=Homo sapiens GN=ALDH3A2 PE=1 SV=1 | 54.81 | 0 | 0 | 3 | 3 | NF | 7.80E+04 |
| sp\|P56134\|ATPK_HUMAN | ATP5J2 | ATP synthase subunit f, mitochondrial OS=Homo sapiens GN=ATP5J2 PE=1 SV=3 | 10.91 | 0 | 0 | 3 | 3 | NF | 7.40E+04 |
| sp\|Q5VYK3\|ECM29_HUMAN | ECM29 | Proteasome-associated protein ECM29 homolog OS=Homo sapiens GN=ECM29 PE=1 SV=2 | 204.16 | 0 | 0 | 3 | 3 | NF | 5.40E+04 |
| sp\|P55786\|PSA_HUMAN | NPEPPS | Puromycin-sensitive aminopeptidase OS=Homo sapiens GN=NPEPPS PE=1 SV=2 | 103.21 | 0 | 0 | 3 | 3 | NF | 4.60E+04 |
| sp\|P22102\|PUR2_HUMAN | GART | Trifunctional purine biosynthetic protein adenosine-3 OS=Homo sapiens GN=GART PE=1 SV=1 | 107.7 | 0 | 0 | 3 | 3 | NF | 3.30E+04 |
| sp\|P22061\|PIMT_HUMAN | PCMT1 | Protein-L-isoaspartate(D-aspartate) O-methyltransferase OS=Homo sapiens GN=PCMT1 PE=1 SV=4 | 24.62 | 0 | 0 | 3 | 3 | NF | 3.30E+04 |
| sp\|A1L0T0\|ILVBL_HUMAN | ILVBL | Acetolactate synthase-like protein OS=Homo sapiens GN=ILVBL PE=1 SV=2 | 67.82 | 0 | 0 | 3 | 3 | NF | 3.20E+04 |
| sp\|Q15386\|UBE3C_HUMAN | UBE3C | Ubiquitin-protein ligase E3C OS=Homo sapiens GN=UBE3C PE=1 SV=3 | 123.84 | 0 | 0 | 3 | 3 | NF | 6.10E+04 |
| sp\|O75489\|NDUS3_HUMAN | NDUFS3 | NADH dehydrogenase [ubiquinone] iron-sulfur protein 3, mitochondrial OS=Homo sapiens GN=NDUFS3 PE=1 SV=1 | 30.22 | 0 | 0 | 3 | 3 | NF | 3.50E+04 |
| sp\|Q8WTW3\|COG1_HUMAN | COG1 | Conserved oligomeric Golgi complex subunit 1 OS=Homo sapiens GN=COG1 PE=1 SV=1 | 108.91 | 0 | 0 | 3 | 3 | NF | 3.90E+04 |
| sp\|P52597\|HNRPF_HUMAN | HNRNPF | Heterogeneous nuclear ribonucleoprotein F OS=Homo sapiens GN=HNRNPF PE=1 SV=3 | 45.64 | 0 | 0 | 2 | 3 | NF | 1.00E+05 |
| sp\|Q3ZCQ8\|TIM50_HUMAN | TIMM50 | Mitochondrial import inner membrane translocase subunit TIM50 OS=Homo sapiens GN=TIMM50 PE=1 SV=2 | 39.62 | 0 | 0 | 2 | 3 | NF | 7.50E+04 |
| sp\|Q8NF37\|PCAT1_HUMAN | LPCAT1 | Lysophosphatidylcholine acyltransferase 1 OS=Homo sapiens GN=LPCAT1 PE=1 SV=2 | 59.11 | 0 | 0 | 2 | 3 | NF | 1.10E+05 |
| sp\|P05141\|ADT2_HUMAN | SLC25A5 | ADP/ATP translocase 2 OS=Homo sapiens GN=SLC25A5 PE=1 SV=7 | 32.83 | 0 | 0 | 2 | 3 | NF | 2.70E+05 |
| sp\|Q9BUN8\|DERL1_HUMAN | DERL1 | Derlin-1 OS=Homo sapiens GN=DERL1 PE=1 SV=1 | 28.78 | 0 | 0 | 2 | 3 | NF | 5.90E+04 |
| sp\|P33993\|MCM7_HUMAN | MCM7 | DNA replication licensing factor MCM7 OS=Homo sapiens GN=MCM7 PE=1 SV=4 | 81.26 | 0 | 0 | 2 | 3 | NF | 5.10E+04 |
| sp\|P62906\|RL10A_HUMAN | RPL10A | 60S ribosomal protein L10a OS=Homo sapiens GN=RPL10A PE=1 SV=2 | 24.82 | 0 | 0 | 2 | 3 | NF | 7.70E+04 |
| sp\|P48444\|COPD_HUMAN | ARCN1 | Coatomer subunit delta OS=Homo sapiens GN=ARCN1 PE=1 SV=1 | 57.17 | 0 | 0 | 2 | 3 | NF | 4.60E+04 |
| sp\|P62269\|RS18_HUMAN | RPS18 | 40S ribosomal protein S18 OS=Homo sapiens GN=RPS18 PE=1 SV=3 | 17.71 | 0 | 0 | 2 | 3 | NF | 9.50E+04 |
| sp\|Q5PRF9\|SMAG2_HUMAN | SAMD4B | Protein Smaug homolog 2 OS=Homo sapiens GN=SAMD4B PE=1 SV=1 | 75.44 | 0 | 0 | 2 | 3 | NF | 5.80E+04 |
| sp\|P61353\|RL27_HUMAN | RPL27 | 60S ribosomal protein L27 OS=Homo sapiens GN=RPL27 PE=1 SV=2 | 15.79 | 0 | 0 | 2 | 3 | NF | 6.50E+04 |
| tr\|Q8IWP6\|Q8IWP6_HUMAN | | Tubulin beta chain OS=Homo sapiens PE=2 SV=1 | 49.72 | 0 | 0 | 1 | 3 | NF | 3.60E+05 |
| sp\|Q9Y5Y2\|NUBP2_HUMAN | NUBP2 | Cytosolic Fe-S cluster assembly factor NUBP2 OS=Homo sapiens GN=NUBP2 PE=1 SV=1 | 28.81 | 0 | 0 | 1 | 3 | NF | 5.60E+04 |
| sp\|P57088\|TMM33_HUMAN | TMEM33 | Transmembrane protein 33 OS=Homo sapiens GN=TMEM33 PE=1 SV=2 | 27.96 | 0 | 0 | 1 | 3 | NF | 4.20E+05 |
| sp\|Q8WY22\|BRI3B_HUMAN | BRI3BP | BRI3-binding protein OS=Homo sapiens GN=BRI3BP PE=1 SV=1 | 27.82 | 0 | 0 | 1 | 3 | NF | 8.00E+04 |
| sp\|Q9H2V7\|SPNS1_HUMAN | SPNS1 | Protein spinster homolog 1 OS=Homo sapiens GN=SPNS1 PE=1 SV=1 | 56.59 | 0 | 0 | 1 | 3 | NF | 5.50E+04 |
| sp\|Q00610\|CLH1_HUMAN | CLTC | Clathrin heavy chain 1 OS=Homo sapiens GN=CLTC PE=1 SV=5 | 191.49 | 0 | 0 | 2 | 2 | NF | 1.30E+04 |
| sp\|O75915\|PRAF3_HUMAN | ARL6IP5 | PRA1 family protein 3 OS=Homo sapiens GN=ARL6IP5 PE=1 SV=1 | 21.6 | 0 | 0 | 1 | 2 | NF | 4.40E+04 |
| sp\|P56192\|SYMC_HUMAN | MARS | Methionine--tRNA ligase, cytoplasmic OS=Homo sapiens GN=MARS PE=1 SV=2 | 101.05 | 0 | 0 | 2 | 2 | NF | 3.20E+04 |
| sp\|O75340\|PDCD6_HUMAN | PDCD6 | Programmed cell death protein 6 OS=Homo sapiens GN=PDCD6 PE=1 SV=1 | 21.85 | 0 | 0 | 1 | 2 | NF | 8.30E+04 |
| sp\|P20020\|AT2B1_HUMAN | ATP2B1 | Plasma membrane calcium-transporting ATPase 1 OS=Homo sapiens GN=ATP2B1 PE=1 SV=3 | 138.67 | 0 | 0 | 1 | 2 | NF | 4.90E+04 |
| sp\|Q01650\|LAT1_HUMAN | SLC7A5 | Large neutral amino acids transporter small subunit 1 OS=Homo sapiens GN=SLC7A5 PE=1 SV=2 | 54.97 | 0 | 0 | 2 | 2 | NF | 4.30E+04 |
| sp\|P23634\|AT2B4_HUMAN | ATP2B4 | Plasma membrane calcium-transporting ATPase 4 OS=Homo sapiens GN=ATP2B4 PE=1 SV=2 | 137.83 | 0 | 0 | 2 | 2 | NF | 7.40E+04 |
| sp\|P15880\|RS2_HUMAN | RPS2 | 40S ribosomal protein S2 OS=Homo sapiens GN=RPS2 PE=1 SV=2 | 31.3 | 0 | 0 | 2 | 2 | NF | 7.10E+04 |
| sp\|Q9Y5L0\|TNPO3_HUMAN | TNPO3 | Transportin-3 OS=Homo sapiens GN=TNPO3 PE=1 SV=3 | 104.14 | 0 | 0 | 2 | 2 | NF | 2.10E+04 |
| sp\|P30101\|PDIA3_HUMAN | PDIA3 | Protein disulfide-isomerase A3 OS=Homo sapiens GN=PDIA3 PE=1 SV=4 | 56.75 | 0 | 0 | 2 | 2 | NF | 2.60E+04 |
| sp\|Q96D53\|COQ8B_HUMAN | COQ8B | Atypical kinase COQ8B, mitochondrial OS=Homo sapiens GN=COQ8B PE=1 SV=2 | 60.03 | 0 | 0 | 2 | 2 | NF | 2.10E+04 |
| sp\|P31689\|DNJA1_HUMAN | DNAJA1 | DnaJ homolog subfamily A member 1 OS=Homo sapiens GN=DNAJA1 PE=1 SV=2 | 44.84 | 0 | 0 | 2 | 2 | NF | 9.00E+04 |
| sp\|Q29836\|1B67_HUMAN | HLA-B | HLA class I histocompatibility antigen, B-67 alpha chain OS=Homo sapiens GN=HLA-B PE=1 SV=1 | 40.32 | 0 | 0 | 2 | 2 | NF | 2.20E+05 |
| sp\|P61204\|ARF3_HUMAN | ARF3 | ADP-ribosylation factor 3 OS=Homo sapiens GN=ARF3 PE=1 SV=2 | 20.59 | 0 | 0 | 2 | 2 | NF | 4.40E+04 |
| sp\|Q96QD8\|S38A2_HUMAN | SLC38A2 | Sodium-coupled neutral amino acid transporter 2 OS=Homo sapiens GN=SLC38A2 PE=1 SV=2 | 55.99 | 0 | 0 | 2 | 2 | NF | 3.50E+04 |
| sp\|Q6NXE6\|ARMC6_HUMAN | ARMC6 | Armadillo repeat-containing protein 6 OS=Homo sapiens GN=ARMC6 PE=1 SV=2 | 54.11 | 0 | 0 | 2 | 2 | NF | 3.90E+04 |
| sp\|O60884\|DNJA2_HUMAN | DNAJA2 | DnaJ homolog subfamily A member 2 OS=Homo sapiens GN=DNAJA2 PE=1 SV=1 | 45.72 | 0 | 0 | 2 | 2 | NF | 4.50E+04 |
| sp\|P17980\|PRS6A_HUMAN | PSMC3 | 26S protease regulatory subunit 6A OS=Homo sapiens GN=PSMC3 PE=1 SV=3 | 49.17 | 0 | 0 | 2 | 2 | NF | 2.00E+04 |
| sp\|P52789\|HXK2_HUMAN | HK2 | Hexokinase-2 OS=Homo sapiens GN=HK2 PE=1 SV=2 | 102.31 | 0 | 0 | 2 | 2 | NF | 3.50E+04 |
| sp\|Q03169\|TNAP2_HUMAN | TNFAIP2 | Tumor necrosis factor alpha-induced protein 2 OS=Homo sapiens GN=TNFAIP2 PE=2 SV=2 | 72.62 | 0 | 0 | 2 | 2 | NF | 1.60E+04 |
| sp\|O75787\|RENR_HUMAN | ATP6AP2 | Renin receptor OS=Homo sapiens GN=ATP6AP2 PE=1 SV=2 | 38.98 | 0 | 0 | 2 | 2 | NF | 3.40E+04 |
| sp\|Q96BM9\|ARL8A_HUMAN | ARL8A | ADP-ribosylation factor-like protein 8A OS=Homo sapiens GN=ARL8A PE=1 SV=1 | 21.4 | 0 | 0 | 2 | 2 | NF | 2.50E+04 |
| sp\|P08238\|HS90B_HUMAN | HSP90AB1 | Heat shock protein HSP 90-beta OS=Homo sapiens GN=HSP90AB1 PE=1 SV=4 | 83.21 | 0 | 0 | 2 | 2 | NF | 5.00E+04 |
| sp\|O14983\|AT2A1_HUMAN | ATP2A1 | Sarcoplasmic/endoplasmic reticulum calcium ATPase 1 OS=Homo sapiens GN=ATP2A1 PE=1 SV=1 | 110.18 | 0 | 0 | 2 | 2 | NF | 6.70E+04 |
| sp\|Q58FF8\|H90B2_HUMAN | HSP90AB2P | Putative heat shock protein HSP 90-beta 2 OS=Homo sapiens GN=HSP90AB2P PE=1 SV=2 | 44.32 | 0 | 0 | 2 | 2 | NF | 1.20E+05 |
| sp\|Q9H1C4\|UN93B_HUMAN | UNC93B1 | Protein unc-93 homolog B1 OS=Homo sapiens GN=UNC93B1 PE=1 SV=2 | 66.59 | 0 | 0 | 2 | 2 | NF | 2.90E+04 |
| sp\|P48047\|ATPO_HUMAN | ATP5O | ATP synthase subunit O, mitochondrial OS=Homo sapiens GN=ATP5O PE=1 SV=1 | 23.26 | 0 | 0 | 2 | 2 | NF | 2.30E+04 |
| sp\|Q8WVX9\|FACR1_HUMAN | FAR1 | Fatty acyl-CoA reductase 1 OS=Homo sapiens GN=FAR1 PE=1 SV=1 | 59.32 | 0 | 0 | 2 | 2 | NF | 5.10E+04 |
| sp\|Q9Y4R8\|TELO2_HUMAN | TELO2 | Telomere length regulation protein TEL2 homolog OS=Homo sapiens GN=TELO2 PE=1 SV=2 | 91.69 | 0 | 0 | 2 | 2 | NF | 3.60E+04 |
| sp\|P40227\|TCPZ_HUMAN | CCT6A | T-complex protein 1 subunit zeta OS=Homo sapiens GN=CCT6A PE=1 SV=3 | 57.99 | 0 | 0 | 2 | 2 | NF | 1.40E+04 |
| sp\|Q13724\|MOGS_HUMAN | MOGS | Mannosyl-oligosaccharide glucosidase OS=Homo sapiens GN=MOGS PE=1 SV=5 | 91.86 | 0 | 0 | 2 | 2 | NF | 2.80E+04 |
| sp\|Q14240\|IF4A2_HUMAN | EIF4A2 | Eukaryotic initiation factor 4A-II OS=Homo sapiens GN=EIF4A2 PE=1 SV=2 | 46.37 | 0 | 0 | 2 | 2 | NF | 2.40E+04 |
| sp\|P12956\|XRCC6_HUMAN | XRCC6 | X-ray repair cross-complementing protein 6 OS=Homo sapiens GN=XRCC6 PE=1 SV=2 | 69.8 | 0 | 0 | 2 | 2 | NF | 2.50E+04 |
| sp\|P40616\|ARL1_HUMAN | ARL1 | ADP-ribosylation factor-like protein 1 OS=Homo sapiens GN=ARL1 PE=1 SV=1 | 20.4 | 0 | 0 | 2 | 2 | NF | 3.20E+04 |
| sp\|P49768\|PSN1_HUMAN | PSEN1 | Presenilin-1 OS=Homo sapiens GN=PSEN1 PE=1 SV=1 | 52.63 | 0 | 0 | 2 | 2 | NF | 1.90E+04 |
| sp\|P21359\|NF1_HUMAN | NF1 | Neurofibromin OS=Homo sapiens GN=NF1 PE=1 SV=2 | 319.17 | 0 | 0 | 2 | 2 | NF | 2.30E+04 |
| sp\|P32969\|RL9_HUMAN | RPL9 | 60S ribosomal protein L9 OS=Homo sapiens GN=RPL9 PE=1 SV=1 | 21.85 | 0 | 0 | 2 | 2 | NF | 2.70E+04 |
| sp\|Q12907\|LMAN2_HUMAN | LMAN2 | Vesicular integral-membrane protein VIP36 OS=Homo sapiens GN=LMAN2 PE=1 SV=1 | 40.2 | 0 | 0 | 2 | 2 | NF | 3.00E+04 |
| sp\|Q92621\|NU205_HUMAN | NUP205 | Nuclear pore complex protein Nup205 OS=Homo sapiens GN=NUP205 PE=1 SV=3 | 227.78 | 0 | 0 | 2 | 2 | NF | 3.50E+04 |
| sp\|Q13283\|G3BP1_HUMAN | G3BP1 | Ras GTPase-activating protein-binding protein 1 OS=Homo sapiens GN=G3BP1 PE=1 SV=1 | 52.13 | 0 | 0 | 2 | 2 | NF | 2.80E+04 |
| sp\|Q9BQ67\|GRWD1_HUMAN | GRWD1 | Glutamate-rich WD repeat-containing protein 1 OS=Homo sapiens GN=GRWD1 PE=1 SV=1 | 49.39 | 0 | 0 | 2 | 2 | NF | 2.10E+04 |
| sp\|O95674\|CDS2_HUMAN | CDS2 | Phosphatidate cytidylyltransferase 2 OS=Homo sapiens GN=CDS2 PE=1 SV=1 | 51.38 | 0 | 0 | 2 | 2 | NF | 1.40E+04 |

(NF: not found)

| Reference | Gene Symbol | Annotation | MWT(kDa) | Unique FA | Total FA | Unique JX4-FA | Total JX4-FA | Sum Intensity FA | Sum Intensity JX4-FA |
|---|---|---|---|---|---|---|---|---|---|
| sp\|Q96JG6\|VPS50_HUMAN | VPS50 | Syndetin OS=Homo sapiens GN=VPS50 PE=1 SV=3 | 111.1 | 0 | 0 | 2 | 2 | NF | 3.50E+04 |
| sp\|Q96S52\|PIGS_HUMAN | PIGS | GPI transamidase component PIG-S OS=Homo sapiens GN=PIGS PE=1 SV=3 | 61.62 | 0 | 0 | 2 | 2 | NF | 4.70E+04 |
| sp\|Q96N66\|MBOA7_HUMAN | MBOAT7 | Lysophospholipid acyltransferase 7 OS=Homo sapiens GN=MBOAT7 PE=1 SV=2 | 52.73 | 0 | 0 | 2 | 2 | NF | 2.40E+04 |
| sp\|P11586\|C1TC_HUMAN | MTHFD1 | C-1-tetrahydrofolate synthase, cytoplasmic OS=Homo sapiens GN=MTHFD1 PE=1 SV=3 | 101.5 | 0 | 0 | 2 | 2 | NF | 2.60E+04 |
| sp\|Q08211\|DHX9_HUMAN | DHX9 | ATP-dependent RNA helicase A OS=Homo sapiens GN=DHX9 PE=1 SV=4 | 140.87 | 0 | 0 | 2 | 2 | NF | 2.30E+04 |
| sp\|P26373\|RL13_HUMAN | RPL13 | 60S ribosomal protein L13 OS=Homo sapiens GN=RPL13 PE=1 SV=4 | 24.25 | 0 | 0 | 2 | 2 | NF | 4.80E+04 |
| sp\|Q96JB2\|COG3_HUMAN | COG3 | Conserved oligomeric Golgi complex subunit 3 OS=Homo sapiens GN=COG3 PE=1 SV=3 | 94.04 | 0 | 0 | 2 | 2 | NF | 4.60E+04 |
| sp\|P48651\|PTSS1_HUMAN | PTDSS1 | Phosphatidylserine synthase 1 OS=Homo sapiens GN=PTDSS1 PE=1 SV=1 | 55.49 | 0 | 0 | 2 | 2 | NF | 2.90E+04 |
| sp\|Q9UBB4\|ATX10_HUMAN | ATXN10 | Ataxin-10 OS=Homo sapiens GN=ATXN10 PE=1 SV=1 | 53.45 | 0 | 0 | 2 | 2 | NF | 1.70E+04 |
| sp\|Q6PGP7\|TTC37_HUMAN | TTC37 | Tetratricopeptide repeat protein 37 OS=Homo sapiens GN=TTC37 PE=1 SV=1 | 175.37 | 0 | 0 | 2 | 2 | NF | 1.60E+04 |
| sp\|Q9Y624\|JAM1_HUMAN | F11R | Junctional adhesion molecule A OS=Homo sapiens GN=F11R PE=1 SV=1 | 32.56 | 0 | 0 | 2 | 2 | NF | 4.80E+04 |
| sp\|Q96I99\|SUCB2_HUMAN | SUCLG2 | Succinate--CoA ligase [GDP-forming] subunit beta, mitochondrial OS=Homo sapiens GN=SUCLG2 PE=1 SV=2 | 46.48 | 0 | 0 | 2 | 2 | NF | 2.30E+04 |
| sp\|P61619\|S61A1_HUMAN | SEC61A1 | Protein transport protein Sec61 subunit alpha isoform 1 OS=Homo sapiens GN=SEC61A1 PE=1 SV=2 | 52.23 | 0 | 0 | 2 | 2 | NF | 4.80E+04 |
| sp\|P24752\|THIL_HUMAN | ACAT1 | Acetyl-CoA acetyltransferase, mitochondrial OS=Homo sapiens GN=ACAT1 PE=1 SV=1 | 45.17 | 0 | 0 | 1 | 2 | NF | 9.80E+04 |
| sp\|P61247\|RS3A_HUMAN | RPS3A | 40S ribosomal protein S3a OS=Homo sapiens GN=RPS3A PE=1 SV=2 | 29.93 | 0 | 0 | 1 | 2 | NF | 6.40E+04 |
| sp\|P62829\|RL23_HUMAN | RPL23 | 60S ribosomal protein L23 OS=Homo sapiens GN=RPL23 PE=1 SV=1 | 14.86 | 0 | 0 | 1 | 2 | NF | 8.60E+04 |
| sp\|P18085\|ARF4_HUMAN | ARF4 | ADP-ribosylation factor 4 OS=Homo sapiens GN=ARF4 PE=1 SV=3 | 20.5 | 0 | 0 | 1 | 2 | NF | 9.20E+04 |
| sp\|P62820\|RAB1A_HUMAN | RAB1A | Ras-related protein Rab-1A OS=Homo sapiens GN=RAB1A PE=1 SV=3 | 22.66 | 0 | 0 | 1 | 2 | NF | 3.70E+04 |
| sp\|O15258\|RER1_HUMAN | RER1 | Protein RER1 OS=Homo sapiens GN=RER1 PE=1 SV=1 | 22.94 | 0 | 0 | 1 | 2 | NF | 5.40E+04 |
| sp\|O15427\|MOT4_HUMAN | SLC16A3 | Monocarboxylate transporter 4 OS=Homo sapiens GN=SLC16A3 PE=1 SV=1 | 49.44 | 0 | 0 | 1 | 2 | NF | 1.60E+05 |
| sp\|P62857\|RS28_HUMAN | RPS28 | 40S ribosomal protein S28 OS=Homo sapiens GN=RPS28 PE=1 SV=1 | 7.84 | 0 | 0 | 1 | 2 | NF | 1.30E+05 |
| sp\|Q92643\|GPI8_HUMAN | PIGK | GPI-anchor transamidase OS=Homo sapiens GN=PIGK PE=1 SV=2 | 45.22 | 0 | 0 | 1 | 2 | NF | 6.60E+04 |
| sp\|Q8IV08\|PLD3_HUMAN | PLD3 | Phospholipase D3 OS=Homo sapiens GN=PLD3 PE=1 SV=1 | 54.67 | 0 | 0 | 1 | 2 | NF | 2.10E+04 |
| sp\|P43307\|SSRA_HUMAN | SSR1 | Translocon-associated protein subunit alpha OS=Homo sapiens GN=SSR1 PE=1 SV=3 | 32.22 | 0 | 0 | 1 | 2 | NF | 1.40E+05 |
| sp\|Q9BRK5\|CAB45_HUMAN | SDF4 | 45 kDa calcium-binding protein OS=Homo sapiens GN=SDF4 PE=1 SV=1 | 41.78 | 0 | 0 | 1 | 2 | NF | 5.00E+04 |
| sp\|Q9BT22\|ALG1_HUMAN | ALG1 | Chitobiosyldiphosphodolichol beta-mannosyltransferase OS=Homo sapiens GN=ALG1 PE=1 SV=2 | 52.48 | 0 | 0 | 1 | 2 | NF | 6.70E+04 |
| sp\|P12004\|PCNA_HUMAN | PCNA | Proliferating cell nuclear antigen OS=Homo sapiens GN=PCNA PE=1 SV=1 | 28.75 | 0 | 0 | 1 | 2 | NF | 6.80E+04 |
| sp\|O75396\|SC22B_HUMAN | SEC22B | Vesicle-trafficking protein SEC22b OS=Homo sapiens GN=SEC22B PE=1 SV=4 | 24.58 | 0 | 0 | 1 | 2 | NF | 9.00E+04 |
| sp\|Q9HCU5\|PREB_HUMAN | PREB | Prolactin regulatory element-binding protein OS=Homo sapiens GN=PREB PE=1 SV=2 | 45.44 | 0 | 0 | 1 | 2 | NF | 4.60E+04 |
| sp\|Q9UM00\|TMCO1_HUMAN | TMCO1 | Calcium load-activated calcium channel OS=Homo sapiens GN=TMCO1 PE=1 SV=1 | 21.16 | 0 | 0 | 1 | 2 | NF | 6.90E+04 |
| sp\|Q8N0U8\|VKORL_HUMAN | VKORC1L1 | Vitamin K epoxide reductase complex subunit 1-like protein 1 OS=Homo sapiens GN=VKORC1L1 PE=1 SV=2 | 19.82 | 0 | 0 | 1 | 2 | NF | 3.50E+04 |
| sp\|Q9NUS5\|AP5S1_HUMAN | AP5S1 | AP-5 complex subunit sigma-1 OS=Homo sapiens GN=AP5S1 PE=1 SV=1 | 22.51 | 0 | 0 | 1 | 2 | NF | 4.00E+04 |
| sp\|P07858\|CATB_HUMAN | CTSB | Cathepsin B OS=Homo sapiens GN=CTSB PE=1 SV=3 | 37.8 | 0 | 0 | 1 | 2 | NF | 4.50E+04 |
| sp\|Q9Y6N7\|ROBO1_HUMAN | ROBO1 | Roundabout homolog 1 OS=Homo sapiens GN=ROBO1 PE=1 SV=1 | 180.82 | 0 | 0 | 1 | 2 | NF | 1.30E+04 |
| sp\|Q8N1F7\|NUP93_HUMAN | NUP93 | Nuclear pore complex protein Nup93 OS=Homo sapiens GN=NUP93 PE=1 SV=2 | 93.43 | 0 | 0 | 1 | 2 | NF | 1.50E+04 |
| sp\|Q70J99\|UN13D_HUMAN | UNC13D | Protein unc-13 homolog D OS=Homo sapiens GN=UNC13D PE=1 SV=1 | 123.2 | 0 | 0 | 1 | 2 | NF | 8.20E+03 |
| sp\|P61313\|RL15_HUMAN | RPL15 | 60S ribosomal protein L15 OS=Homo sapiens GN=RPL15 PE=1 SV=2 | 24.13 | 0 | 0 | 1 | 2 | NF | 4.30E+04 |
| sp\|Q9Y276\|BCS1_HUMAN | BCS1L | Mitochondrial chaperone BCS1 OS=Homo sapiens GN=BCS1L PE=1 SV=1 | 47.5 | 0 | 0 | 1 | 2 | NF | 5.60E+03 |
| sp\|Q6R327\|RICTR_HUMAN | RICTOR | Rapamycin-insensitive companion of mTOR OS=Homo sapiens GN=RICTOR PE=1 SV=1 | 192.1 | 0 | 0 | 1 | 2 | NF | 1.10E+04 |
| sp\|P62701\|RS4X_HUMAN | RPS4X | 40S ribosomal protein S4, X isoform OS=Homo sapiens GN=RPS4X PE=1 SV=2 | 29.58 | 0 | 0 | 1 | 2 | NF | 9.30E+04 |
| sp\|Q9BVC6\|TM109_HUMAN | TMEM109 | Transmembrane protein 109 OS=Homo sapiens GN=TMEM109 PE=1 SV=1 | 26.19 | 0 | 0 | 1 | 2 | NF | 3.30E+05 |
| sp\|O43852\|CALU_HUMAN | CALU | Calumenin OS=Homo sapiens GN=CALU PE=1 SV=2 | 37.08 | 0 | 0 | 1 | 2 | NF | 2.30E+04 |
| sp\|O14579\|COPE_HUMAN | COPE | Coatomer subunit epsilon OS=Homo sapiens GN=COPE PE=1 SV=3 | 34.46 | 0 | 0 | 1 | 2 | NF | 3.10E+04 |
| sp\|Q08380\|LG3BP_HUMAN | LGALS3BP | Galectin-3-binding protein OS=Homo sapiens GN=LGALS3BP PE=1 SV=1 | 65.29 | 0 | 0 | 1 | 2 | NF | 2.70E+04 |
| sp\|O43808\|PM34_HUMAN | SLC25A17 | Peroxisomal membrane protein PMP34 OS=Homo sapiens GN=SLC25A17 PE=1 SV=1 | 34.54 | 0 | 0 | 1 | 2 | NF | 2.20E+04 |
| sp\|Q9Y6K0\|CEPT1_HUMAN | CEPT1 | Choline/ethanolaminephosphotransferase 1 OS=Homo sapiens GN=CEPT1 PE=1 SV=1 | 46.52 | 0 | 0 | 1 | 2 | NF | 5.50E+04 |
| tr\|H3BM99\|H3BM99_HUMAN | MAP1LC3B | Microtubule-associated proteins 1A/1B light chain 3B OS=Homo sapiens GN=MAP1LC3B PE=4 SV=1 | 6.86 | 0 | 0 | 1 | 2 | NF | 1.50E+06 |
| sp\|Q15036\|SNX17_HUMAN | SNX17 | Sorting nexin-17 OS=Homo sapiens GN=SNX17 PE=1 SV=1 | 52.87 | 0 | 0 | 1 | 1 | NF | 6.50E+03 |
| sp\|P05023\|AT1A1_HUMAN | ATP1A1 | Sodium/potassium-transporting ATPase subunit alpha-1 OS=Homo sapiens GN=ATP1A1 PE=1 SV=1 | 112.82 | 0 | 0 | 1 | 1 | NF | 1.50E+04 |
| sp\|Q14126\|DSG2_HUMAN | DSG2 | Desmoglein-2 OS=Homo sapiens GN=DSG2 PE=1 SV=2 | 122.22 | 0 | 0 | 1 | 1 | NF | 1.90E+04 |
| sp\|Q04721\|NOTC2_HUMAN | NOTCH2 | Neurogenic locus notch homolog protein 2 OS=Homo sapiens GN=NOTCH2 PE=1 SV=3 | 265.23 | 0 | 0 | 1 | 1 | NF | 8.20E+03 |
| sp\|P09382\|LEG1_HUMAN | LGALS1 | Galectin-1 OS=Homo sapiens GN=LGALS1 PE=1 SV=2 | 14.71 | 0 | 0 | 1 | 1 | NF | 2.30E+04 |
| sp\|P51151\|RAB9A_HUMAN | RAB9A | Ras-related protein Rab-9A OS=Homo sapiens GN=RAB9A PE=1 SV=1 | 22.82 | 0 | 0 | 1 | 1 | NF | 8.70E+03 |
| sp\|P19438\|TNR1A_HUMAN | TNFRSF1A | Tumor necrosis factor receptor superfamily member 1A OS=Homo sapiens GN=TNFRSF1A PE=1 SV=1 | 50.46 | 0 | 0 | 1 | 1 | NF | 6.00E+03 |
| sp\|P51149\|RAB7A_HUMAN | RAB7A | Ras-related protein Rab-7a OS=Homo sapiens GN=RAB7A PE=1 SV=1 | 23.47 | 0 | 0 | 1 | 1 | NF | 1.20E+04 |
| sp\|P52569\|CTR2_HUMAN | SLC7A2 | Cationic amino acid transporter 2 OS=Homo sapiens GN=SLC7A2 PE=1 SV=2 | 71.63 | 0 | 0 | 1 | 1 | NF | 7.50E+04 |
| sp\|Q99653\|CHP1_HUMAN | CHP1 | Calcineurin B homologous protein 1 OS=Homo sapiens GN=CHP1 PE=1 SV=3 | 22.44 | 0 | 0 | 1 | 1 | NF | 4.10E+04 |
| sp\|Q9BXS4\|TMM59_HUMAN | TMEM59 | Transmembrane protein 59 OS=Homo sapiens GN=TMEM59 PE=1 SV=1 | 36.2 | 0 | 0 | 1 | 1 | NF | 8.30E+03 |
| sp\|Q96B67\|ARRD3_HUMAN | ARRDC3 | Arrestin domain-containing protein 3 OS=Homo sapiens GN=ARRDC3 PE=1 SV=1 | 46.37 | 0 | 0 | 1 | 1 | NF | 7.30E+03 |
| sp\|Q96AG4\|LRC59_HUMAN | LRRC59 | Leucine-rich repeat-containing protein 59 OS=Homo sapiens GN=LRRC59 PE=1 SV=1 | 34.91 | 0 | 0 | 1 | 1 | NF | 5.80E+04 |
| sp\|P25789\|PSA4_HUMAN | PSMA4 | Proteasome subunit alpha type-4 OS=Homo sapiens GN=PSMA4 PE=1 SV=1 | 29.47 | 0 | 0 | 1 | 1 | NF | 1.40E+04 |
| sp\|Q16795\|NDUA9_HUMAN | NDUFA9 | NADH dehydrogenase [ubiquinone] 1 alpha subcomplex subunit 9, mitochondrial OS=Homo sapiens GN=NDUFA9 PE=1 SV=2 | 42.48 | 0 | 0 | 1 | 1 | NF | 9.00E+03 |
| sp\|Q9NP58\|ABCB6_HUMAN | ABCB6 | ATP-binding cassette sub-family B member 6, mitochondrial OS=Homo sapiens GN=ABCB6 PE=1 SV=1 | 93.83 | 0 | 0 | 1 | 1 | NF | 1.40E+04 |
| sp\|P48643\|TCPE_HUMAN | CCT5 | T-complex protein 1 subunit epsilon OS=Homo sapiens GN=CCT5 PE=1 SV=1 | 59.63 | 0 | 0 | 1 | 1 | NF | 2.40E+04 |
| sp\|P00533\|EGFR_HUMAN | EGFR | Epidermal growth factor receptor OS=Homo sapiens GN=EGFR PE=1 SV=2 | 134.19 | 0 | 0 | 1 | 1 | NF | 8.10E+03 |
| sp\|P62333\|PRS10_HUMAN | PSMC6 | 26S protease regulatory subunit 10B OS=Homo sapiens GN=PSMC6 PE=1 SV=1 | 44.15 | 0 | 0 | 1 | 1 | NF | 1.20E+04 |
| sp\|P15311\|EZRI_HUMAN | EZR | Ezrin OS=Homo sapiens GN=EZR PE=1 SV=4 | 69.37 | 0 | 0 | 1 | 1 | NF | 2.30E+04 |
| sp\|Q8TB37\|NUBPL_HUMAN | NUBPL | Iron-sulfur protein NUBPL OS=Homo sapiens GN=NUBPL PE=1 SV=3 | 34.06 | 0 | 0 | 1 | 1 | NF | 1.10E+04 |
| sp\|P34932\|HSP74_HUMAN | HSPA4 | Heat shock 70 kDa protein 4 OS=Homo sapiens GN=HSPA4 PE=1 SV=4 | 94.27 | 0 | 0 | 1 | 1 | NF | 5.10E+03 |
| sp\|Q9BVX2\|T106C_HUMAN | TMEM106C | Transmembrane protein 106C OS=Homo sapiens GN=TMEM106C PE=1 SV=1 | 27.86 | 0 | 0 | 1 | 1 | NF | 1.50E+04 |
| sp\|Q8WUY1\|THEM6_HUMAN | THEM6 | Protein THEM6 OS=Homo sapiens GN=THEM6 PE=1 SV=2 | 23.85 | 0 | 0 | 1 | 1 | NF | 7.00E+03 |
| sp\|P54709\|AT1B3_HUMAN | ATP1B3 | Sodium/potassium-transporting ATPase subunit beta-3 OS=Homo sapiens GN=ATP1B3 PE=1 SV=1 | 31.49 | 0 | 0 | 1 | 1 | NF | 2.40E+04 |
| sp\|Q9Y3D0\|MIP18_HUMAN | FAM96B | Mitotic spindle-associated MMXD complex subunit MIP18 OS=Homo sapiens GN=FAM96B PE=1 SV=1 | 17.65 | 0 | 0 | 1 | 1 | NF | 2.50E+04 |
| sp\|P61026\|RAB10_HUMAN | RAB10 | Ras-related protein Rab-10 OS=Homo sapiens GN=RAB10 PE=1 SV=1 | 22.53 | 0 | 0 | 1 | 1 | NF | 2.90E+04 |
| sp\|O75509\|TNR21_HUMAN | TNFRSF21 | Tumor necrosis factor receptor superfamily member 21 OS=Homo sapiens GN=TNFRSF21 PE=1 SV=1 | 71.8 | 0 | 0 | 1 | 1 | NF | 6.10E+03 |
| sp\|Q6P3X3\|TTC27_HUMAN | TTC27 | Tetratricopeptide repeat protein 27 OS=Homo sapiens GN=TTC27 PE=1 SV=1 | 96.57 | 0 | 0 | 1 | 1 | NF | 1.50E+04 |
| sp\|P49755\|TMEDA_HUMAN | TMED10 | Transmembrane emp24 domain-containing protein 10 OS=Homo sapiens GN=TMED10 PE=1 SV=2 | 24.96 | 0 | 0 | 1 | 1 | NF | 2.10E+04 |
| sp\|Q96FQ6\|S10AG_HUMAN | S100A16 | Protein S100-A16 OS=Homo sapiens GN=S100A16 PE=1 SV=1 | 11.79 | 0 | 0 | 1 | 1 | NF | 1.20E+04 |
| sp\|Q7L9L4\|MOB1B_HUMAN | MOB1B | MOB kinase activator 1B OS=Homo sapiens GN=MOB1B PE=1 SV=3 | 25.07 | 0 | 0 | 1 | 1 | NF | 1.00E+04 |
| sp\|Q12965\|MYO1E_HUMAN | MYO1E | Unconventional myosin-Ie OS=Homo sapiens GN=MYO1E PE=1 SV=2 | 126.98 | 0 | 0 | 1 | 1 | NF | 1.30E+04 |
| sp\|P55795\|HNRH2_HUMAN | HNRNPH2 | Heterogeneous nuclear ribonucleoprotein H2 OS=Homo sapiens GN=HNRNPH2 PE=1 SV=1 | 49.23 | 0 | 0 | 1 | 1 | NF | 2.40E+04 |
| sp\|P24390\|ERD21_HUMAN | KDELR1 | ER lumen protein-retaining receptor 1 OS=Homo sapiens GN=KDELR1 PE=1 SV=1 | 24.53 | 0 | 0 | 1 | 1 | NF | 2.50E+04 |
| sp\|P31946\|1433B_HUMAN | YWHAB | 14-3-3 protein beta/alpha OS=Homo sapiens GN=YWHAB PE=1 SV=3 | 28.06 | 0 | 0 | 1 | 1 | NF | 1.00E+04 |
| sp\|P24534\|EF1B_HUMAN | EEF1B2 | Elongation factor 1-beta OS=Homo sapiens GN=EEF1B2 PE=1 SV=3 | 24.75 | 0 | 0 | 1 | 1 | NF | 2.60E+04 |
| sp\|P37802\|TAGL2_HUMAN | TAGLN2 | Transgelin-2 OS=Homo sapiens GN=TAGLN2 PE=1 SV=3 | 22.38 | 0 | 0 | 1 | 1 | NF | 7.00E+03 |
| sp\|P17813\|EGLN_HUMAN | ENG | Endoglin OS=Homo sapiens GN=ENG PE=1 SV=2 | 70.53 | 0 | 0 | 1 | 1 | NF | 1.80E+04 |
| sp\|P43246\|MSH2_HUMAN | MSH2 | DNA mismatch repair protein Msh2 OS=Homo sapiens GN=MSH2 PE=1 SV=1 | 104.68 | 0 | 0 | 1 | 1 | NF | 2.00E+04 |
| sp\|Q96KA5\|CLP1L_HUMAN | CLPTM1L | Cleft lip and palate transmembrane protein 1-like protein OS=Homo sapiens GN=CLPTM1L PE=1 SV=1 | 62.19 | 0 | 0 | 1 | 1 | NF | 1.80E+04 |
| sp\|P27348\|1433T_HUMAN | YWHAQ | 14-3-3 protein theta OS=Homo sapiens GN=YWHAQ PE=1 SV=1 | 27.75 | 0 | 0 | 1 | 1 | NF | 7.90E+03 |
| sp\|Q6AI08\|HEAT6_HUMAN | HEATR6 | HEAT repeat-containing protein 6 OS=Homo sapiens GN=HEATR6 PE=1 SV=1 | 128.7 | 0 | 0 | 1 | 1 | NF | 8.80E+03 |
| sp\|Q02878\|RL6_HUMAN | RPL6 | 60S ribosomal protein L6 OS=Homo sapiens GN=RPL6 PE=1 SV=3 | 32.71 | 0 | 0 | 1 | 1 | NF | 3.20E+04 |
| sp\|Q96P70\|IPO9_HUMAN | IPO9 | Importin-9 OS=Homo sapiens GN=IPO9 PE=1 SV=3 | 115.89 | 0 | 0 | 1 | 1 | NF | 3.70E+04 |
| sp\|O95433\|AHSA1_HUMAN | AHSA1 | Activator of 90 kDa heat shock protein ATPase homolog 1 OS=Homo sapiens GN=AHSA1 PE=1 SV=1 | 38.25 | 0 | 0 | 1 | 1 | NF | 2.30E+04 |
| sp\|P53999\|TCP4_HUMAN | SUB1 | Activated RNA polymerase II transcriptional coactivator p15 OS=Homo sapiens GN=SUB1 PE=1 SV=3 | 14.39 | 0 | 0 | 1 | 1 | NF | 3.60E+04 |
| sp\|Q14118\|DAG1_HUMAN | DAG1 | Dystroglycan OS=Homo sapiens GN=DAG1 PE=1 SV=2 | 97.38 | 0 | 0 | 1 | 1 | NF | 5.30E+04 |
| sp\|Q6NTF9\|RHBD2_HUMAN | RHBDD2 | Rhomboid domain-containing protein 2 OS=Homo sapiens GN=RHBDD2 PE=2 SV=2 | 39.18 | 0 | 0 | 1 | 1 | NF | 1.50E+04 |
| sp\|O75352\|MPU1_HUMAN | MPDU1 | Mannose-P-dolichol utilization defect 1 protein OS=Homo sapiens GN=MPDU1 PE=1 SV=2 | 26.62 | 0 | 0 | 1 | 1 | NF | 1.00E+04 |
| sp\|P23246\|SFPQ_HUMAN | SFPQ | Splicing factor, proline- and glutamine-rich OS=Homo sapiens GN=SFPQ PE=1 SV=2 | 76.1 | 0 | 0 | 1 | 1 | NF | 3.00E+04 |
| sp\|Q92882\|OSTF1_HUMAN | OSTF1 | Osteoclast-stimulating factor 1 OS=Homo sapiens GN=OSTF1 PE=1 SV=2 | 23.77 | 0 | 0 | 1 | 1 | NF | 7.90E+03 |
| sp\|Q9Y657\|SPIN1_HUMAN | SPIN1 | Spindlin-1 OS=Homo sapiens GN=SPIN1 PE=1 SV=3 | 29.58 | 0 | 0 | 1 | 1 | NF | 7.40E+03 |
| sp\|Q15477\|SKIV2_HUMAN | SKIV2L | Helicase SKI2W OS=Homo sapiens GN=SKIV2L PE=1 SV=3 | 137.67 | 0 | 0 | 1 | 1 | NF | 4.90E+03 |

(NF: not found)

| Reference | Gene Symbol | Annotation | MWT(kDa) | Unique FA | Total FA | Unique JX4-FA | Total JX4-FA | Sum Intensity FA | Sum Intensity JX4-FA |
|---|---|---|---|---|---|---|---|---|---|
| sp\|P24539\|AT5F1_HUMAN | ATP5F1 | ATP synthase F(0) complex subunit B1, mitochondrial OS=Homo sapiens GN=ATP5F1 PE=1 SV=2 | 28.89 | 0 | 0 | 1 | 1 | NF | 2.10E+04 |
| sp\|O14735\|CDIPT_HUMAN | CDIPT | CDP-diacylglycerol--inositol 3-phosphatidyltransferase OS=Homo sapiens GN=CDIPT PE=1 SV=1 | 23.52 | 0 | 0 | 1 | 1 | NF | 2.50E+04 |
| sp\|P62266\|RS23_HUMAN | RPS23 | 40S ribosomal protein S23 OS=Homo sapiens GN=RPS23 PE=1 SV=3 | 15.8 | 0 | 0 | 1 | 1 | NF | 3.50E+04 |
| sp\|Q8TEM1\|PO210_HUMAN | NUP210 | Nuclear pore membrane glycoprotein 210 OS=Homo sapiens GN=NUP210 PE=1 SV=3 | 204.98 | 0 | 0 | 1 | 1 | NF | 1.00E+04 |
| sp\|Q9Y678\|COPG1_HUMAN | COPG1 | Coatomer subunit gamma-1 OS=Homo sapiens GN=COPG1 PE=1 SV=1 | 97.66 | 0 | 0 | 1 | 1 | NF | 3.00E+04 |
| sp\|P00966\|ASSY_HUMAN | ASS1 | Argininosuccinate synthase OS=Homo sapiens GN=ASS1 PE=1 SV=2 | 46.5 | 0 | 0 | 1 | 1 | NF | 1.10E+04 |
| sp\|Q8WUK0\|PTPM1_HUMAN | PTPMT1 | Phosphatidylglycerophosphatase and protein-tyrosine phosphatase 1 OS=Homo sapiens GN=PTPMT1 PE=1 SV=1 | 22.83 | 0 | 0 | 1 | 1 | NF | 5.10E+04 |
| sp\|Q5JU69\|TOR2A_HUMAN | TOR2A | Torsin-2A OS=Homo sapiens GN=TOR2A PE=2 SV=1 | 35.69 | 0 | 0 | 1 | 1 | NF | 1.80E+04 |
| sp\|Q15366\|PCBP2_HUMAN | PCBP2 | Poly(rC)-binding protein 2 OS=Homo sapiens GN=PCBP2 PE=1 SV=1 | 38.56 | 0 | 0 | 1 | 1 | NF | 2.40E+04 |
| sp\|P11166\|GTR1_HUMAN | SLC2A1 | Solute carrier family 2, facilitated glucose transporter member 1 OS=Homo sapiens GN=SLC2A1 PE=1 SV=2 | 54.05 | 0 | 0 | 1 | 1 | NF | 1.50E+04 |
| sp\|P49721\|PSB2_HUMAN | PSMB2 | Proteasome subunit beta type-2 OS=Homo sapiens GN=PSMB2 PE=1 SV=1 | 22.82 | 0 | 0 | 1 | 1 | NF | 1.50E+04 |
| sp\|Q8TF09\|DLRB2_HUMAN | DYNLRB2 | Dynein light chain roadblock-type 2 OS=Homo sapiens GN=DYNLRB2 PE=1 SV=1 | 10.85 | 0 | 0 | 1 | 1 | NF | 2.30E+04 |
| sp\|P55084\|ECHB_HUMAN | HADHB | Trifunctional enzyme subunit beta, mitochondrial OS=Homo sapiens GN=HADHB PE=1 SV=3 | 51.26 | 0 | 0 | 1 | 1 | NF | 8.50E+03 |
| sp\|Q9Y3D6\|FIS1_HUMAN | FIS1 | Mitochondrial fission 1 protein OS=Homo sapiens GN=FIS1 PE=1 SV=2 | 16.93 | 0 | 0 | 1 | 1 | NF | 3.80E+04 |
| sp\|P51809\|VAMP7_HUMAN | VAMP7 | Vesicle-associated membrane protein 7 OS=Homo sapiens GN=VAMP7 PE=1 SV=3 | 24.92 | 0 | 0 | 1 | 1 | NF | 1.70E+04 |
| sp\|Q9BSJ8\|ESYT1_HUMAN | ESYT1 | Extended synaptotagmin-1 OS=Homo sapiens GN=ESYT1 PE=1 SV=1 | 122.78 | 0 | 0 | 1 | 1 | NF | 5.60E+03 |
| sp\|Q16718\|NDUA5_HUMAN | NDUFA5 | NADH dehydrogenase [ubiquinone] 1 alpha subcomplex subunit 5 OS=Homo sapiens GN=NDUFA5 PE=1 SV=3 | 13.45 | 0 | 0 | 1 | 1 | NF | 7.60E+03 |
| tr\|E5KLJ5\|E5KLJ5_HUMAN | OPA1 | Dynamin-like 120 kDa protein, mitochondrial OS=Homo sapiens GN=OPA1 PE=1 SV=1 | 117.67 | 0 | 0 | 1 | 1 | NF | 4.70E+03 |
| sp\|Q96CS3\|FAF2_HUMAN | FAF2 | FAS-associated factor 2 OS=Homo sapiens GN=FAF2 PE=1 SV=2 | 52.59 | 0 | 0 | 1 | 1 | NF | 5.00E+03 |
| sp\|P62241\|RS8_HUMAN | RPS8 | 40S ribosomal protein S8 OS=Homo sapiens GN=RPS8 PE=1 SV=2 | 24.19 | 0 | 0 | 1 | 1 | NF | 3.30E+04 |
| sp\|Q96GQ5\|RUS1_HUMAN | C16orf58 | RUS1 family protein C16orf58 OS=Homo sapiens GN=C16orf58 PE=1 SV=2 | 50.99 | 0 | 0 | 1 | 1 | NF | 8.40E+03 |
| sp\|Q8NC51\|PAIRB_HUMAN | SERBP1 | Plasminogen activator inhibitor 1 RNA-binding protein OS=Homo sapiens GN=SERBP1 PE=1 SV=2 | 44.94 | 0 | 0 | 1 | 1 | NF | 8.60E+03 |
| sp\|Q6IAN0\|DRS7B_HUMAN | DHRS7B | Dehydrogenase/reductase SDR family member 7B OS=Homo sapiens GN=DHRS7B PE=1 SV=2 | 35.1 | 0 | 0 | 1 | 1 | NF | 2.10E+04 |
| sp\|P30048\|PRDX3_HUMAN | PRDX3 | Thioredoxin-dependent peroxide reductase, mitochondrial OS=Homo sapiens GN=PRDX3 PE=1 SV=3 | 27.68 | 0 | 0 | 1 | 1 | NF | 2.70E+04 |
| tr\|B2RWN5\|B2RWN5_HUMAN | HEATR1 | HEAT repeat containing 1 OS=Homo sapiens GN=HEATR1 PE=2 SV=1 | 242.11 | 0 | 0 | 1 | 1 | NF | 1.30E+04 |
| sp\|Q13509\|TBB3_HUMAN | TUBB3 | Tubulin beta-3 chain OS=Homo sapiens GN=TUBB3 PE=1 SV=2 | 50.4 | 0 | 0 | 1 | 1 | NF | 1.20E+04 |
| sp\|Q9NWW5\|CLN6_HUMAN | CLN6 | Ceroid-lipofuscinosis neuronal protein 6 OS=Homo sapiens GN=CLN6 PE=1 SV=1 | 35.9 | 0 | 0 | 1 | 1 | NF | 3.10E+04 |
| sp\|Q9UNL2\|SSRG_HUMAN | SSR3 | Translocon-associated protein subunit gamma OS=Homo sapiens GN=SSR3 PE=1 SV=1 | 21.07 | 0 | 0 | 1 | 1 | NF | 3.50E+04 |
| sp\|O60506\|HNRPQ_HUMAN | SYNCRIP | Heterogeneous nuclear ribonucleoprotein Q OS=Homo sapiens GN=SYNCRIP PE=1 SV=2 | 69.56 | 0 | 0 | 1 | 1 | NF | 9.80E+03 |
| sp\|Q14390\|GGTL2_HUMAN | GGTLC2 | Gamma-glutamyltransferase light chain 2 OS=Homo sapiens GN=GGTLC2 PE=2 SV=4 | 23.65 | 0 | 0 | 1 | 1 | NF | 3.60E+03 |
| sp\|Q14185\|DOCK1_HUMAN | DOCK1 | Dedicator of cytokinesis protein 1 OS=Homo sapiens GN=DOCK1 PE=1 SV=2 | 215.21 | 0 | 0 | 1 | 1 | NF | 1.20E+04 |
| sp\|Q9BXW9\|FACD2_HUMAN | FANCD2 | Fanconi anemia group D2 protein OS=Homo sapiens GN=FANCD2 PE=1 SV=2 | 164.02 | 0 | 0 | 1 | 1 | NF | 3.80E+03 |
| sp\|P49411\|EFTU_HUMAN | TUFM | Elongation factor Tu, mitochondrial OS=Homo sapiens GN=TUFM PE=1 SV=2 | 49.51 | 0 | 0 | 1 | 1 | NF | 6.60E+03 |
| sp\|Q15165\|PON2_HUMAN | PON2 | Serum paraoxonase/arylesterase 2 OS=Homo sapiens GN=PON2 PE=1 SV=3 | 39.37 | 0 | 0 | 1 | 1 | NF | 1.30E+04 |
| sp\|P62750\|RL23A_HUMAN | RPL23A | 60S ribosomal protein L23a OS=Homo sapiens GN=RPL23A PE=1 SV=1 | 17.68 | 0 | 0 | 1 | 1 | NF | 1.00E+04 |
| sp\|Q9H845\|ACAD9_HUMAN | ACAD9 | Acyl-CoA dehydrogenase family member 9, mitochondrial OS=Homo sapiens GN=ACAD9 PE=1 SV=1 | 68.72 | 0 | 0 | 1 | 1 | NF | 1.20E+04 |
| sp\|Q9UP83\|COG5_HUMAN | COG5 | Conserved oligomeric Golgi complex subunit 5 OS=Homo sapiens GN=COG5 PE=1 SV=3 | 92.68 | 0 | 0 | 1 | 1 | NF | 2.00E+04 |
| sp\|Q9Y2V7\|COG6_HUMAN | COG6 | Conserved oligomeric Golgi complex subunit 6 OS=Homo sapiens GN=COG6 PE=1 SV=2 | 73.23 | 0 | 0 | 1 | 1 | NF | 1.10E+04 |
| sp\|P0CG08\|GPHRB_HUMAN | GPR89B | Golgi pH regulator B OS=Homo sapiens GN=GPR89B PE=1 SV=1 | 52.88 | 0 | 0 | 1 | 1 | NF | 1.20E+04 |
| sp\|Q04837\|SSBP_HUMAN | SSBP1 | Single-stranded DNA-binding protein, mitochondrial OS=Homo sapiens GN=SSBP1 PE=1 SV=1 | 17.25 | 0 | 0 | 1 | 1 | NF | 9.30E+03 |
| sp\|Q9NU22\|MDN1_HUMAN | MDN1 | Midasin OS=Homo sapiens GN=MDN1 PE=1 SV=2 | 632.42 | 0 | 0 | 1 | 1 | NF | 3.00E+03 |
| sp\|P52907\|CAZA1_HUMAN | CAPZA1 | F-actin-capping protein subunit alpha-1 OS=Homo sapiens GN=CAPZA1 PE=1 SV=3 | 32.9 | 0 | 0 | 1 | 1 | NF | 6.40E+03 |
| sp\|Q6P1M0\|S27A4_HUMAN | SLC27A4 | Long-chain fatty acid transport protein 4 OS=Homo sapiens GN=SLC27A4 PE=1 SV=1 | 72.02 | 0 | 0 | 1 | 1 | NF | 1.30E+04 |
| sp\|Q643R3\|LPCT4_HUMAN | LPCAT4 | Lysophospholipid acyltransferase LPCAT4 OS=Homo sapiens GN=LPCAT4 PE=1 SV=1 | 57.18 | 0 | 0 | 1 | 1 | NF | 7.40E+03 |
| sp\|P46977\|STT3A_HUMAN | STT3A | Dolichyl-diphosphooligosaccharide--protein glycosyltransferase subunit STT3A OS=Homo sapiens GN=STT3A PE=1 SV=2 | 80.48 | 0 | 0 | 1 | 1 | NF | 1.50E+04 |
| sp\|Q13148\|TADBP_HUMAN | TARDBP | TAR DNA-binding protein 43 OS=Homo sapiens GN=TARDBP PE=1 SV=1 | 44.71 | 0 | 0 | 1 | 1 | NF | 6.10E+03 |
| sp\|P61289\|PSME3_HUMAN | PSME3 | Proteasome activator complex subunit 3 OS=Homo sapiens GN=PSME3 PE=1 SV=1 | 29.49 | 0 | 0 | 1 | 1 | NF | 5.30E+04 |
| sp\|Q96T76\|MMS19_HUMAN | MMS19 | MMS19 nucleotide excision repair protein homolog OS=Homo sapiens GN=MMS19 PE=1 SV=2 | 113.22 | 0 | 0 | 1 | 1 | NF | 1.40E+04 |
| sp\|Q86X83\|COMD2_HUMAN | COMMD2 | COMM domain-containing protein 2 OS=Homo sapiens GN=COMMD2 PE=1 SV=2 | 22.73 | 0 | 0 | 1 | 1 | NF | 1.60E+04 |
| sp\|Q9C0C9\|UBE2O_HUMAN | UBE2O | (E3-independent) E2 ubiquitin-conjugating enzyme OS=Homo sapiens GN=UBE2O PE=1 SV=3 | 141.21 | 0 | 0 | 1 | 1 | NF | 1.20E+04 |
| sp\|P83436\|COG7_HUMAN | COG7 | Conserved oligomeric Golgi complex subunit 7 OS=Homo sapiens GN=COG7 PE=1 SV=1 | 86.29 | 0 | 0 | 1 | 1 | NF | 8.50E+03 |
| sp\|Q13433\|S39A6_HUMAN | SLC39A6 | Zinc transporter ZIP6 OS=Homo sapiens GN=SLC39A6 PE=1 SV=3 | 84.99 | 0 | 0 | 1 | 1 | NF | 7.60E+03 |
| sp\|Q9P260\|K1468_HUMAN | KIAA1468 | LisH domain and HEAT repeat-containing protein KIAA1468 OS=Homo sapiens GN=KIAA1468 PE=1 SV=2 | 134.55 | 0 | 0 | 1 | 1 | NF | 1.20E+04 |
| sp\|Q96EY1\|DNJA3_HUMAN | DNAJA3 | DnaJ homolog subfamily A member 3, mitochondrial OS=Homo sapiens GN=DNAJA3 PE=1 SV=2 | 52.46 | 0 | 0 | 1 | 1 | NF | 1.30E+04 |
| sp\|Q29RF7\|PDS5A_HUMAN | PDS5A | Sister chromatid cohesion protein PDS5 homolog A OS=Homo sapiens GN=PDS5A PE=1 SV=1 | 150.73 | 0 | 0 | 1 | 1 | NF | 8.90E+03 |
| sp\|P39019\|RS19_HUMAN | RPS19 | 40S ribosomal protein S19 OS=Homo sapiens GN=RPS19 PE=1 SV=2 | 16.05 | 0 | 0 | 1 | 1 | NF | 3.00E+04 |
| sp\|O75153\|CLU_HUMAN | CLUH | Clustered mitochondria protein homolog OS=Homo sapiens GN=CLUH PE=1 SV=2 | 146.58 | 0 | 0 | 1 | 1 | NF | 6.50E+03 |
| sp\|Q8WTV0\|SCRB1_HUMAN | SCARB1 | Scavenger receptor class B member 1 OS=Homo sapiens GN=SCARB1 PE=1 SV=1 | 60.84 | 0 | 0 | 1 | 1 | NF | 1.10E+04 |
| sp\|O43156\|TTI1_HUMAN | TTI1 | TELO2-interacting protein 1 homolog OS=Homo sapiens GN=TTI1 PE=1 SV=3 | 121.99 | 0 | 0 | 1 | 1 | NF | 8.50E+03 |
| sp\|Q6P474\|PDXD2_HUMAN | PDXDC2P | Putative pyridoxal-dependent decarboxylase domain-containing protein 2 OS=Homo sapiens GN=PDXDC2P PE=5 SV=3 | 51.78 | 0 | 0 | 1 | 1 | NF | 6.60E+03 |
| sp\|O95831\|AIFM1_HUMAN | AIFM1 | Apoptosis-inducing factor 1, mitochondrial OS=Homo sapiens GN=AIFM1 PE=1 SV=1 | 66.86 | 0 | 0 | 1 | 1 | NF | 3.20E+04 |
| sp\|P49368\|TCPG_HUMAN | CCT3 | T-complex protein 1 subunit gamma OS=Homo sapiens GN=CCT3 PE=1 SV=4 | 60.5 | 0 | 0 | 1 | 1 | NF | 1.80E+04 |
| sp\|O76027\|ANXA9_HUMAN | ANXA9 | Annexin A9 OS=Homo sapiens GN=ANXA9 PE=1 SV=3 | 38.34 | 0 | 0 | 1 | 1 | NF | 1.60E+04 |
| sp\|P35606\|COPB2_HUMAN | COPB2 | Coatomer subunit beta' OS=Homo sapiens GN=COPB2 PE=1 SV=2 | 102.42 | 0 | 0 | 1 | 1 | NF | 1.10E+04 |
| sp\|P43005\|EAA3_HUMAN | SLC1A1 | Excitatory amino acid transporter 3 OS=Homo sapiens GN=SLC1A1 PE=1 SV=2 | 57.06 | 0 | 0 | 1 | 1 | NF | 1.50E+04 |
| sp\|P11940\|PABP1_HUMAN | PABPC1 | Polyadenylate-binding protein 1 OS=Homo sapiens GN=PABPC1 PE=1 SV=2 | 70.63 | 0 | 0 | 1 | 1 | NF | 1.10E+04 |
| sp\|P52790\|HXK3_HUMAN | HK3 | Hexokinase-3 OS=Homo sapiens GN=HK3 PE=1 SV=2 | 98.96 | 0 | 0 | 1 | 1 | NF | 1.90E+04 |
| sp\|Q96QU8\|XPO6_HUMAN | XPO6 | Exportin-6 OS=Homo sapiens GN=XPO6 PE=1 SV=1 | 128.8 | 0 | 0 | 1 | 1 | NF | 1.00E+04 |
| sp\|Q9Y5M8\|SRPRB_HUMAN | SRPRB | Signal recognition particle receptor subunit beta OS=Homo sapiens GN=SRPRB PE=1 SV=3 | 29.68 | 0 | 0 | 1 | 1 | NF | 2.90E+04 |
| sp\|Q9H490\|PIGU_HUMAN | PIGU | Phosphatidylinositol glycan anchor biosynthesis class U protein OS=Homo sapiens GN=PIGU PE=1 SV=3 | 50.02 | 0 | 0 | 1 | 1 | NF | 2.80E+03 |
| sp\|P17301\|ITA2_HUMAN | ITGA2 | Integrin alpha-2 OS=Homo sapiens GN=ITGA2 PE=1 SV=1 | 129.21 | 0 | 0 | 1 | 1 | NF | 8.10E+03 |
| sp\|P62244\|RS15A_HUMAN | RPS15A | 40S ribosomal protein S15a OS=Homo sapiens GN=RPS15A PE=1 SV=2 | 14.83 | 0 | 0 | 1 | 1 | NF | 9.50E+03 |
| sp\|P04114\|APOB_HUMAN | APOB | Apolipoprotein B-100 OS=Homo sapiens GN=APOB PE=1 SV=2 | 515.28 | 0 | 0 | 1 | 1 | NF | 1.60E+04 |
| sp\|Q6P582\|MZT2A_HUMAN | MZT2A | Mitotic-spindle organizing protein 2A OS=Homo sapiens GN=MZT2A PE=1 SV=2 | 16.21 | 0 | 0 | 1 | 1 | NF | 3.90E+03 |
| sp\|Q96JJ3\|ELMO2_HUMAN | ELMO2 | Engulfment and cell motility protein 2 OS=Homo sapiens GN=ELMO2 PE=1 SV=2 | 82.56 | 0 | 0 | 1 | 1 | NF | 1.50E+04 |
| sp\|Q6YHU6\|THADA_HUMAN | THADA | Thyroid adenoma-associated protein OS=Homo sapiens GN=THADA PE=1 SV=1 | 219.47 | 0 | 0 | 1 | 1 | NF | 6.70E+03 |
| sp\|P49748\|ACADV_HUMAN | ACADVL | Very long-chain specific acyl-CoA dehydrogenase, mitochondrial OS=Homo sapiens GN=ACADVL PE=1 SV=1 | 70.35 | 0 | 0 | 1 | 1 | NF | 1.50E+04 |
| sp\|Q9NPI8\|FANCF_HUMAN | FANCF | Fanconi anemia group F protein OS=Homo sapiens GN=FANCF PE=1 SV=1 | 42.23 | 0 | 0 | 1 | 1 | NF | 1.40E+04 |
| sp\|Q9BQ95\|ECSIT_HUMAN | ECSIT | Evolutionarily conserved signaling intermediate in Toll pathway, mitochondrial OS=Homo sapiens GN=ECSIT PE=1 SV=1 | 49.12 | 0 | 0 | 1 | 1 | NF | 5.20E+03 |
| sp\|P62753\|RS6_HUMAN | RPS6 | 40S ribosomal protein S6 OS=Homo sapiens GN=RPS6 PE=1 SV=1 | 28.66 | 0 | 0 | 1 | 1 | NF | 2.90E+04 |
| sp\|O15360\|FANCA_HUMAN | FANCA | Fanconi anemia group A protein OS=Homo sapiens GN=FANCA PE=1 SV=2 | 162.67 | 0 | 0 | 1 | 1 | NF | 5.10E+03 |
| sp\|P43007\|SATT_HUMAN | SLC1A4 | Neutral amino acid transporter A OS=Homo sapiens GN=SLC1A4 PE=1 SV=1 | 55.69 | 0 | 0 | 1 | 1 | NF | 3.80E+04 |
| sp\|P31153\|METK2_HUMAN | MAT2A | S-adenosylmethionine synthase isoform type-2 OS=Homo sapiens GN=MAT2A PE=1 SV=1 | 43.63 | 0 | 0 | 1 | 1 | NF | 1.20E+04 |
| sp\|Q7Z4Q2\|HEAT3_HUMAN | HEATR3 | HEAT repeat-containing protein 3 OS=Homo sapiens GN=HEATR3 PE=1 SV=2 | 74.53 | 0 | 0 | 1 | 1 | NF | 1.20E+04 |
| sp\|Q9UKM7\|MA1B1_HUMAN | MAN1B1 | Endoplasmic reticulum mannosyl-oligosaccharide 1,2-alpha-mannosidase OS=Homo sapiens GN=MAN1B1 PE=1 SV=2 | 79.53 | 0 | 0 | 1 | 1 | NF | 7.00E+03 |
| sp\|Q2NL82\|TSR1_HUMAN | TSR1 | Pre-rRNA-processing protein TSR1 homolog OS=Homo sapiens GN=TSR1 PE=1 SV=1 | 91.75 | 0 | 0 | 1 | 1 | NF | 1.30E+04 |
| sp\|Q99832\|TCPH_HUMAN | CCT7 | T-complex protein 1 subunit eta OS=Homo sapiens GN=CCT7 PE=1 SV=2 | 59.33 | 0 | 0 | 1 | 1 | NF | 2.80E+04 |
| sp\|P33527\|MRP1_HUMAN | ABCC1 | Multidrug resistance-associated protein 1 OS=Homo sapiens GN=ABCC1 PE=1 SV=3 | 171.48 | 0 | 0 | 1 | 1 | NF | 1.20E+04 |
| sp\|Q9Y312\|AAR2_HUMAN | AAR2 | Protein AAR2 homolog OS=Homo sapiens GN=AAR2 PE=1 SV=2 | 43.44 | 0 | 0 | 1 | 1 | NF | 5.90E+03 |
| sp\|Q6RW13\|ATRAP_HUMAN | AGTRAP | Type-1 angiotensin II receptor-associated protein OS=Homo sapiens GN=AGTRAP PE=1 SV=1 | 17.41 | 0 | 0 | 1 | 1 | NF | 6.70E+03 |
| sp\|P35268\|RL22_HUMAN | RPL22 | 60S ribosomal protein L22 OS=Homo sapiens GN=RPL22 PE=1 SV=2 | 14.78 | 0 | 0 | 1 | 1 | NF | 1.40E+05 |
| sp\|P46781\|RS9_HUMAN | RPS9 | 40S ribosomal protein S9 OS=Homo sapiens GN=RPS9 PE=1 SV=3 | 22.58 | 0 | 0 | 1 | 1 | NF | 1.60E+04 |
| sp\|P62913\|RL11_HUMAN | RPL11 | 60S ribosomal protein L11 OS=Homo sapiens GN=RPL11 PE=1 SV=2 | 20.24 | 0 | 0 | 1 | 1 | NF | 1.40E+04 |
| sp\|Q969N2\|PIGT_HUMAN | PIGT | GPI transamidase component PIG-T OS=Homo sapiens GN=PIGT PE=1 SV=1 | 65.66 | 0 | 0 | 1 | 1 | NF | 8.50E+03 |
| tr\|H7C087\|H7C087_HUMAN | ZCWPW2 | Zinc finger CW-type PWWP domain protein 2 (Fragment) OS=Homo sapiens GN=ZCWPW2 PE=4 SV=1 | 19.43 | 0 | 0 | 1 | 1 | NF | 2.20E+04 |
| sp\|Q58FF7\|H90B3_HUMAN | HSP90AB3P | Putative heat shock protein HSP 90-beta-3 OS=Homo sapiens GN=HSP90AB3P PE=5 SV=1 | 68.28 | 0 | 0 | 1 | 1 | NF | 1.90E+04 |
| tr\|Q9NZ23\|Q9NZ23_HUMAN | YA61 | Drug-sensitive protein 1 OS=Homo sapiens GN=YA61 PE=2 SV=1 | 14.86 | 0 | 0 | 1 | 1 | NF | 3.20E+04 |
| sp\|P08237\|PFKAM_HUMAN | PFKM | ATP-dependent 6-phosphofructokinase, muscle type OS=Homo sapiens GN=PFKM PE=1 SV=2 | 85.13 | 0 | 0 | 1 | 1 | NF | 1.30E+04 |
| sp\|Q5VIR6\|VPS53_HUMAN | VPS53 | Vacuolar protein sorting-associated protein 53 homolog OS=Homo sapiens GN=VPS53 PE=1 SV=1 | 79.6 | 0 | 0 | 1 | 1 | NF | 1.10E+04 |
| sp\|O00217\|NDUS8_HUMAN | NDUFS8 | NADH dehydrogenase [ubiquinone] iron-sulfur protein 8, mitochondrial OS=Homo sapiens GN=NDUFS8 PE=1 SV=1 | 23.69 | 0 | 0 | 1 | 1 | NF | 5.00E+03 |
| sp\|Q7L8W6\|DPH6_HUMAN | DPH6 | Diphthine--ammonia ligase OS=Homo sapiens GN=DPH6 PE=1 SV=3 | 30.29 | 0 | 0 | 1 | 1 | NF | 9.10E+03 |

(NF: not found)

| Reference | Gene Symbol | Annotation | MWT(kDa) | Unique FA | Total FA | Unique JX4-FA | Total JX4-FA | Sum Intensity FA | Sum Intensity JX4-FA |
|---|---|---|---|---|---|---|---|---|---|
| sp\|Q16637\|SMN_HUMAN | SMN1 | Survival motor neuron protein OS=Homo sapiens GN=SMN1 PE=1 SV=1 | 31.83 | 0 | 0 | 1 | 1 | NF | 8.10E+03 |
| sp\|P14625\|ENPL_HUMAN | HSP90B1 | Endoplasmin OS=Homo sapiens GN=HSP90B1 PE=1 SV=1 | 92.41 | 0 | 0 | 1 | 1 | NF | 8.40E+03 |
| sp\|Q99943\|PLCA_HUMAN | AGPAT1 | 1-acyl-sn-glycerol-3-phosphate acyltransferase alpha OS=Homo sapiens GN=AGPAT1 PE=1 SV=2 | 31.7 | 0 | 0 | 1 | 1 | NF | 5.00E+03 |
| sp\|Q13162\|PRDX4_HUMAN | PRDX4 | Peroxiredoxin-4 OS=Homo sapiens GN=PRDX4 PE=1 SV=1 | 30.52 | 0 | 0 | 1 | 1 | NF | 2.70E+04 |
| sp\|Q13671\|RIN1_HUMAN | RIN1 | Ras and Rab interactor 1 OS=Homo sapiens GN=RIN1 PE=1 SV=4 | 84.05 | 0 | 0 | 1 | 1 | NF | 3.40E+03 |
| sp\|O00165\|HAX1_HUMAN | HAX1 | HCLS1-associated protein X-1 OS=Homo sapiens GN=HAX1 PE=1 SV=2 | 31.6 | 0 | 0 | 1 | 1 | NF | 9.90E+03 |
| sp\|P55010\|IF5_HUMAN | EIF5 | Eukaryotic translation initiation factor 5 OS=Homo sapiens GN=EIF5 PE=1 SV=2 | 49.19 | 0 | 0 | 1 | 1 | NF | 1.30E+04 |
| sp\|Q9BSJ2\|GCP2_HUMAN | TUBGCP2 | Gamma-tubulin complex component 2 OS=Homo sapiens GN=TUBGCP2 PE=1 SV=2 | 102.47 | 0 | 0 | 1 | 1 | NF | 1.10E+04 |
| sp\|Q15652\|JHD2C_HUMAN | JMJD1C | Probable JmjC domain-containing histone demethylation protein 2C OS=Homo sapiens GN=JMJD1C PE=1 SV=2 | 284.35 | 0 | 0 | 1 | 1 | NF | 2.10E+04 |
| sp\|P28340\|DPOD1_HUMAN | POLD1 | DNA polymerase delta catalytic subunit OS=Homo sapiens GN=POLD1 PE=1 SV=2 | 123.55 | 0 | 0 | 1 | 1 | NF | 8.20E+03 |
| sp\|Q14103\|HNRPD_HUMAN | HNRNPD | Heterogeneous nuclear ribonucleoprotein D0 OS=Homo sapiens GN=HNRNPD PE=1 SV=1 | 38.41 | 0 | 0 | 1 | 1 | NF | 1.20E+04 |
| sp\|P04792\|HSPB1_HUMAN | HSPB1 | Heat shock protein beta-1 OS=Homo sapiens GN=HSPB1 PE=1 SV=2 | 22.77 | 1 | 1 | 8 | 18 | 8.40E+03 | 2.40E+06 |
| sp\|P62987\|RL40_HUMAN | UBA52 | Ubiquitin-60S ribosomal protein L40 OS=Homo sapiens GN=UBA52 PE=1 SV=2 | 14.72 | 1 | 1 | 1 | 6 | 4.20E+04 | 2.60E+06 |
| sp\|P62736\|ACTA_HUMAN | ACTA2 | Actin, aortic smooth muscle OS=Homo sapiens GN=ACTA2 PE=1 SV=1 | 41.98 | 1 | 1 | 2 | 4 | 5.80E+04 | 2.30E+05 |
| sp\|P30050\|RL12_HUMAN | RPL12 | 60S ribosomal protein L12 OS=Homo sapiens GN=RPL12 PE=1 SV=1 | 17.81 | 1 | 1 | 3 | 4 | 7.10E+03 | 2.10E+05 |
| sp\|P63173\|RL38_HUMAN | RPL38 | 60S ribosomal protein L38 OS=Homo sapiens GN=RPL38 PE=1 SV=2 | 8.21 | 1 | 1 | 2 | 4 | 1.50E+04 | 1.70E+05 |
| sp\|P26641\|EF1G_HUMAN | EEF1G | Elongation factor 1-gamma OS=Homo sapiens GN=EEF1G PE=1 SV=3 | 50.09 | 1 | 1 | 2 | 4 | 4.20E+03 | 1.60E+05 |
| sp\|P27824\|CALX_HUMAN | CANX | Calnexin OS=Homo sapiens GN=CANX PE=1 SV=2 | 67.53 | 1 | 1 | 2 | 3 | 4.70E+03 | 6.20E+04 |
| sp\|P63104\|1433Z_HUMAN | YWHAZ | 14-3-3 protein zeta/delta OS=Homo sapiens GN=YWHAZ PE=1 SV=1 | 27.73 | 1 | 1 | 2 | 3 | 4.40E+03 | 7.00E+04 |
| sp\|P46783\|RS10_HUMAN | RPS10 | 40S ribosomal protein S10 OS=Homo sapiens GN=RPS10 PE=1 SV=1 | 18.89 | 1 | 1 | 2 | 3 | 1.10E+04 | 1.30E+05 |
| sp\|P18124\|RL7_HUMAN | RPL7 | 60S ribosomal protein L7 OS=Homo sapiens GN=RPL7 PE=1 SV=1 | 29.21 | 1 | 1 | 2 | 3 | 8.10E+03 | 1.50E+05 |
| sp\|P23396\|RS3_HUMAN | RPS3 | 60S ribosomal protein L7 OS=Homo sapiens GN=RPL7 PE=1 SV=1 | 26.67 | 1 | 1 | 2 | 3 | 6.80E+03 | 1.00E+05 |
| sp\|P05388\|RLA0_HUMAN | RPLP0 | 60S acidic ribosomal protein P0 OS=Homo sapiens GN=RPLP0 PE=1 SV=1 | 34.25 | 1 | 1 | 2 | 3 | 7.10E+03 | 1.40E+05 |
| sp\|P62249\|RS16_HUMAN | RPS16 | 40S ribosomal protein S16 OS=Homo sapiens GN=RPS16 PE=1 SV=2 | 16.44 | 1 | 1 | 2 | 3 | 9.50E+03 | 1.90E+05 |
| sp\|Q15084\|PDIA6_HUMAN | PDIA6 | Protein disulfide-isomerase A6 OS=Homo sapiens GN=PDIA6 PE=1 SV=1 | 48.09 | 1 | 1 | 1 | 2 | 4.90E+03 | 3.90E+04 |
| sp\|Q32P51\|RA1L2_HUMAN | HNRNPA1L2 | Heterogeneous nuclear ribonucleoprotein A1-like 2 OS=Homo sapiens GN=HNRNPA1L2 PE=2 SV=2 | 34.2 | 1 | 1 | 1 | 2 | 4.50E+03 | 7.10E+04 |
| sp\|P62937\|PPIA_HUMAN | PPIA | Peptidyl-prolyl cis-trans isomerase A OS=Homo sapiens GN=PPIA PE=1 SV=2 | 18 | 1 | 1 | 1 | 1 | 1.50E+04 | 4.60E+04 |
| sp\|Q2VIR3\|IF2GL_HUMAN | EIF2S3L | Putative eukaryotic translation initiation factor 2 subunit 3-like protein OS=Homo sapiens GN=EIF2S3L PE=5 SV=2 | 51.2 | 1 | 1 | 1 | 1 | 5.10E+03 | 2.40E+04 |
| sp\|P06744\|G6PI_HUMAN | GPI | Glucose-6-phosphate isomerase OS=Homo sapiens GN=GPI PE=1 SV=4 | 63.11 | 1 | 1 | 1 | 1 | 1.30E+04 | 1.30E+04 |
| sp\|Q16629\|SRSF7_HUMAN | SRSF7 | Serine/arginine-rich splicing factor 7 OS=Homo sapiens GN=SRSF7 PE=1 SV=1 | 27.35 | 1 | 1 | 1 | 1 | 7.40E+03 | 1.60E+04 |
| sp\|P02545\|LMNA_HUMAN | LMNA | Prelamin-A/C OS=Homo sapiens GN=LMNA PE=1 SV=1 | 74.09 | 1 | 1 | 1 | 1 | 5.30E+03 | 4.80E+04 |
| sp\|P30044\|PRDX5_HUMAN | PRDX5 | Peroxiredoxin-5, mitochondrial OS=Homo sapiens GN=PRDX5 PE=1 SV=4 | 22.07 | 1 | 1 | 1 | 1 | 4.10E+03 | 1.10E+04 |
| tr\|U5LGW0\|U5LGW0_HUMAN | APOL1 | Apolipoprotein L1 (Fragment) OS=Homo sapiens GN=APOL1 PE=4 SV=1 | 32.64 | 1 | 1 | 1 | 1 | 1.80E+04 | 4.50E+04 |
| sp\|O00299\|CLIC1_HUMAN | CLIC1 | Chloride intracellular channel protein 1 OS=Homo sapiens GN=CLIC1 PE=1 SV=4 | 26.91 | 1 | 1 | 0 | 0 | 4.00E+03 | NF |
| sp\|P51659\|DHB4_HUMAN | HSD17B4 | Peroxisomal multifunctional enzyme type 2 OS=Homo sapiens GN=HSD17B4 PE=1 SV=3 | 79.64 | 1 | 1 | 0 | 0 | 3.80E+03 | NF |
| sp\|P69905\|HBA_HUMAN | HBA1 | Hemoglobin subunit alpha OS=Homo sapiens GN=HBA1 PE=1 SV=2 | 15.25 | 1 | 1 | 0 | 0 | 2.00E+03 | NF |
| sp\|P07195\|LDHB_HUMAN | LDHB | L-lactate dehydrogenase B chain OS=Homo sapiens GN=LDHB PE=1 SV=2 | 36.62 | 1 | 1 | 0 | 0 | 8.90E+03 | NF |
| sp\|P31949\|S10AB_HUMAN | S100A11 | Protein S100-A11 OS=Homo sapiens GN=S100A11 PE=1 SV=2 | 11.73 | 1 | 1 | 0 | 0 | 4.70E+03 | NF |
| sp\|P62899\|RL31_HUMAN | RPL31 | 60S ribosomal protein L31 OS=Homo sapiens GN=RPL31 PE=1 SV=1 | 14.45 | 1 | 1 | 0 | 0 | 2.10E+04 | NF |
| sp\|P51116\|FXR2_HUMAN | FXR2 | Fragile X mental retardation syndrome-related protein 2 OS=Homo sapiens GN=FXR2 PE=1 SV=2 | 74.18 | 1 | 1 | 0 | 0 | 2.80E+04 | NF |
| sp\|P62304\|RUXE_HUMAN | SNRPE | Small nuclear ribonucleoprotein E OS=Homo sapiens GN=SNRPE PE=1 SV=1 | 10.8 | 1 | 1 | 0 | 0 | 2.70E+03 | NF |
| sp\|P07437\|TBB5_HUMAN | TUBB | Tubulin beta chain OS=Homo sapiens GN=TUBB PE=1 SV=2 | 49.64 | 1 | 2 | 4 | 34 | 6.00E+04 | 1.90E+07 |
| sp\|Q16790\|CAH9_HUMAN | CA9 | Carbonic anhydrase 9 OS=Homo sapiens GN=CA9 PE=1 SV=2 | 49.67 | 1 | 2 | 7 | 11 | 2.80E+04 | 6.00E+05 |
| sp\|P11142\|HSP7C_HUMAN | HSPA8 | Heat shock cognate 71 kDa protein OS=Homo sapiens GN=HSPA8 PE=1 SV=1 | 70.85 | 1 | 2 | 5 | 8 | 1.80E+04 | 6.60E+05 |
| sp\|P10809\|CH60_HUMAN | HSPD1 | 60 kDa heat shock protein, mitochondrial OS=Homo sapiens GN=HSPD1 PE=1 SV=2 | 61.02 | 2 | 2 | 4 | 5 | 1.90E+04 | 1.10E+05 |
| sp\|P62258\|1433E_HUMAN | YWHAE | 14-3-3 protein epsilon OS=Homo sapiens GN=YWHAE PE=1 SV=1 | 29.16 | 2 | 2 | 3 | 5 | 1.50E+04 | 2.30E+05 |
| sp\|P13639\|EF2_HUMAN | EEF2 | Elongation factor 2 OS=Homo sapiens GN=EEF2 PE=1 SV=4 | 95.28 | 2 | 2 | 3 | 4 | 1.50E+04 | 1.20E+05 |
| sp\|Q96KK5\|H2A1H_HUMAN | HIST1H2AH | Histone H2A type 1-H OS=Homo sapiens GN=HIST1H2AH PE=1 SV=3 | 13.9 | 1 | 2 | 2 | 4 | 1.50E+04 | 1.70E+05 |
| sp\|P23284\|PPIB_HUMAN | PPIB | Peptidyl-prolyl cis-trans isomerase B OS=Homo sapiens GN=PPIB PE=1 SV=2 | 23.73 | 2 | 2 | 2 | 3 | 2.40E+04 | 1.10E+05 |
| sp\|Q8NHW5\|RLA0L_HUMAN | RPLP0P6 | 60S acidic ribosomal protein P0-like OS=Homo sapiens GN=RPLP0P6 PE=5 SV=1 | 34.34 | 2 | 2 | 2 | 3 | 1.80E+04 | 1.60E+05 |
| sp\|P23528\|COF1_HUMAN | CFL1 | Cofilin-1 OS=Homo sapiens GN=CFL1 PE=1 SV=3 | 18.49 | 2 | 2 | 2 | 3 | 1.80E+04 | 4.40E+04 |
| sp\|P62424\|RL7A_HUMAN | RPL7A | 60S ribosomal protein L7a OS=Homo sapiens GN=RPL7A PE=1 SV=2 | 29.98 | 1 | 2 | 2 | 3 | 2.20E+04 | 1.50E+05 |
| sp\|P61978\|HNRPK_HUMAN | HNRNPK | Heterogeneous nuclear ribonucleoprotein K OS=Homo sapiens GN=HNRNPK PE=1 SV=1 | 50.94 | 1 | 2 | 1 | 2 | 1.20E+04 | 4.00E+04 |
| sp\|P00338\|LDHA_HUMAN | LDHA | L-lactate dehydrogenase A chain OS=Homo sapiens GN=LDHA PE=1 SV=2 | 36.67 | 2 | 2 | 1 | 2 | 1.70E+04 | 1.00E+05 |
| sp\|Q9UHX1\|PUF60_HUMAN | PUF60 | Poly(U)-binding-splicing factor PUF60 OS=Homo sapiens GN=PUF60 PE=1 SV=1 | 59.84 | 2 | 2 | 1 | 2 | 1.00E+04 | 2.30E+04 |
| sp\|P13929\|ENOB_HUMAN | ENO3 | Beta-enolase OS=Homo sapiens GN=ENO3 PE=1 SV=5 | 46.96 | 1 | 2 | 1 | 2 | 6.00E+04 | 7.40E+04 |
| sp\|P05386\|RLA1_HUMAN | RPLP1 | 60S acidic ribosomal protein P1 OS=Homo sapiens GN=RPLP1 PE=1 SV=1 | 11.51 | 1 | 2 | 1 | 2 | 4.50E+04 | 3.70E+05 |
| sp\|P50914\|RL14_HUMAN | RPL14 | 60S ribosomal protein L14 OS=Homo sapiens GN=RPL14 PE=1 SV=4 | 23.42 | 1 | 2 | 1 | 2 | 3.30E+04 | 1.60E+05 |
| sp\|P01275\|GLUC_HUMAN | GCG | Glucagon OS=Homo sapiens GN=GCG PE=1 SV=3 | 20.9 | 1 | 2 | 1 | 2 | 4.90E+04 | 4.00E+04 |
| sp\|P52272\|HNRPM_HUMAN | HNRNPM | Heterogeneous nuclear ribonucleoprotein M OS=Homo sapiens GN=HNRNPM PE=1 SV=3 | 77.46 | 1 | 2 | 1 | 2 | 2.20E+04 | 8.40E+04 |
| IgG1_bovine | | heavy chain constant region [Bos taurus] | 35.83 | 2 | 2 | 1 | 1 | 5.70E+04 | 1.30E+06 |
| sp\|P60174\|TPIS_HUMAN | TPI1 | Triosephosphate isomerase OS=Homo sapiens GN=TPI1 PE=1 SV=3 | 30.77 | 2 | 2 | 0 | 0 | 1.70E+04 | NF |
| sp\|P18669\|PGAM1_HUMAN | PGAM1 | Phosphoglycerate mutase 1 OS=Homo sapiens GN=PGAM1 PE=1 SV=2 | 28.79 | 1 | 2 | 0 | 0 | 1.40E+04 | NF |
| sp\|P29401\|TKT_HUMAN | TKT | Transketolase OS=Homo sapiens GN=TKT PE=1 SV=3 | 67.83 | 2 | 2 | 0 | 0 | 7.10E+03 | NF |
| sp\|P68871\|HBB_HUMAN | HBB | Hemoglobin subunit beta OS=Homo sapiens GN=HBB PE=1 SV=2 | 15.99 | 2 | 2 | 0 | 0 | 4.00E+04 | NF |
| sp\|P07478\|TRY2_HUMAN | PRSS2 | Trypsin-2 OS=Homo sapiens GN=PRSS2 PE=1 SV=1 | 26.47 | 1 | 2 | 0 | 0 | 1.20E+06 | NF |
| sp\|P07355\|ANXA2_HUMAN | ANXA2 | Annexin A2 OS=Homo sapiens GN=ANXA2 PE=1 SV=2 | 38.58 | 1 | 2 | 0 | 0 | 2.70E+04 | NF |
| sp\|Q07021\|C1QBP_HUMAN | C1QBP | Complement component 1 Q subcomponent-binding protein, mitochondrial OS=Homo sapiens GN=C1QBP PE=1 SV=1 | 31.34 | 2 | 3 | 9 | 90 | 3.10E+04 | 8.30E+08 |
| sp\|P38646\|GRP75_HUMAN | HSPA9 | Stress-70 protein, mitochondrial OS=Homo sapiens GN=HSPA9 PE=1 SV=2 | 73.63 | 2 | 3 | 8 | 15 | 1.80E+04 | 6.20E+05 |
| sp\|P34931\|HS71L_HUMAN | HSPA1L | Heat shock 70 kDa protein 1-like OS=Homo sapiens GN=HSPA1L PE=1 SV=2 | 70.33 | 3 | 3 | 6 | 9 | 2.70E+04 | 5.00E+05 |
| sp\|P0DMV9\|HS71B_HUMAN | HSPA1B | Heat shock 70 kDa protein 1B OS=Homo sapiens GN=HSPA1B PE=1 SV=1 | 70.01 | 2 | 3 | 4 | 6 | 3.10E+04 | 3.20E+05 |
| sp\|P36578\|RL4_HUMAN | RPL4 | 60S ribosomal protein L4 OS=Homo sapiens GN=RPL4 PE=1 SV=5 | 47.67 | 3 | 3 | 3 | 5 | 3.10E+04 | 3.40E+05 |
| sp\|P05387\|RLA2_HUMAN | RPLP2 | 60S acidic ribosomal protein P2 OS=Homo sapiens GN=RPLP2 PE=1 SV=1 | 11.66 | 2 | 3 | 3 | 4 | 2.70E+04 | 7.90E+06 |
| sp\|P08865\|RSSA_HUMAN | RPSA | 40S ribosomal protein SA OS=Homo sapiens GN=RPSA PE=1 SV=4 | 32.83 | 2 | 3 | 2 | 4 | 2.10E+04 | 1.30E+05 |
| sp\|P00558\|PGK1_HUMAN | PGK1 | Phosphoglycerate kinase 1 OS=Homo sapiens GN=PGK1 PE=1 SV=3 | 44.59 | 2 | 3 | 2 | 3 | 6.30E+04 | 8.60E+04 |
| sp\|P07737\|PROF1_HUMAN | PFN1 | Profilin-1 OS=Homo sapiens GN=PFN1 PE=1 SV=2 | 15.04 | 2 | 3 | 2 | 3 | 4.30E+04 | 7.20E+04 |
| sp\|P32119\|PRDX2_HUMAN | PRDX2 | Peroxiredoxin-2 OS=Homo sapiens GN=PRDX2 PE=1 SV=5 | 21.88 | 2 | 3 | 2 | 3 | 3.30E+04 | 4.50E+04 |
| sp\|Q00839\|HNRPU_HUMAN | HNRNPU | Heterogeneous nuclear ribonucleoprotein U OS=Homo sapiens GN=HNRNPU PE=1 SV=6 | 90.53 | 2 | 3 | 2 | 3 | 1.40E+04 | 8.80E+04 |
| IgG1b_bovine | | immunoglobulin gamma 1 heavy chain constant region [Bos taurus] | 35.88 | 1 | 3 | 1 | 2 | 1.30E+05 | 5.10E+04 |
| sp\|P07900\|HS90A_HUMAN | HSP90AA1 | Heat shock protein HSP 90-alpha OS=Homo sapiens GN=HSP90AA1 PE=1 SV=5 | 84.61 | 2 | 3 | 1 | 1 | 1.20E+04 | 1.10E+04 |
| | HIST1H2BB | Histone H2B type 1-B OS=Homo sapiens GN=HIST1H2BB PE=1 SV=2 | 13.94 | 2 | 3 | 1 | 1 | 2.30E+04 | 1.30E+04 |
| sp\|P04083\|ANXA1_HUMAN | ANXA1 | Annexin A1 OS=Homo sapiens GN=ANXA1 PE=1 SV=2 | 38.69 | 2 | 3 | 0 | 0 | 2.10E+04 | NF |
| sp\|P22626\|ROA2_HUMAN | HNRNPA2B1 | Heterogeneous nuclear ribonucleoproteins A2/B1 OS=Homo sapiens GN=HNRNPA2B1 PE=1 SV=2 | 37.41 | 1 | 3 | 0 | 0 | 3.90E+05 | NF |
| sp\|P19338\|NUCL_HUMAN | NCL | Nucleolin OS=Homo sapiens GN=NCL PE=1 SV=3 | 76.57 | 3 | 4 | 7 | 12 | 8.60E+04 | 5.00E+05 |
| sp\|P11021\|GRP78_HUMAN | HSPA5 | 78 kDa glucose-regulated protein OS=Homo sapiens GN=HSPA5 PE=1 SV=2 | 72.29 | 4 | 5 | 14 | 25 | 5.00E+04 | 1.40E+06 |
| sp\|P06748\|NPM_HUMAN | NPM1 | Nucleophosmin OS=Homo sapiens GN=NPM1 PE=1 SV=2 | 32.55 | 3 | 5 | 5 | 7 | 2.50E+04 | 1.60E+05 |
| sp\|P68104\|EF1A1_HUMAN | EEF1A1 | Elongation factor 1-alpha 1 OS=Homo sapiens GN=EEF1A1 PE=1 SV=1 | 50.11 | 3 | 7 | 9 | 30 | 5.20E+04 | 4.20E+06 |
| sp\|P06733\|ENOA_HUMAN | ENO1 | Alpha-enolase OS=Homo sapiens GN=ENO1 PE=1 SV=2 | 47.14 | 5 | 7 | 3 | 3 | 1.10E+05 | 4.80E+04 |
| sp\|O76031\|CLPX_HUMAN | CLPX | ATP-dependent Clp protease ATP-binding subunit clpX-like, mitochondrial OS=Homo sapiens GN=CLPX PE=1 SV=2 | 69.18 | 5 | 8 | 6 | 7 | 1.10E+05 | 1.90E+05 |
| sp\|Q9BQE3\|TBA1C_HUMAN | TUBA1C | Tubulin alpha-1C chain OS=Homo sapiens GN=TUBA1C PE=1 SV=1 | 49.86 | 4 | 9 | 16 | 97 | 3.20E+05 | 1.60E+08 |
| sp\|P19474\|RO52_HUMAN | TRIM21 | E3 ubiquitin-protein ligase TRIM21 OS=Homo sapiens GN=TRIM21 PE=1 SV=1 | 54.14 | 5 | 9 | 3 | 4 | 8.70E+05 | 5.00E+05 |
| sp\|P68371\|TBB4B_HUMAN | TUBB4B | Tubulin beta-4B chain OS=Homo sapiens GN=TUBB4B PE=1 SV=1 | 49.8 | 6 | 10 | 24 | 124 | 1.40E+05 | 7.20E+07 |
| sp\|P14618\|KPYM_HUMAN | PKM | Pyruvate kinase PKM OS=Homo sapiens GN=PKM PE=1 SV=4 | 57.9 | 7 | 10 | 6 | 8 | 1.10E+05 | 2.40E+05 |
| sp\|P63261\|ACTG_HUMAN | ACTG1 | Actin, cytoplasmic 2 OS=Homo sapiens GN=ACTG1 PE=1 SV=1 | 41.77 | 5 | 10 | 4 | 7 | 2.40E+05 | 3.20E+05 |
| sp\|P04406\|G3P_HUMAN | GAPDH | Glyceraldehyde-3-phosphate dehydrogenase OS=Homo sapiens GN=GAPDH PE=1 SV=3 | 36.03 | 5 | 10 | 4 | 5 | 1.20E+05 | 1.40E+05 |
| sp\|P49327\|FAS_HUMAN | FASN | Fatty acid synthase OS=Homo sapiens GN=FASN PE=1 SV=3 | 273.25 | 10 | 12 | 3 | 3 | 1.10E+05 | 2.60E+04 |
| sp\|Q15208\|STK38_HUMAN | STK38 | Serine/threonine-protein kinase 38 OS=Homo sapiens GN=STK38 PE=1 SV=1 | 54.16 | 14 | 56 | 14 | 49 | 7.00E+06 | 9.90E+06 |
| IGKC_MOUSE | | Ig kappa chain C region OS=Mus musculus PE=1 SV=1 | 11.77 | 10 | 94 | 6 | 32 | 5.40E+07 | 1.40E+07 |
| KV2A7_MOUSE | | Ig kappa chain V-II region 26-10 OS=Mus musculus PE=1 SV=1 | 12.27 | 2 | 215 | 2 | 17 | 1.10E+09 | 2.10E+07 |
| IGH1M_MOUSE | Ighg1 | Ig gamma-1 chain C region, membrane-bound form OS=Mus musculus GN=Ighg1 PE=1 SV=2 | 43.36 | 7 | 310 | 6 | 93 | 1.60E+09 | 3.00E+08 |

(NF: not found)
